# Transcriptome analysis of the effect of AHR on productive and unproductive pathways of *in vitro* megakaryocytopoiesis

**DOI:** 10.1101/2021.05.17.443961

**Authors:** Léa Mallo, Valentin Do Sacramento, Christian Gachet, François Lanza, Henri de la Salle, Catherine Strassel

## Abstract

Human CD34^+^ progenitors can be differentiated *in vitro* into proplatelet-producing megakaryocytes (MKs) within 17 days. During this time, four cell populations emerge, phenotypically defined as CD34^+^CD41^+^ on day 7 (D7) and CD34^+^CD41^+^CD9^-^ on D10 and D14 - qualified as “productive” because they can differentiate into proplatelet-forming cells during the D14-D17 period - and CD34^-^CD41^+^ or CD34^+^CD41^+^CD9^+^ on day 10 - qualified as “unproductive” because they are unable to form proplatelets later. Coculture with mesenchymal stem cells, or the presence of the AHR antagonist SR1, boosts the productive pathway in two ways: firstly, it increases the yield of D10 and D14 CD34^+^CD41^+^CD9^-^ cells and secondly, it greatly increases their ability to generate proplatelets; in contrast, SR1 has no noticeable effect on the unproductive cell types. A transcriptome analysis was performed to decipher the genetic basis of these properties. This work represents the first extensive description of the genetic perturbations which accompany the differentiation of CD34^+^ progenitors into mature MKs at a subpopulation level. It highlights a wide variety of biological changes modulated in a time-dependent manner and allows anyone, according to his/her interests, to focus on specific biological processes accompanying MK differentiation. For example, the modulation of the expression of genes associated with cell proliferation, lipid and cholesterol synthesis, extracellular matrix components, intercellular interacting receptors and MK and platelet functions reflected the chronological development of the productive cells and pointed to unsuspected pathways. Surprisingly, SR1 only affected the gene expression profile of D10 CD34^+^CD41^+^CD9^-^ cells; thus, as compared to these cells and those present on D14, the poorly productive D10 CD34^+^CD41^+^CD9^-^ cells obtained in the absence of SR1 and the two unproductive populations present on D10 displayed an intermediate gene expression pattern. In other words, the ability to generate proplatelets between D10 and D14 appeared to be linked to the capacity of SR1 to delay MK differentiation, meanwhile avoiding intermediate and inappropriate genetic perturbations. Paradoxically, the D14 CD34^+^CD41^+^CD9^-^ cells obtained under SR1^-^ or SR1^+^ conditions were virtually identical, raising the question as to whether their strong differences in terms of proplatelet production, in the absence of SR1 and between D14 and D17, are mediated by miRNAs or by memory post-translational regulatory mechanisms.

## INTRODUCTION

About 10^11^ platelets are produced each day in a human adult; these anucleated cells are generated by the fragmentation, in the circulating blood, of bone marrow megakaryocytes (MKs). This process is controlled by thrombopoietin and other cytokines and by as yet uncharacterized signals from the environment. Some aspects of the process can be reproduced *in vitro* by culturing and differentiating CD34^+^ hematopoietic stem cells or progenitors, which results in the generation of proplatelet-forming cells, the stage just before platelet production. Our laboratory uses a protocol that leads to the development of CD34^+^CD41^+^ cells between D0 and D7, CD34^+^CD41^+^CD9^-^ cells between D7 and D14 and proplatelet-forming MKs between D14 and D17 (**Figure 1**). Under these experimental conditions, three cell populations co-exist on D10, phenotypically defined as CD34^+^CD41^+^CD9^-^, CD34^-^CD41^+^ and CD34^+^CD41^+^CD9^+^, but only the CD34^+^CD41^+^CD9^-^ cells can mature into proplatelet-forming cells. For convenience, the D10 and D14 CD34^+^CD41^+^CD9^-^ cells are qualified as “productive” cells and the differentiation process allowing their development as the “productive pathway”, whereas the two other D10 cell types are said to be “unproductive”. To facilitate reading, the cell populations will be designated as follows: D0 CD34^+^= D0p (for progenitor), D7 CD34^+^CD41^+^= D7MKp (for megakaryocytic precursor), D10 or D14 CD34^+^CD41^+^CD9^-^= D10MKp or D14MKp (for precursor of productive MKs) and D10 CD34^-^CD41^+^ or CD34^+^CD41^+^CD9^+^= D10MKu_34-_ or D10MKu_9+_ (for unproductive cells).

**Figure 1.**
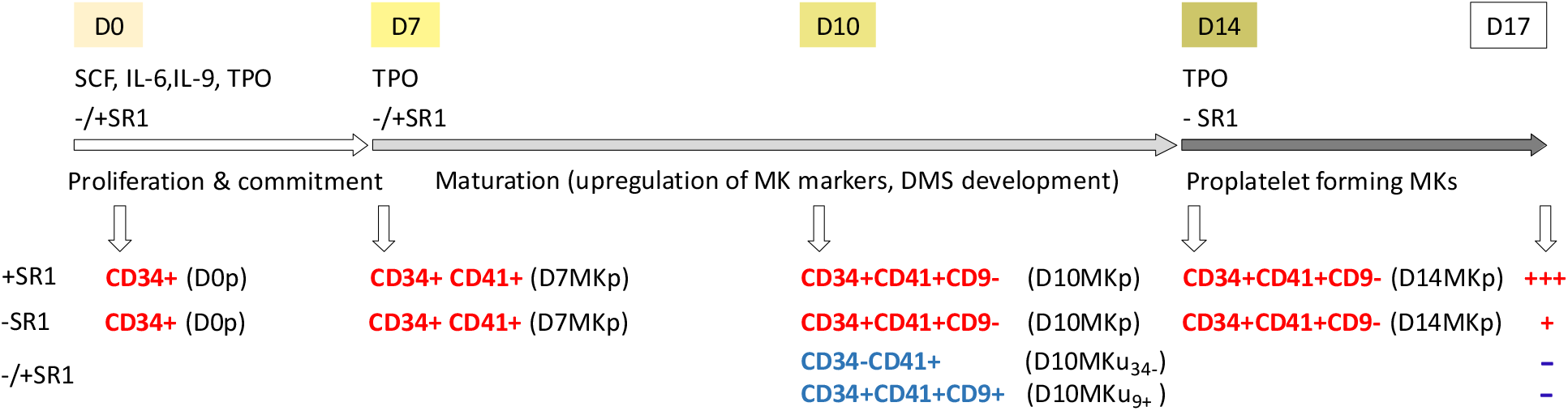
To generate MKs in vitro, CD34^+^ progenitors are cultured for 7 days in StemSpan medium (with SCF, IL-6, IL-9 and TPO) and then for 7 days in a medium containing TPO alone. The D0-D7 period allows the proliferation of MK progenitors and/or the commitment of hematopoietic progenitors. Between D7 and D10 cell proliferation decreases, while between D7 and D14 phenotypical MK maturation progresses. An additional 3-day culture period (D14-D17), not investigated in the present study, permits the generation of proplatelet-producing MKs. The productive pathway (red) can be improved by adding the AHR antagonist SR1 on D0 and D7, which boosts the yield of CD34^+^CD41^+^CD9^-^ productive cells on D10 and their ability to form proplatelets between D14 and D17. On D10, two other megakaryocytic cell populations are also present (D10 CD34^-^CD41^+^ and CD34^+^CD41^+^CD9^+^ cells). If cultured up to D17, these cells remain viable but are unable to differentiate into proplatelet-forming cells (unproductive cells, blue). The respective cell denominations used in the main text are indicated in parentheses.

We previously showed that the introduction, on D0 and D7, of bone marrow mesenchymal cells into this *in vitro* differentiation program boosts not only the generation of productive cells, but also their ability to mature into proplatelet-forming cells. The influence of the microenvironment during MK differentiation can be mimicked by the addition on D0 and D7 of the AHR antagonist SR1. In contrast, neither mesenchymal cells nor SR1 have any observable effect on the unproductive cells (1).

To gain a better understanding of this *in vitro* differentiation process, an RNA-Seq transcriptome analysis was performed on the various cell populations obtained at the different steps of the culture protocol (D0, D7, D10 and D14). The analysis was designed to study five questions. Which features differentially characterize each step of the productive pathway under standard conditions, i.e., in the absence of SR1 (**Table A**, question 1)? How does SR1 modulate the gene expression at each step, with respect to the cells generated in the absence of SR1, or to the experimental time frame (questions 2 and 3, respectively)? How are the differences temporarily coordinated along the productive pathway (question 4)? Finally, which features characterize the two unproductive cell populations present on D10, in terms of the differences and common features between them and with respect to the D10 and D14 productive cells (question 5)? To tackle these problems, we compared the gene expression profiles of selected pairs of conditions or larger sets of conditions (binary analyses or clustering studies, respectively, see Methods and **Table A**). We note that this study was not intended to describe the short-term effects of SR1.

**Table A.** Questions investigated in the transcriptome analyses with the corresponding experimental conditions (- and/or + SR1, cell populations, time points) and methods of analysis (see Methods section for details and Figure 2 for a schematic representation).

|  | Question | SR1 | Analysis type | Conditions compared |
| --- | --- | --- | --- | --- |
| 1 | Analysis of the productive pathway in the absence of SR1 | - | binary | D0p vs D7MKp; D7MKp vs D10MKp; D10MKp vs D14MKp |
| 2 | Effect of SR1 on the productive cells | + or - | binary | D7MKp, SR1- vs SR1+; D10MKp, SR1- vs SR1+; D14MKp, SR1- vs SR1+ |
| 3 | Analysis of the productive pathway in the presence of SR1 | + | binary | D0p vs D7MKp; D7MKp vs D10MKp; D10MKp vs D14MKp |
| 4 | Time dependent gene expression patterns associated with the productive pathway | + or - | clustering | D0p, D7MKp, D10MKp and D14MKp |
| 5 | Gene perturbations associated with the unproductive pathway | + or - | binary and clustering | D10MKu<sub>34-</sub>, D10MKu<sub>9+</sub>, D10MKp and D14MKp |

## MATERIAL AND METHODS

Peripheral blood CD34^+^ progenitors from human adults were purified using Miltenyi technology as previously described and differentiated into MKs using a two-step protocol in StemSpan(tm) Megakaryocyte Expansion Supplement (containing SCF, IL-6, IL-9, TPO) containing human LDL (1) (**Figure 1**), in the absence of SR1, or with SR1 added on D0 and D7. For transcriptome analyses, megakaryocytic cells were isolated by flow cytometry according to their phenotype, more precisely D7 CD34^+^CD41^+^ and D10 and D14 CD41^+^CD41^+^CD9^-^ cells (productive pathway) and D10 CD34^+^CD41^+^CD9^+^ and CD34^-^CD41^+^ cells (unproductive pathway). All cell samples were generated from independent batches of CD34^+^ cells, each prepared from 5 leukocyte-depletion filters and thus representing 20 unrelated donors.

RNAs were prepared from purified D0 cells and from D7 to D14 populations isolated by flow cytometric cell sorting using an RNeasy Plus kit (Qiagen), and then depleted of rRNAs and processed for RNA-Seq library construction using a TruSeq Stranded mRNA Library Prep Kit (Illumina). RNA-Seq data were acquired on an Illumina Hiseq 4000 sequencer. Three independent biological replicates were processed for each condition, i.e., cell subpopulation, day of culture, or with or without SR1. In each RNA-Seq experiment, 16 to 25 10^6^ different non-ribosomal reads were uniquely mapped onto the hg38 human genome assembly. Read counts were normalized across the samples using the median of ratios method (2), then divided by the median value of the length of the transcripts associated with the gene in the ENSEMBL databank and finally log_2_-transformed (in this paper, “normalized expression values”).

To clarify the features characterizing the differences between the cell subsets (questions 1, 2, 3 and 5, Introduction), binary analyses were performed. In practice, selected pairs of conditions (**Table A**) were compared by analyzing their differential gene expression (DGE) using DEseq-2 1.16.1 (Bioconductor). The pValues were adjusted for multiple testing using the Benjamini-Hochberg method (3). The genetic modulations were found to be much more important during the D0-D7 step than in the other ones. Such differences would result in a bias during the normalization process and DEseq2 analysis. Hence two sets of normalized expression values were generated, one including all the data (V1) to study the D0-D7 step, and a second excluding the D0 data (V2) to study the other steps (*Supplemental Table 1 and2*). Expression data and corresponding statistical analyses have been deposited at the Gene Expression Omnibus repository (series record GSE167866).

A first global overview of all binary analyses was obtained using a low stringent threshold of differential expression (|log_2_ FC|> log_2_ 1.5 and ApVals <0.05). To perform a deeper analysis, for each pair of conditions analyzed, the |log_2_ FC| and ApVal thresholds were empirically adapted to generate, when possible, a large but restricted number of DEGs (<2000, see Results for details). Furthermore, for each comparison, to exclude genes not faithfully expressed in all cells, only those displaying a mean normalized expression value of >5 in at least one of the two conditions analyzed were taken into consideration. This latter criterion was supported by analyzing the correlations between the estimated copy numbers of human platelet proteins per platelet and the respective normalized gene expression values in D14 megakaryocytic cells (***Supplemental Figure 1 and Table 3***).

In addition, to reveal time-dependent or cell type-dependent differential expression patterns, larger combinations of conditions were analyzed together (questions 3 and 4): (i) D0p, D7MKp, D10MKp and D14MKp cells - to analyze the time-dependent variations in gene expression along the productive pathway - and (ii) D10MKp, D14MKp, D10MKu_34-_ and D10MKu_CD9+_ cells - to compare productive and unproductive cells. As for binary analyses, only the genes displaying a mean normalized expression of >5 for at least one of the conditions analyzed were taken into account. The normalized expression values were compared by ANOVA followed by a Benjamini-Hochberg correction. Genes exhibiting an expression pattern with a false discovery rate (FDR) of <0.001 were selected and clustered according to their cell-dependent gene expression patterns using a fuzzy C-mean algorithm (4). Briefly, for this clustering, the expression values were transformed: for each gene and sample, the normalized expression was centered on the mean of the expression across all conditions, after which the standard deviation of these centered expression values was normalized to 1. The resulting transformed data were used for fuzzy C-mean clustering (softwares kindly provided by D. Dembelé, IGBMC).

These analyses defined lists of dDEGs and uDEGs or DEGs associated with cell- and/or time-dependent expression patterns, which were then further analyzed to reveal correlated enriched biological attributes. Panther Gene Statistical Enrichment Analyses (GSEAs) of cellular compartments (GO CC) or biological processes (GO BP) and of Reactome pathways were implemented, using a Fisher exact test and imposing an FDR of <0.05 and a positive enrichment of >1.3 (http://pantherdb.org/) (5).

The DEG lists were also analyzed with a ChEA3 algorithm to unravel co-regulation patterns associated with transcription factors or regulators (TFRs) (6). In this paper, the mean ranks and scores were considered. Since this bioinformatics tool sometimes points to TFRs which are not expressed in the cells analyzed, only TFRs exhibiting a mean expression value of >5 for at least one of the two conditions analyzed were taken into account. It should nevertheless be acknowledged that this threshold does not necessarily imply that these TFRs are actually expressed at biological levels in some or all of the cells.

Finally, Venn diagram analyses of multiple DEG lists were performed (http://www.interactivenn.net/).

## RESULTS

### Overview of the gene expression perturbations occurring during the differentiation of CD34^+^ cells into MKs, in the absence or presence of SR1

To gain an overview of the variations in gene expression occurring in the different cell populations during the *in vitro* differentiation of CD34^+^ progenitors into MKs, in the absence or presence of SR1 (SR1^-^or SR1^+^), we first compared biologically significant pairs of conditions (binary analyses), using low fold change (FC) and adjusted pValue (ApVal) thresholds (|log_2_ FC| > log_2_ 1.5, ApVal <0.05). The differential analysis data are available in the ***Supplemental Tables 1 and 2***.

Under these criteria and in the absence of SR1, more than 50% of the genes analyzed were differentially expressed during the D0-D7 period (14009 genes). Between D7 and D10 and between D10 and D14, the differentiation of MKp cells was accompanied by a reduced but still high percentage of DEGs (20 and 24% of the genes analyzed, or 4005 and 5849 genes, arrows b and c, respectively).

Unexpectedly, when comparing SR1^-^ and SR1^+^ conditions, the transcriptomes of D7MKp cells, D10 unproductive subsets (D10MKu_34-_ or D10MKu_9+_ cells) and D14MKp cells were similar (arrows d0 and d2); the expression of only 598 genes was modulated by SR1 in D10MKp cells (arrow d1). In agreement with these observations and the analysis of SR1^-^ conditions, large numbers of DEGs were associated with the SR1^+^ D0-D7 and D10-D14 steps (13742 and 6414 genes, arrows a’ and c’, respectively). In sharp contrast with SR1^-^ conditions, only 148 genes were differentially expressed between D7MKp and D10MKp cells in the presence of SR1 (arrow b’). The numbers of DEGs associated with SR1^-^ vs SR1^+^ D10MKp cells (arrow d1) and with SR1^+^ D7MKp vs D10MKp cells (arrow b’) might appear paradoxical when considering the other numbers of DEGs during the D7-D14 period (arrows b, c and c’), but they reflect the fact that among the 4005 genes differently expressed between SR1^-^ D7MKp and D10MKp cells (arrow b), 3884 displayed an |FC_SR1+_|<|FC_SR1-_|, whereas the corresponding ApVals had an inverse relationship for 3871 of them, resulting in a reduced contrast between D10MKp cells generated under SR1^-^ and SR1^+^ conditions (arrow d1).

Thus, in terms of period-associated genetic modulations, SR1 appeared to affect specifically D10MKp cells; we do not exclude that SR1 could also have short term effects on the cells, but this study was not intended to investigate such effects. In a general manner, the variations in gene expression in D10MKp cells were less under SR1^+^ than under SR1^-^ conditions. Remarkably, the transcriptomes of D14MKp cells generated in the presence or absence of SR1 were virtually the same. This observation is paradoxical, since in the absence of SR1, the abilities of these two D14 cell populations to mature into proplatelet-forming MKs are very different.

### Detailed biological interpretations of the binary analyses

The binary comparisons were further analyzed to decipher the biological features associated with the D0 to D14 productive pathway, firstly under SR1^-^ conditions (**Table A**, question 1), to elucidate the effect of SR1 on D10 and D14 productive cells or their D7 precursors (question 2), and then to clarify how SR1 influences time-dependent MK differentiation (question 3). These investigations were complemented with a global analysis of MK differentiation along the D0 to D14 *in vitro* pathway (question 4). Finally, we attempted to unravel the differences between D10 unproductive and D10 or D14 productive cells (question 5).

To clarify the biological significance of the DGE profiles associated with the different pairs of conditions, the numbers of DEGs were adapted by choosing more or less stringent FC and/or ApVal thresholds. In addition, as justified in the Methods section, only genes displaying a mean expression of >5 in at least one of the two conditions were analyzed. The upregulated and downregulated genes (uDEGs and dDEGs) were independently investigated using two algorithms. Firstly, gene set enrichment analyses (GSEAs) of GO BP, GO CC and Reactome pathways (hereafter designated as “attributes”) were performed (**section A**) and secondly, correlations between the condition-specific dDEG or uDEG lists and transcription factors or regulators (TFRs) were analyzed using ChEA3 (**section B**).

### A- Gene set enrichment analyses

Concerning the binary analyses, **Figures 3a** and **3b** indicate the thresholds used to define the DEGs, the corresponding numbers of uDEGs and dDEGs and the main findings derived from GSEA of the productive and the unproductive pathways, as described in detail in the following paragraphs. All data are available in the *Supplemental Tables 4* (DEG lists) and Table 5-28.xlsx (GSEAs).

### A1 - Three biologically distinct steps are associated with the productive differentiation pathway in the absence of SR1

This section refers to the arrows a, b and c in Figures 2 and 3a.

**Figure 2.**
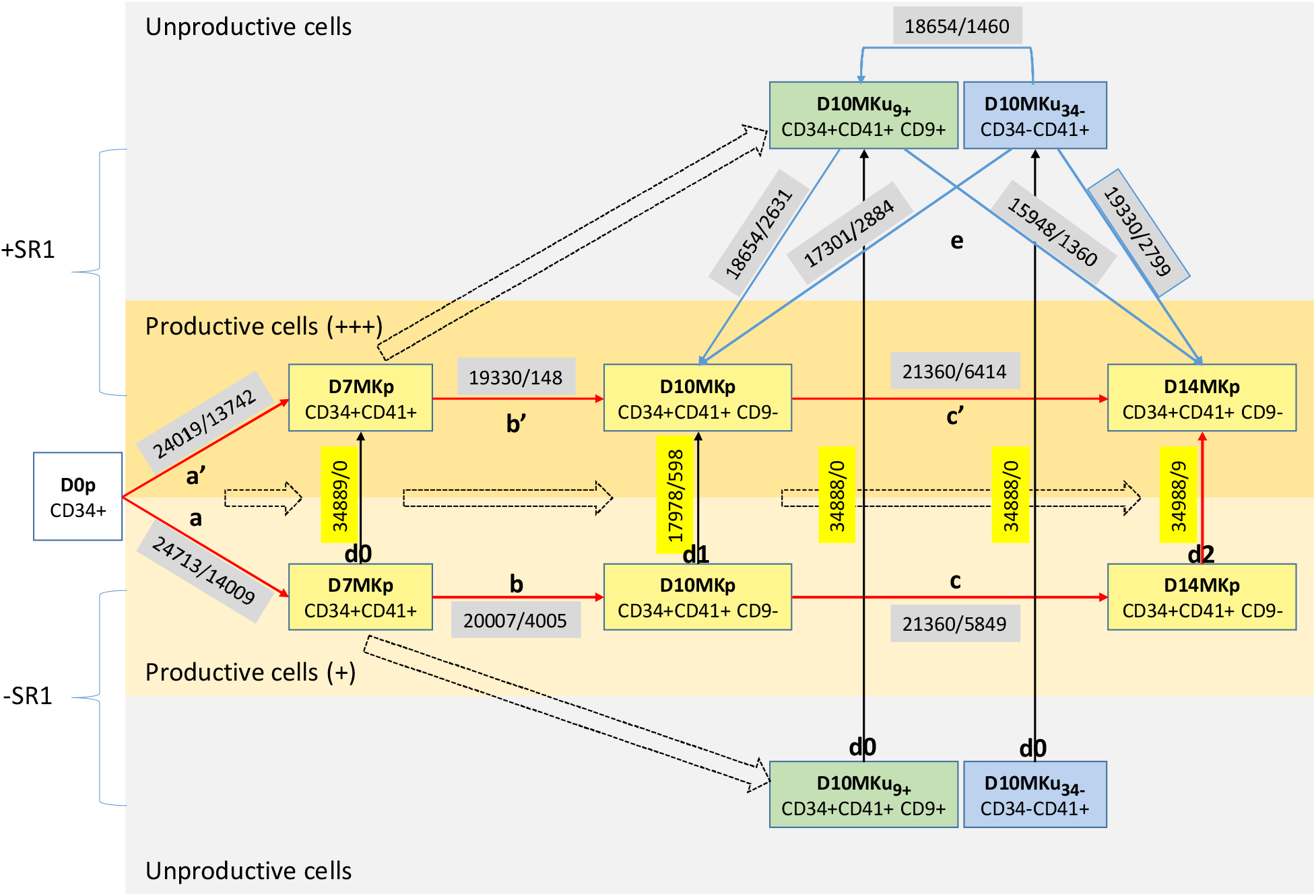
CD34^+^ progenitors (D0p) were differentiated into MKs according the schema depicted in Figure 1, in the presence (+SR1, top) or absence of SR1 (-SR1, bottom). The dashed arrows show the chronological dependence between the cell types. The indicated cell populations were isolated by flow cytometry according to their phenotypes on D7, D10 and D14 and then processed for RNA-seq analysis. Differential gene expression was evaluated using DEseq2 to compare pairs of conditions (cell types and culture conditions). For each pair of conditions analyzed (joined with an arrow), two numbers are provided: the number of DEseq2-analyzed genes (i.e., after having rejected the DEseq2-calculated outliers) and the numbers of DEGs (|log_2_ FC|> log_2_ 1.5, ApVal <0.05, no threshold for the mean expression values). The yellow boxes highlight these numbers for the direct comparisons between SR1^-^ and SR1^+^ conditions (black arrows). The letter-labeled arrows refer to the binary analyses detailed below. Arrows a, b and c on the one hand and a’, b’ and c’ on the other hand are relative to the productive pathway under SR1^-^ (§A1a-c) and SR1^+^ (§A2a-c) conditions, respectively; arrow d corresponds to the comparison of D10MKp cells generated under SR1^-^ and SR1^+^ conditions (§A2c-e). The time-dependent differential expression patterns between D0 and D14 MKp cells were analyzed in §A2f. Blue arrows e indicate the analyses performed to clarify the differences between the two D10 unproductive cell subsets and D10 or D14 productive cells, all generated in the presence of SR1 (§A3a, b). The differences between D10 unproductive and D10 or D14 productive cells were analyzed in §A3c.

**Figure 3a.**
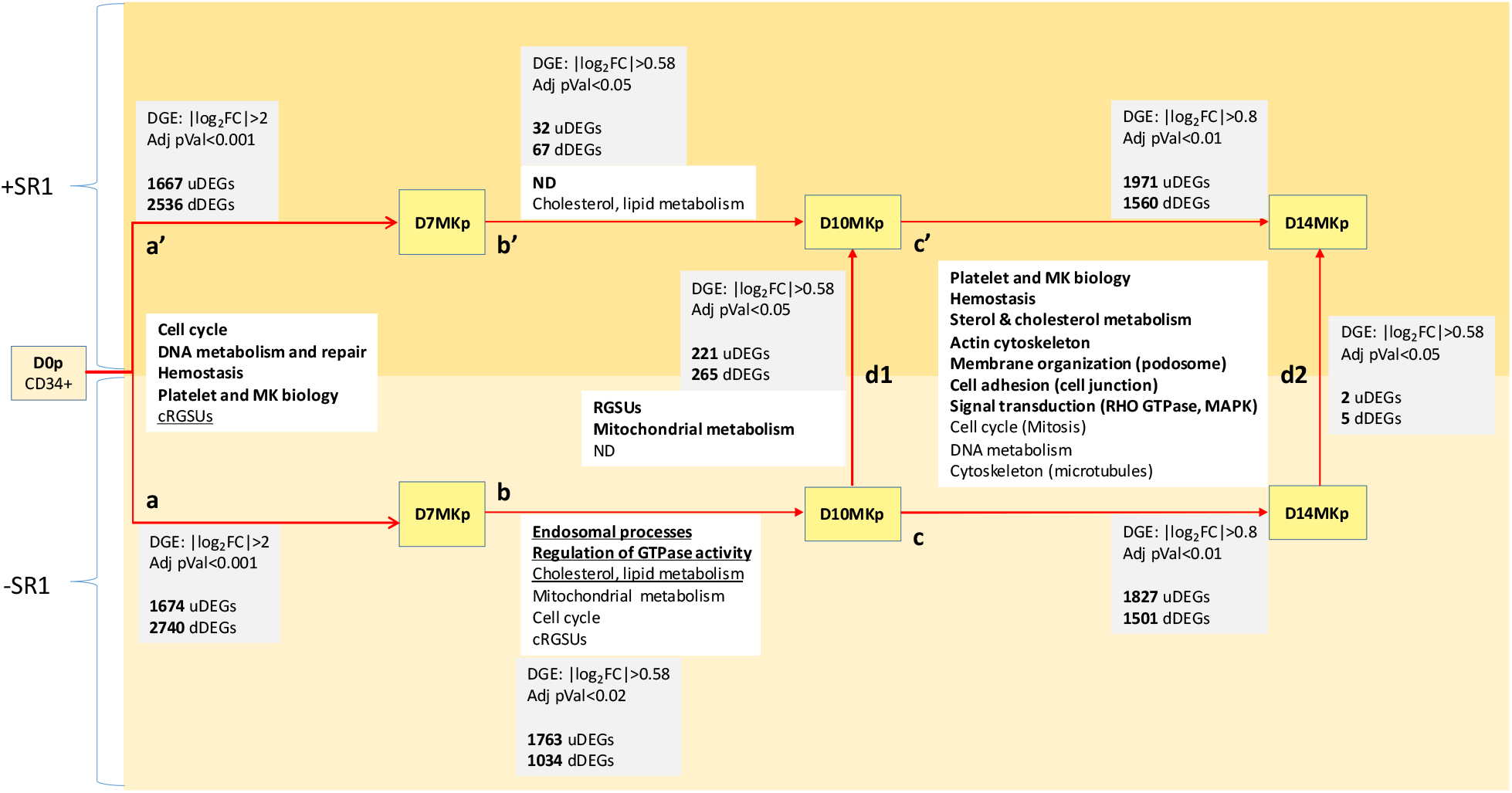
Analysis of the productive pathway. The transcriptomes of pairs of conditions were analyzed using DEseq2. The lists of positively and negatively modulated DEGs (calculated between the origin and target conditions represented by the arrows) were independently probed with Panther gene enrichment analysis tools; the FC and ApVal thresholds used are indicated. Only genes exhibiting a mean expression of >5 for at least one condition were taken into account. In this figure, the enriched attributes (detailed in the inserts) are designated in general terms; cRGSUs, cytoplasmic ribosomal subunit-biased attributes; ND, no enriched entities detected. Bold and normal characters correspond to attributes associated with uDEGs and dDEGs, respectively. The arrow code is the same as in Figure 2.

**Figure 3b.**
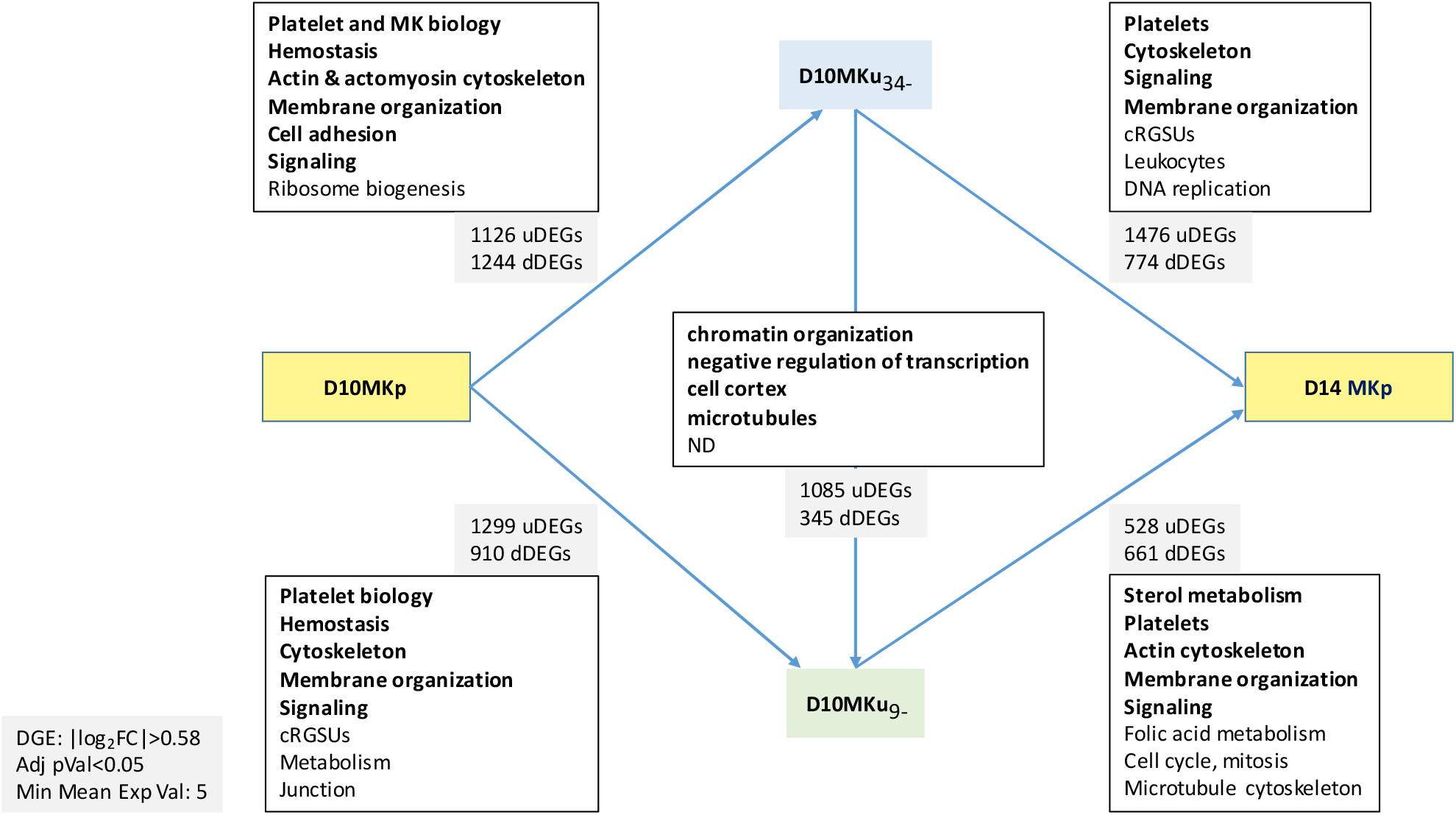
Comparison of the unproductive D10MKu_34-_ and D10MKu_9+_ cells with the productive D10MKp and D14MKp cells generated in the presence of SR1 (corresponding to arrows e in Figure 2). The lists of positively and negatively modulated DEGs (calculated between the origin and target conditions represented by the arrows) were independently analyzed with Panther gene enrichment analysis tools; |log_2_ FC| > log_2_ 1.5 and ApVal <0.05. In this figure, the enriched attributes (detailed in the inserts) are designated in general terms; bold and normal characters correspond to attributes associated with uDEGs and dDEGs, respectively.

#### A1a - The D0-D7 expansion and commitment phase (Supplemental Tables 6 and 7)

Rather stringent thresholds (|Log_2_ FC| > 2 and ApVal <0.001) were used to compare D0 CD34^+^ cells and D7 CD41^+^ cells, allowing us to restrict the selection to 2740 dDEGs and 1,674 uDEGs. The GSEA of the uDEGs highlighted many GO BP (407) and GO CC (100) terms and Reactome pathways (197) related to cell cycle, DNA replication and DNA repair-associated processes and in addition, GO BP terms related to metabolic processes, including the cholesterol biosynthesis pathway. All these attributes were somewhat expected since quiescent progenitor cells were compared to proliferating cells cultured *in vitro*. Moreover, attributes linked to platelets (113 genes) and their response to wounding (95 genes), totalizing 142 genes, were identified, reflecting the megakaryocytic commitment of the D7 CD41^+^ cells.

Analysis of the dDEG list pointed to translation-associated processes, which could reflect the relatively high number of large or small cytoplasmic ribosomal subunit genes (cRSUGs) among the dDEGs. A systematic analysis revealed that 81 of the 100 cRSUGs displayed a log_2_ FC of <-1 (ApVal <0.001), whereas the expression of mitochondrial RSUGs displayed an inverse tendency - among 78 of them, 22 were upregulated and 7 downregulated (log_2_ FC > 1 ApVal <0.001) (**Figure 4**). This important bias of the cRSUGs strongly suggests that the ribosomal architecture is dramatically modified during the early step of the culture of CD34^+^ progenitors and highlights the importance of the regulation of translation, or alternatively of translation-independent functions of these proteins, during megakaryocytic commitment (see **Discussion**). The upregulation of mitochondrial RSUGs could be related to the increased metabolism of the cultured cells.

**Figure 4.**
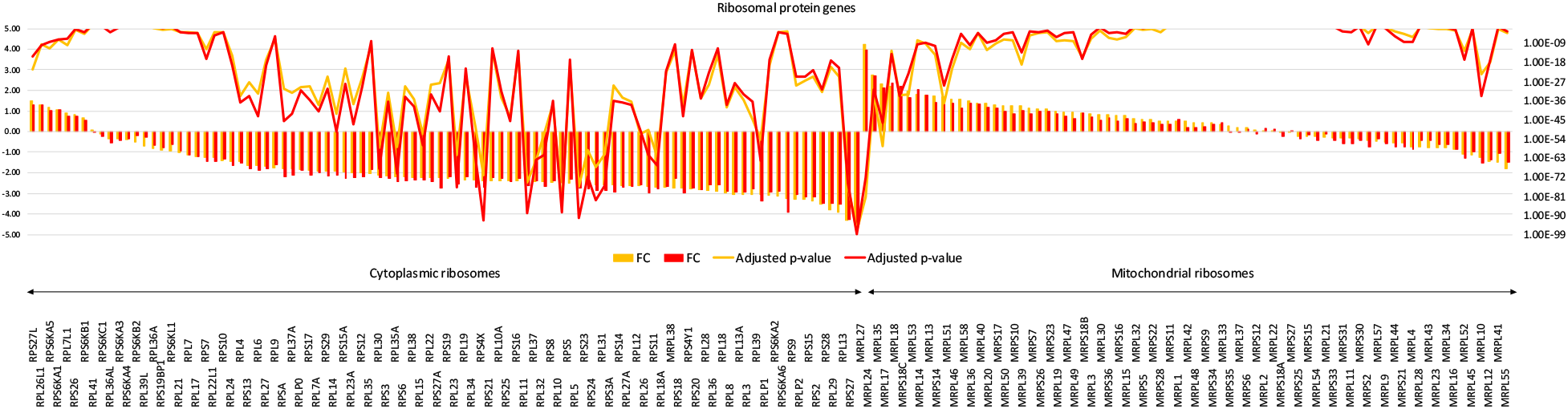
FCs of cytoplasmic and mitochondrial RSUGs (left cytosolic, right mitochondrial; orange and red bars, SR1^-^ and SR1^+^ conditions, respectively; FC, log_2_ scale, left axis). Genes were plotted according to their decreasing |log_2_ FC| under SR1^-^ conditions. The corresponding ApVals are joined by broken lines (log_10_ scale, right axis; only values <0.01 are represented).

Many other GO BP terms were related to immunological cells, which might reflect the loss of these contaminating cells during the 7 days of culture. This loss could be documented by analyzing the dDEG list with ENRICHR (https://amp.pharm.mssm.edu/Enrichr) (7), which indicated a decreased expression on D7 of genes mapped in the Human Gene Atlas and expressed by CD14^+^ monocytes, CD19^+^ B cells, CD33^+^ myeloid cells and CD4^+^ T cells (***Supplemental Table A***).

#### A1b - The D7-D10 step, an intermediate period (Supplemental Tables 10 and 11)

Lower stringency parameters were set (Log_2_ FC>log_2_1.5 and ApVal<0.02), resulting in 1,763 uDEGs and 1,034 dDEGs. Although their number is relatively high, the GSEA of the uDEGs pointed to relatively few enriched attributes (18 GO BP and 12 GO CC terms, no Reactome pathways). The most enriched GO BP terms were related to endosomal processes, autophagy, regulation of GTPase activity and, negative regulation of transcription. Of note, 49 uDEGs belonged to MK-or platelet-related attributes displaying non-significant Fold Enrichment Scores (FESs) (Reactome and GO BP, FDR=1) (***supplemental Tables B and C***). Thus, during this period, in the absence of SR1, MK differentiation-associated processes appear to be marginal, whereas the biological relevance of the enriched attributes needs to be clarified.

In contrast, GSEA of the 1,034 D7-D10 dDEGs revealed larger numbers of enriched attributes (319 GO BP and 146 GO CC terms and 294 reactome pathways). Many of these attributes were associated with metabolic processes, in particular cholesterol, steroid, alcohol and lipid biosynthetic functions, and with mitochondrial respiration pathways, especially energy production and mitotic processes. Since the generation of mature MKs requires energy-demanding membrane biosynthesis and endomitosis, the downregulation of these functions was not anticipated.

#### A1c - The D10-D14 maturation phase: enrichment of upregulated MK-associated functions (Supplemental Tables 12 and 13)

For this analysis, we chose more stringent thresholds (|log_2_ FC| > 0.8 and ApVal <0.01), which resulted in 1501 dDEGs and 1827 uDEGs. GSEA of the uDEG list highlighted many enriched attributes (626 GO BP and 149 GO CC terms and 110 Reactome pathways), several of these being highly relevant in terms of megakaryocyte biology and platelet generation, including platelet functions, metabolism, cytoskeleton, plasma membrane dynamics, signaling and cell adhesion, and intercellular processes. Notably, genes involved in cholesterol biosynthesis were upregulated during this period. Analysis of the dDEGs identified 355 GO BP and 76 GO CC terms and 127 Reactome pathways, mostly related to cell cycle and DNA repair processes. Many of these attributes were also found under SR1^+^ condition and will be mentioned again below (**section A2c**).

### A2- Two biologically distinctive steps are associated to the productive differentiation pathway in the presence of SR1

#### A2a-Paradoxically, the transcriptomes of D7MKp and D14MKp cells are not significantly modulated by SR1

The transcriptomes of D7MKp cells under SR1^-^ and SR1^+^ conditions were practically the same; even when applying low stringency thresholds (|log_2_ FC| > log_2_ 1.5, ApVal <0.05) no DEGs were identified. Thus, the numbers of uDEGs and dDEGs appearing between D0 and D7 were comparable under SR1^+^ and SR1^-^ conditions (|log_2_ FC| > 2 and ApVal <0.001; 1674 vs 1667 uDEGs and 2740 vs 2536 dDEGs, respectively, **Figure 3a**) and the vast majority of them were shared between the two conditions (1551 uDEGs and 2393 dDEGs).

Only 7 genes had a mean expression value >5 and could be considered to be differentially expressed between the SR1^-^ and SR1^+^ D14 CD34^+^CD41^+^CD9^-^ cells (|Log_2_FC|>Log_2_1.5, ApVal <0.05), 5 being less expressed in the SR1^+^ condition. Among the later, ARHGAP24 (Log_2_FC=-1.42), a Rac-specific GTPase activating protein, and MICAL2 (Log_2_FC -2.46), a regulator of stress fiber formation, could be relevant to processes associated to MK maturation, but the statistical significance of the differential expression was borderline (ApVal=0.046 and 0.041, respectively).

This absence of a major effect of SR1 on the transcriptomes of D7MKp and D14MKp cells was not anticipated for two reasons. Firstly, addition of SR1 on D0 and D7 promotes the generation of precursors of productive MKs and secondly, D14 cells generated in the presence of SR1 between D0 and D10 display a very different proplatelet-generation potential in the following SR1-independent period.

#### A2b - SR1 added on D7 preserves D7-D10 modulations of lipid and cholesterol metabolism and prevents MKp cells from other other intermediate gene perturbations (supplemental data Table 14)

Whereas 2397 genes were differently expressed between D7 and D10 under SR1^-^ conditions (|log_2_ FC| > log_2_ 1.5 and ApVal <0.02), only 105 genes were modulated under SR1^+^ conditions (|log_2_ FC| > log_2_ 1.5 and ApVal <0.05). Six of these were dDEGs encoding biologically irrelevant variable or constant regions of immunoglobulins and were thus excluded from the analysis. Among the 99 remaining genes, 32 were upregulated and 67 downregulated, and a Venn representation showed that 24 and 33 of them, respectively, were significantly modulated in the same direction as under SR1^-^ conditions (***Supplemental Figure 2***).

The 32 uDEGs were not significantly enriched in GSEA attributes, which could be expected in view of their low number. Although the list of dDEGs was also small, 67 genes, their GSEA pointed to GO BP and Reactome pathways associated with lipid, sterol and cholesterol metabolism, which were also identified by analysis of the 33 genes downregulated under both SR1^-^ and SR1^+^ conditions. In contrast, during this period, the genes involved in mitochondrial metabolism and downregulated under SR1^-^ conditions were not differentially expressed in the presence of SR1; accordingly, among the 67 dDEGs, only CYP27A1 is predicted to encode a mitochondrial protein (http://mitominer.mrc-mbu.cam.ac.uk/release-4.0).

Thus, SR1 appeared to limit the gene expression perturbations occurring during the D7 to D10 step. During this period, as under SR1^-^ conditions, genes participating in cholesterol and lipid metabolism were significantly downregulated. One exception to this tendency was the overexpression of CYP51A1 under SR1^+^ conditions, which might have major biological consequences (see **Discussion**).

#### A2c-SR1 reduces gene expression perturbations during the D7-D10 intermediate period

A Venn diagram analysis revealed that 185 dDEGs and 499 uDEGs were differently expressed in MKp cells between D7 and D10 in the absence of SR1, on the one hand, and between D10 and D14 under SR1^-^ or SR1^+^ conditions, on the other hand (***Supplemental Figure 3a***, *upper row*). GSEA failed to reveal any enriched attributes associated with the 499 upregulated genes, while cRSUG-biased attributes and others related to metabolism or cell cycle were associated with the 193 downregulated genes. Only 5% of the genes were regulated in opposite directions (***Supplemental Figure 3a***, *bottom row*). Thus, most of the genes expressed differentially between SR1^-^ D7MKp and D10MKp cells did not display consistent modulations between D7 and D10 or between D10 and D14 under SR1^+^ conditions, suggesting that most of the gene perturbations associated with the SR1^-^ D7-D10 step were dispensable for MK differentiation and could therefore be detrimental, as compared to SR1^+^ conditions.

#### A2d - GSEA attributes associated with D10-D14 genetic modulations in MKp cells are mostly conserved under SR1^-^ and SR1^+^ conditions (Supplemental Tables 15, 16, 17 and 18)

In the presence of SR1, the number of genes differentially expressed during the D10-D14 step was 25% higher than in its absence. More than 70% of the dDEGs (1079) or uDEGs (1,433) (|log_2_ FC| > 0.8 and ApVal <0.01) displayed qualitative and significant down- or upregulation under both SR1^-^ and SR1^+^ conditions (***Supplemental Figure 3a***, *upper row*). Indeed, GSEA of the DEGs relative to the D10-D14 step for MKp cells, in the presence or absence of SR1, pointed to a majority of common attributes (***Supplemental Table 12 vs 15 and Table 13 vs 16***). Relatively large numbers of attributes associated with uDEGs highlighted various relevant biological mechanisms involving metabolism (e.g., cholesterol biosynthesis), cytoskeleton (actin, myosin and tubulin), cellular transport, intercellular contacts and interactions with the extracellular matrix (integrin, plexin and semaphorins), signaling (Rho GTPase and MAPK), membrane structure (endomembrane and plasma membrane organization) and platelets. The common dDEG attributes were strongly associated with cell cycle or mitotic processes, purine metabolism or DNA repair.

Since SR1^-^ and SR1^+^ D14MKp cells displayed the same gene expression profile, the condition-specific attributes related to the SR1^-^ and SR1^+^ D10-D14 periods reflected the differences between the respective D10MKp cells. Thus, in D10MKp cells, 221 and 265 genes were respectively more or less expressed under SR1^+^ than under SR1^-^ conditions (arrow d, **Figure 3a**) (|log_2_ FC| > log_2_ 1.5 and ApVal <0.05; ***Supplemental Tables 17 and 18***). The majority of the former genes (141/221) were downregulated between D10 and D14 (***Supplemental Figure 4a***), whereas the vast majority of the later ones (232/265 genes) were upregulated (***Supplemental Figure 4b***). Hence SR1 delayed the modulation of the expression of most of the genes differently expressed between SR1^-^ and SR1^+^ D10MKp cells.

GSEA of the 221-gene set pointed to gene lists containing relatively high ratios of genes encoding proteasome subunits A or B (PSMAs or PSMBs) or cRSUGs. GSEA of the 265-gene set yielded only a poorly meaningful GO CC “nucleoplasm” term (1.8-fold enrichment) and no significantly enriched GO BP terms or Reactome pathways. These latter GSEAs were thus poorly informative and a gene-by-gene approach had to be employed

#### A2e - Gene-by-gene analysis of D10MKp cells points to SR1- and gene-dependent expression patterns relevant to mature MK functions

Further analysis of the genes expressed differentially between SR1^-^ and SR1^+^ D10MKp cells focused on those encoding proteins, i.e., 165 and 184 genes respectively more or less expressed under SR1^+^ than under SR1^-^ conditions. Moreover, the focus was narrowed to genes also differentially expressed between D10 and D14 under SR1^+^ conditions (|log_2_ FC| > 0.8, ApVal <0.01). Among these genes, respectively 66 and 131 were differently expressed during the D10-D14 step in the presence of SR1 (65 dDEGs and 1 uDEG, and 131 uDEGs) (***Supplemental Table 29***). By definition, these genes correspond to those for which the expression was most strongly affected by SR1 on D10, before the cells reached the SR1-independent D14 phase. Since intuitively they might play important roles in the physiology of MK maturation, their reported properties were checked for their relevance to MK differentiation.

Among the 65 dDEGs, 15 encoded small or large ribosomal proteins. Several of these (RPS3, RPS7, RPS6, RPS10, RPL24, RPS30 and RPS38) displayed GO annotations reflecting diverse biological processes and/or involved cellular components not directly related to translation, an indication that complex biological functions remain to be clarified in the context of the later steps of *in vitro* megakaryocytopoiesis. Among the 12 dDEGs differently expressed under SR1^+^ and SR1^-^ conditions, PCLAF (a regulator of centrosome number), MDFIC (stabilizing beta catenin), SNRPE (controlling RNA splicing) and TPD52 (a positive regulator of lipid storage and cell proliferation) might participate in the control of MK differentiation. The only gene in this group to be upregulated between D10 and D14 encodes an atypical chemokine, CKLF (***Supplemental Figure 5***). Although this latter protein has been the subject of a limited number of investigations, its upregulation during the later stages of MK differentiation is puzzling and the biological significance of its expression in these cells with respect to crosstalk between hematopoiesis, the megakaryocytic bone marrow niche and/or platelet biology needs to be clarified.

The 131 uDEGs represented a wide variety of biological processes. Several appeared to regulate diverse cell functions relevant to MK biology, at the levels of (i) transcription - MZF1, LYL1 and ZFPM1, three transcription factors known to participate in megakaryocytopoiesis (8) -, (ii) cytoskeletal organization - PLEC, involved in the regulation of cytoskeletal dynamics; TPGS1, a subunit of the tubulin polyglutaminase complex found to regulate the cellular localization of this complex in neural cells (9), which illustrates the importance of post-translational modifications of tubulin in megakaryocytopoiesis (10) -, (iii) membrane receptors potentially participating in DMS organization, or in interactions with bone marrow cells such as endothelial and myeloid cells - PLXN3B could interact with endogenously expressed SEMA4B and SEMA4D, or with the endothelial semaphorins 4A and 5A (11); ICAM5 interacts with integrins and vitronectin – and (iv) signaling processes - PRR7 negatively regulates the exosomal secretion of Wnt molecules; ZNF467 promotes the transactivation of STAT3 (12); TNK2 catalyzes the phosphorylation of STAT1 and STAT3 and might complement or synergize with ZNF467; RADIL, a Gβγ subunit of heterotrimeric G proteins, regulates cell-matrix adhesion by triggering Rap1a-dependent inside-out signals and integrin activation (***Supplemental Figure 5***).

Thus, many of the genes of D10MKp cells differently expressed under SR1^+^ and SR1^-^ conditions and overexpressed between D10 and D14 point to promising new fields of investigation relevant to major functions activated during *in vitro* and/or *in vivo* megakaryocytopoiesis.

#### A2f - Analysis of time- and cell-dependent biological processes of the thrombocytogenic pathway: findings complementary to the binary analyses (Supplemental Tables_30-37)

In this section, we looked for time-dependent patterns of differential gene expression in productive cells, between D0 and D14 and under SR1^-^ or SR1^+^ conditions. This method takes into account the variations in gene expression between all conditions (cells and time points). As compared to binary analyses, it establishes gene lists based on different thresholds of significance (ANOVA of the triplicate expression values and FDR thresholds) and therefore is complementary - in particular, the gene lists may partially intersect. The normalized expression values were analyzed by ANOVA and then by fuzzy C-means clustering, using six groups of expression patterns (**Figure 5**). Fuzzy C-means clustering allowed us to assign half of the DEGs to cell type-dependent differential gene expression profiles, after which each group was analyzed by GSEA.

**Figure 5.**
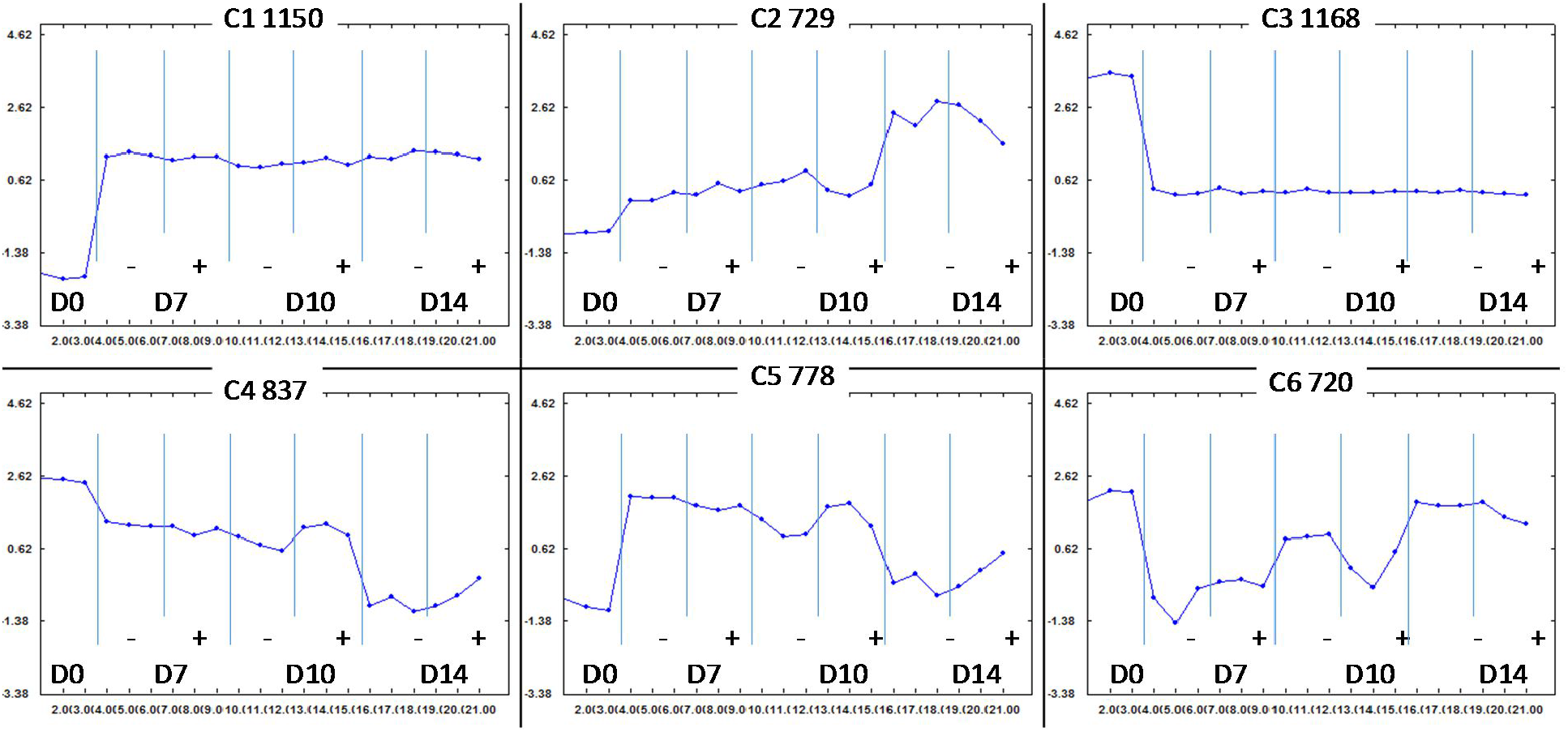
D0-D14 expression patterns of CD34^+^ progenitors (D0) and cell populations associated with the thrombocytogenic pathway (D7, D10 and D14MKp cells, treated (+) or not (-) with SR1). The normalized expression values were analyzed by ANOVA (see Methods) and gene expression profiles with an FDR of <0.001 (10722 genes) were selected for fuzzy C-means clustering after choosing 6 clusters.

Genes belonging to clusters C3 and C1 displayed a predominant down- or upregulation respectively between D0 and D7, as compared to the subsequent D7-D14 periods. As expected, these clusters strongly intersected with the D0-D7 dDEG and uDEG lists (701 and 768 genes, respectively), thereby figuring in the same GSEA categories. In particular, cluster 1 was enriched in genes involved in phospholipid metabolism, Golgi compartment organization, cell division-associated processes, microtubule organization, MK differentiation and platelet responses; these genes are thus coordinated in their MK commitment.

Cluster 2 included 729 genes progressively upregulated from D0 to D14, mostly already between D0 and D7, but above all from D10 to D14. GSEA identified 26 genes participating in the “Platelet activation, signaling and aggregation” reactome pathway, reflecting megakaryocytic commitment between D0 and D7 and overexpression during D10-D14 MK maturation. The genes of this group were also involved in the organization of cellular structures important in MK maturation: (i) the trans-Golgi network, which has previously been shown to expand and to lie in the proximity of the demarcation membrane system (DMS) of mature MKs (13)—, (ii) secretory vesicles, to which platelet granules are related, and (iii) membrane organization (podosomes, ruffles, the leading edge of cells), which highlights the role of plasma membrane dynamics in mature MKs and is in agreement with the importance of podosomes in intravasation (14).

Genes in cluster 5 were upregulated from D0 to D10 and then progressively downregulated between D10 and D14 in the SR1^-^ condition, whereas cluster 6 displayed an inverse expression profile. These two clusters were also characterized by SR1-controlled delayed genetic modulations on D10. The former one was enriched in genes participating in cell division and DNA repair-associated processes, cellular compartments, RNA processing, ribosome biogenesis and mitochondrial translation. These gene categories found in metabolically active cells reflect the differentiation processes occurring between D0 and D10 and indicate a relatively reduced metabolism on D14. The later cluster had few GSEA attributes. One of these was “autophagy” (GO: 0006914) (26 genes, 3.1-fold enrichment, FDR=0.008). In this group, 24 genes were involved in various forms of autophagy (mitophagy, xenophagy, chaperone-mediated autophagy). This progressive overexpression of genes associated with autophagy during MK differentiation might reflect the implementation in MKs of the control of autophagy-dependent hemostatic functions in platelets (15), or could suggest that autophagic functions participate in the optimal differentiation of MKs, for example by eliminating midbodies at the end of endomitosis.

Cluster 4 contained 837 genes progressively downregulated between D0 and D14 but was characterized by a small number of GSEA attributes. These were related to mRNA splicing or RNA processing (36 genes), suggesting that the maturation of MKs is accompanied by changes in the regulation of alternative RNA splicing, a topic which could not be investigated using the experimental strategy chosen for this study.

In summary, genes associated with MK and platelet biogenesis and functions were mostly upregulated during the D0-D7 and D10-D14 periods (clusters 1 and 2). Although these conclusions agree with the respective binary analyses, the gene sets poorly overlaid; among the 388 genes shown to be overexpressed during the D0-D7 and D10-D14 periods (Supplemental Figure 3b), only 75 were present in cluster 1, and 57 in cluster 2. This demonstrates the biological complementarity of the arithmetic (binary) and geometric (fuzzy C-means clustering) analyses.

### A3-Unproductive subsets

The D10MKu_34-_ or D10MKu_9+_ cell populations were unable to mature into proplatelet-forming MKs under our experimental conditions. As mentioned above, SR1 did not significantly affect the transcriptome of these cells, so we chose to compare these populations to the productive SR1^+^ D10 or D14MKp cells.

#### A3a- D10MKu_34-_ or D10MKu_9+_ cells display maturation profiles intermediate between those of D10 and D14MKp cells

Relatively to D10MKp cells, 1,126 genes were better expressed in D10MKu_34-_ cells (|log_2_FC|>log_2_1.5, ApVal <0.05) and, the associated enriched attributes were related to platelet or MK biology (platelet organization and activity, hemostasis, chemokines expression, organization of the membranes and of the cytoskeleton, adhesion) ***(supplemental files Tables 19, 20***). On the other hand, the 1244 genes less expressed in the D10MKu_34-_ cells were over-represented in the gene lists associated to DNA replication, cell cycle, ribosome biogenesis.

Comparison of the D10MKu_34-_ cells to the more mature D14MKp cells (log_2_FC>0.58, ApVal< 0.05) resulted in a somewhat reversed pattern of attributes (***supplemental file Tables 21, 22***). The 775 genes better expressed in the D10MKu_34-_ cells enriched cRGSU-biased pathways, metabolic, DNA replication terms, whereas the 1476 less expressed genes mostly enriched attributes related to platelet biology, the organization of the cytoskeleton and membranes, cholesterol biosynthesis and, the participation of Rho GTPases. Of note, the best-enriched category was related to histone H3-K4 monomethylation (10-fold enrichment, FDR=0.04), suggesting a possible contribution of epigenetic control of gene expression to the different phenotype of the D14 productive cells.

Thus, the D10MKu_34-_ cells appeared to display a gene expression landscape between those of SR1^+^ D10MKp and D14MKp cells. Analysis of the D10MKu_9+_ cells also revealed a similar intermediate expression pattern (|log_2_FC|>log_2_1.5, ApVal <0.05) (***supplemental files Tables 23, 24,25, 26***). In other words, the D10MKu_34-_ and D10MKu_9+_ cells displayed a more mature but incomplete MK phenotype.

#### A3b- D10MKu_34-_ and D10MKu_9+_ cells mostly differ in cellular functions

Respectively 1085 and 345 genes were better or less expressed in D10MKu_9+_ cells, as compared to D10MKu_34-_ cells. The 1,085-gene list of uDEGs was enriched in GO BP terms related to chromatin organization and negative regulation of transcription, in cell cortex, microtubules and nucleus GO CC terms, but not in Reactome pathways (***supplemental files Tables 27, 28***). Thus, in terms of GSEA attributes, the 2 unproductive subsets mostly differed in cellular biological functions.

#### A3c- Shared and distinctive features between unproductive cells and productive cells (supplemental files Tables 38-43)

The preceding binary analyses were then completed by the comparison of D10 unproductive and the D10 and D14 productive cells obtained in the presence of SR1. After choosing an FDR <0.02 threshold, 1712 DEGs were selected. For this analysis, a number of 5 expression profiles was empirically chosen (**Figure 6**).

**Figure 6.**
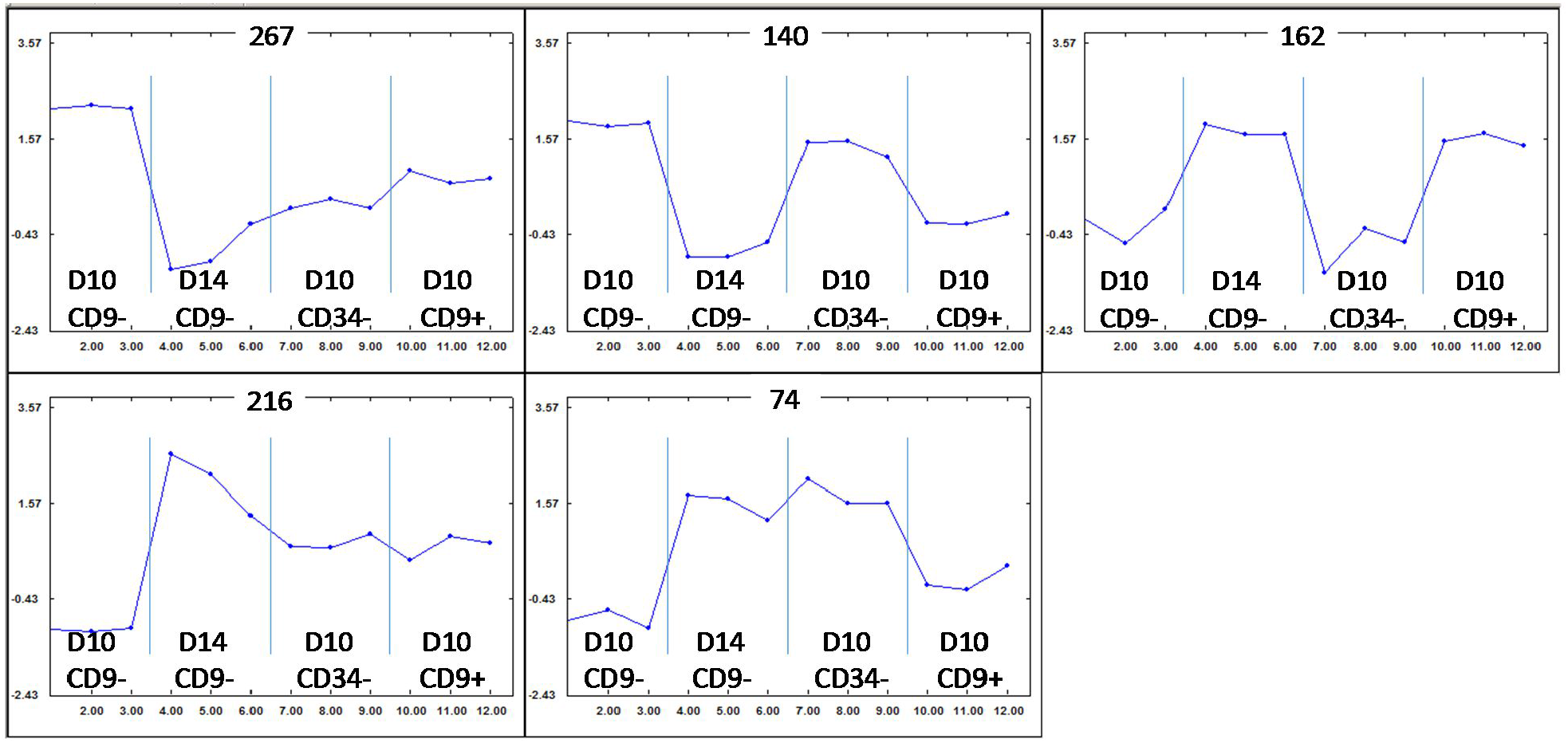
Fuzzy C-means clustering of DEGs between D10 and D14 MKp cells, D10MKu_34-_ and D10MKu_9+_ cells. Expression values were analyzed by ANOVA, genes with a differential expression pattern displaying an FDR<0.02 were selected (1712 genes). The expression patterns were unraveled by fuzzy C-means clustering after having chosen 5 expression patterns. For each expression pattern or cluster, the numbers of genes in the clusters are indicated.

The 267-gene cluster represented genes better or less expressed in the unproductive cells than in D10 or D14 MKp cells, respectively. So, the GSEA attributes, related to mitosis and DNA replication-associated processes, reflected the respective binary analyses. An interesting gene belonging to this cluster was DEPTOR, a major component and negative regulator of the mTOR complex 1 and 2, which negatively regulates the common subunit mTOR kinase (RPTOR) (16).

The 216-gene cluster was characterized by an inversed profile and, GSEA attributes were related to megakaryocytopoiesis/thrombopoiesis processes, platelet functions, extracellular matrix or cell adhesion or signaling. Of note, PTPRJ, a protein phosphatase receptor known to be important in megakaryocytopoiesis/thrombopoiesis, belonged to this cluster and, was associated to many enriched GO BP terms and Reactome pathways (regulation of cell-matrix adhesion, regulation of protein kinase B signaling, positive regulation of MAPK cascade, 62 attributes out 139). Thus, the expression of this major regulator of megakaryocytopoiesis might appear to be inappropriately expressed in unproductive cells, as compared to the productive ones.

The 3 other expression profiles appeared to be associated to combinations of intermediate expression profiles, reflecting the features shared between one and only one unproductive cell type and D10 or D14 productive cells. The 140-gene cluster included genes downregulated between D10 and D14 MKp cells, whereas the expression in D10MKu_34-_ and MKu_9+_ cells resembles D10 and D14 MKp expression, respectively. This cluster was enriched in genes involved in metabolism or associated to GO terms or Reactome pathways biased by the presence of a relatively high number of cRSUGs and PSMGs.

The 162-gene cluster displayed a symmetric profile, with genes upregulated between D10 and D14 in MKp cells, whereas the expressions in D10MKu_34-_ and MKu_9+_ cells resemble D10 and D14 MKp expression, respectively. Among them, 40 were associated to the regulation of transcription, including LYL1 and ZFPM1, pointing to direct contribution of TFRs to the differential megakaryocytic features of these cells.

Last, the 74-gene cluster displayed an antisymmetric regulation pattern (displaying a close expression profile in D10MKu_34-_ and in mature D14MKp cells, but an intermediate one in D10MKu_9+_ cells with respect to those upregulated between D10 and D14 in MKp cells). It was enriched in genes participating to organelle homeostasis, in particular GO:0031201 “SNARE complex” (fold enrichment score =26). Of note, 4 DEGs are associated to this term, BET1, NAPB, SNAP23 and STX7, and, according to Bioplex plateform (https://bioplex.hms.harvard.edu/) (17), the product of the first 3 interact with the product of the 4^th^. This correlation suggests a key role of these proteins during MK differentiation.

This analysis thus confirmed that the unproductive cells displayed a common intermediate profile of cell cycle-related downregulated genes, platelet- and MK-related upregulated ones, but displayed distinctive expression profiles for genes involved RSUG-, transcription-associated processes.

### B - ChEA3 analysis

We finally investigated how the DEG lists identified by the binary analyses could be correlated with the activity of transcription factors or regulators (TFRs). For this purpose, the ChEA3 algorithm was used, focusing on the calculated mean rank tables. The tables of results included 1632 TFRs, whereby those unlikely to be expressed in the majority of the cells - arbitrarily defined as displaying a mean expression value of <5 for the two conditions of the respective binary analysis - were not taken into account; below, by abuse of language, the TFRs filtered out are qualified as “expressed”.

### B1 - Analysis of genes controlling megakaryocytopoiesis (Supplemental Tables 44-61)

We first focused on the ChEA3-calculated mean scores of MK TFs reported to regulate megakaryocytopoiesis, namely ETV6, FLI1, GATA1, GATA2, GFI1B, IKZF1, IKZF5, LYL1, MECOM, MEIS1, NFE2, RUNX1, TAL1 and ZFPM1. The recently identified factor ZNF648 (18) was not included in the analysis because its expression was virtually undetectable under all conditions.

Analysis of the productive pathway in the absence of SR1 showed that the upregulation profiles associated with the D0-D7, D7-D10 and D10-D14 steps could be correlated with the activity of respectively (i) TAL1 or GATA1, (ii) ZFPM1 and (iii) GATA1, TAL1, GFI1B, LYL1, NFE2 or FLI1 (based on a cut-off score of <100) (*Supplemental Figure 6*). Notably, GATA1, TAL1, GFI1B, LYL1 and ZFPM1 were found to be significantly upregulated when their involvement was analyzed using ChEA3. In the presence of SR1, D10-D14 uDEGs were further found to be co-regulated not only by the TFs of group (iii), but also by ZFPM1, while except for GATA1, these TFs were again significantly upregulated. In agreement with these findings, when directly comparing SR1^-^ and SR1^+^ D10MKp cells, among the 265 genes less strongly expressed under the latter condition - in other words, those displaying delayed MK differentiation - a subset of 58 genes was seen to be co-regulated by/with ZFPM1 activity. Since the score of ZFPM1 for the SR1^-^ D10-D14 step was poor (440.2), these analyses suggested that the time- and SR1-dependent modulation of ZFPM1 activity could engender differences between the cells generated under SR1^+^ and SR1^-^ conditions, in particular with respect to their ability to develop proplatelets later on.

We then looked at the ChEA3 profile of the genes differentially expressed between SR1^+^ D10 MKu_34-_ or MKu_9+_ cells and SR1^+^ D10 or D14 MKp cells. The differential expression profiles appeared to be associated with the activity of most of the MK TFs in a way that reflected the GSEA analyses, where D10 unproductive cells displayed a MK maturation profile intermediate between those of SR1^+^ D10 and D14 MKp cells. However, some distinctive features were associated with ZFPM1 activity: its involvement was correlated with the genes less strongly expressed in D10MKu_34-_ than in D14MKp cells, but not with those more strongly expressed in D10MKu_34-_ than in D10MKp cells. In contrast, ChEA3 analysis of the genes more or less strongly expressed in D10MKu_9+_ cells than in D10 or D14 MKp cells resulted in a qualitatively inversed ranking for ZFPM1, reflecting the more mature megakaryocytic profile of the former cells, as mentioned above.

Considering the intermediate megakaryocytic expression profile of D10MKu_34-_ cells, these observations suggest that the activity of ZFPM1 is suboptimal with respect to MK differentiation in D10MKu_34-_ but not in D10MKu_9+_ cells. Thus, the different megakaryocytic features of the two D10 unproductive cell populations might be linked to ZFPM1 activity.

### B2 - ChEA3 TFR profiling of D0-D7 DEG lists: the D0-D7 step is dominated by genes associated with cell cycle control and the involvement of ZNF367

The D0-D7 step highlighted relevant biological correlations, the D0-D7 uDEG and dDEG lists and their respective ChEA3-deduced TFR lists being biologically related, whereas the other phases were more difficult to interpret.

Among the 10 and 100 best-ranked TFRs found to be associated with the co-regulation of gene subsets from SR1^-^ D0-D7 uDEGs, respectively 9 and 22 were upregulated (|log_2_ FC| > 2, ApVal <0.001), while 0 and 8 were downregulated (*Supplemental Figure 7*). With respect to the D0-D7 uDEG list, the 11 ChEA3 best-ranked TFRs (ZNF367, FOXM1, CENPA, MYBL2, PA2G4, ZNF695, TFDP1, E2F7, DNMT1, PRMT3 and E2F1) displayed a highly significant difference|log_2_ FC|, between 1 and 7. The gene sub-lists associated with these TFRs shared 73 items; their GSEA highlighted cell division processes, with fold enrichment scores up to 16-fold higher than those obtained by GSEA of the complete uDEG list. These correlations are in agreement with the involvement of these attributes during the D0-D7 period, already revealed in the corresponding binary analyses, and with the known participation of most of these TFRs in cell cycle regulation (FOXM1, MYBL2, PA2G4, TFDP1, E2F7, E2F1 and DNMT1, as checked using GeneCards®). Interestingly, in the literature, the best-ranked TFR, ZNF367, often features in cancer studies and was originally identified as a TF which controls the expression of erythroid genes (19). Moreover, GSEA of the ZNF367-associated uDEG sub-list not only pointed to cell cycle-associated attributes but also to others related to “platelet activation” or “degranulation” and to “regulation of megakaryocyte differentiation”, with enrichment scores higher than those obtained by GSEA of the complete uDEG list. This analysis thus revealed that ZNF367 deserves attention because it might also control the commitment and/or differentiation of MKs from hematopoietic precursors, a property not yet documented.

Among the 10 and 100 ChEA3 best-ranked TFRs associated with dDEGs, respectively 9 and 56 were significantly downregulated between D0 and D7 (|log_2_ FC| >2, ApVal <0.001), while only 1 was upregulated (***Supplemental Figure 8***). Analysis of the gene lists associated with the top 10 TFRs indicated that they were enriched in genes related to lymphocytes, monocytes and CD33^+^ myeloid cells, reflecting the loss of committed or contaminating non-megakaryocytic cells during the first week (ENRICHR analysis, data not shown).

These features support the biological relevance of the ChEA3 analysis of D0-D7 DEGs and emphasize the potential role of ZNF367 during the early steps of *in vitro* megakaryopoiesis, through its effects on bipotent erythroid-megakaryocytic precursors.

## Discussion

The present work investigated the genetic basis of the improvement of a well standardized *in vitro* differentiation protocol to produce MKs from peripheral CD34^+^ progenitors (**Figure 1**). This improvement relies on co-culture with mesenchymal cells, or addition of an AHR antagonist (SR1) mimicking aspects of the *in vivo* stromal environment. The differentiation is accompanied by the sequential development of cell populations phenotypically defined as CD34^+^CD41^+^ on D7 (D7MKp cells) and CD34^+^CD41^+^CD9^-^ on D10 and D14 (D10/D14MKp cells), which then produce proplatelets between D14 and D17. On D10 two other populations expressing megakaryocytic markers are present but are unable to mature into proplatelet-forming cells (unproductive CD34^-^CD41^+^ and CD34^+^CD41^+^CD9^+^, respectively D10MKu_34-_ and D10MKu_9+_ cells). The AHR antagonist boosts the yield of D10MKp cells and the ability of D14MKp cells to generate proplatelets, but has no obvious effects on the unproductive cells. The first aim of this work was to determine how the megakaryocytic gene expression program is progressively implemented; the second was to clarify why the different D10 and D14 cell populations, under the same culture conditions, mature into MKs more or less capable of producing proplatelets, and thus to elucidate the impact of SR1 on MK differentiation (**Figure 1**).

The analysis of the gene expression profile of megakaryocytes is a subject of intensive research. In December 2020, a PUBMED search found 132 papers about “megakaryocyte and transcriptome” and 345 about “megakaryocyte and gene profiling”. To our knowledge, this work represents the first analysis of the progressive *in vitro* differentiation of CD34^+^ progenitors into MKs at the level of the transcriptome of phenotypically defined cell populations. Since such an approach allows the analysis of a most extensive gene repertoire, it is complementary to published single cell transcriptomic studies, which provide a fine description of the cell diversity but in practice are based on the assessment of at least 10 times fewer genes. Further studies will be useful to clarify the correlations between the datasets generated using the two techniques.

Binary analyses (**Figure 3**) were preferred to compare the cell populations obtained at the different steps of *in vitro* culture and to investigate the effects of SR1. The resulting uDEG and dDEG lists were analyzed independently for enriched GO BP and GO CC or Reactome pathway attributes and for TFRs associated with the co-regulation of gene subsets from the lists (ChEA3). Thus, the differential expression profiles associated with the D0-D7, D7-D10 and D10-D14 steps of the productive pathway (MKp cells) were found to be enriched in distinctive combinations of biological attributes reflecting aspects of MK differentiation. Moreover, the transcriptomes of D7MKp and D14MKp cells appeared to be SR1-independent, in other words, SR1 seemed to have a major effect only on D10MKp cells

### The D0-D7 commitment period is associated with an SR1-independent gene expression profile

As numerous and important genetic modulations accompanied the D0-D7 phase, stringent differential expression thresholds needed to be used to extract meaningful information (|log_2_ FC| > 2, ApVal <0.001). The large number of metabolic and cell division-associated enriched attributes reflected the *in vitro* expansion of the cultured cells. Moreover, the induction or overexpression of 142 genes enriching GO BP, GO CC or Reactome attributes associated with hemostasis, response to wounding and platelet and MK biology documented the megakaryocytic commitment of the D7 cells. Use of the ChEA3 algorithm to rank TFRs predicted to regulate the network of D0-D7 uDEGs confirmed this megakaryocytic commitment, since TFRs known to regulate megakaryocytopoiesis (TAL1, GATA1, GFI1B, MEIS1 and NFE2) were ranked within the first decile.

Genes encoding cRSUGs were over-represented in the dDEG list, so that many translation-related or - unrelated GSEA attributes were selected. Consequently, the ribosomal architecture might be predicted to be dramatically modified during the culture of CD34^+^ progenitors. Since ribosomal composition regulates translation (20), this observation raises the question as to whether the control of translation is biologically a major mechanism of MK commitment and/or of the control of the quiescence of CD34^+^ progenitors. Of note, hematopoietic stem cells (HSCs) were previously shown to need a tightly regulated protein synthesis rate (21). Alternatively, as many cytosolic ribosomal subunits are moonlighting proteins, in other words, participate in processes other than the translation of mRNAs, e.g., modulate cellular responses to particular stimuli, their downregulation might disturb biochemical pathways in the cells.

The absence of a significant effect of SR1 on the gene expression profile of D7MKp cells suggests that its effects between D0 and D7 on the generation of proplatelet-forming D14MKp cells could result from its action on a numerically minor MK precursor. Indeed, the lack of a significant effect of SR1 on the gene expression profiles of the unproductive D10MKu_34-_ and D10MKu_9+_ subsets, which are unable to differentiate into proplatelet-forming cells, points to a cell specificity of SR1 with respect to MK differentiation. A single cell transcriptome analysis might be required to identify this putative SR1-sensitive MK precursor.

### The D7-D10 step, an intermediate period: many DEGs but few SR1^-^ GSEA attributes; few SR1^+^ DEGs, but lipid and cholesterol biosynthesis-associated dDEGs, as under SR1^-^ conditions

Under SR1^-^ conditions, the D7-D10 phase was accompanied by the up- or down-regulation of respectively 1763 and 1034 genes in MKp cells (|log_2_ FC| > log_2_ 1.5, ApVal <0.02). In contrast, the numbers of DEGs identified under SR1^+^ conditions were comparatively very low - 32 uDEGs and 67 dDEGs.

Surprisingly, in the absence of SR1, although the number uDEGs was relatively high, only a limited number of GSEA attributes were associated with these genes. The enriched GO BP terms were related to endosomal processes, autophagy, regulation of GTPase activity (60 genes) and negative regulation of transcription (123 genes). These processes have many consequences for cell biology and might affect the fate of the cells with respect to MK differentiation.

Although dDEGs were markedly less numerous, their GSEA highlighted SR1^-^-specific enriched metabolic processes including mitochondrial respiration processes, in particular ATP production, RNA splicing and mitotic mechanisms. The early combined downregulation of ATP biosynthesis and mitotic functions might be detrimental to downstream endomitotic mechanisms which are essential for MK maturation.

Interestingly, the downregulation of lipid and cholesterol biosynthetic pathways under SR1^-^ and SR1^+^ conditions, in spite of the small number of dDEGs for the latter condition, supports a biological relevance of the modulation of these pathways during the intermediate D7-D10 period of MK differentiation *(Supplemental Figures 9 and 10)*. Genes of the lipid biosynthetic pathway participate in the synthesis of membrane-associated lipids; in particular, the downregulated SPTLC3 encodes a subunit of the rate limiting enzyme complex involved in *de novo* sphingolipid synthesis. These unexpected modulations suggest that the development of MKs is accompanied by a change in the composition of their membrane lipids (gangliosides and long-chain fatty acid-containing lipids), which could be delayed by SR1. The downregulation of cholesterol biosynthesis under SR1^-^ conditions might result from the simultaneous downregulation of mitochondrial energy production (16). However, this hypothesis cannot be invoked for SR1^+^ conditions, where mitochondrial metabolism was not significantly altered. Therefore, at least in the presence of SR1, the downregulation of cholesterol biosynthesis must be controlled by other or additional regulatory mechanisms.

Other stimulating issues emerge from a deep analysis of the cholesterol biosynthetic pathway in D7MKp cells. Several enzymes of the mevalonate-lanosterol pathway, in particular the two rate limiting enzymes HMGCR and SQE (22), were downregulated in the presence or absence of SR1, with no statistically significant difference between the two conditions. This downregulation could result in a decrease in the amount of lanosterol in MKp cells, which would lead to a reduction in membrane fluidity and the signaling processes dependent on it (23). Furthermore, most of the genes participating in the downstream Bloch and Kandutsch-Russel pathways of cholesterol biosynthesis were also downregulated between D7 and D10. One surprising exception to this general trend was the upregulation of CYP51A1 under SR1^-^ conditions and its marginal downregulation in the presence of SR1. An anticipated consequence of the upregulation of CYP51A1 would be an exacerbation of the decrease in the cellular content of lanosterol, its substrate, under SR1^-^ but not SR1^+^ conditions, with its resultant cellular effects. Since enzymes downstream of the two cholesterol biosynthetic pathways were downregulated, CYP51A1 products might also accumulate differently in the cells generated under SR1^-^ and SR1^+^ conditions, a poorly characterized topic which needs to be clarified. We propose that these different effects on cholesterol biosynthesis could contribute to the impact of SR1; illustrating the role of cholesterol in AHR-dependent responses, it was recently found to participate in the modulation of the AHR-mediated expression of inflammatory cytokines in macrophages (24). Additional experiments will be necessary to clarify the biological consequences with respect to MK differentiation.

### SR1^-^ and SR1^+^ D14MKp cells display the same gene expression pattern; common and distinct features of the SR1^-^ and SR1^+^ D10-D14 periods

The transcriptomes of the D14 cells obtained in the presence or absence of SR1 were virtually identical. Indeed, even the use of low statistical thresholds (|log_2_ FC| > log_2_ 1.5, ApVal <0.05) identified only 7 genes differently expressed between SR1^-^ and SR1^+^ D14MKp cells. Among the 5 upregulated genes, ARHGAP24 and MICAL2 could be relevant to the biology of mature MKs. However, as the statistical significance of the differential expression was marginally lower than 0.05 (**A2a**), complementary experiments will be required to confirm the differences in the regulation of these genes.

Reflecting the differences between the transcriptomes of SR1^-^ and SR1^+^ D10MKp cells, about two thirds of the DEGs were shared between SR1^-^ and SR1^+^ conditions during the D10-D14 period (|log_2_ FC| > 0.8, ApVal <0.01; *<u>Supplemental Figure 3a</u>*). Thus, GSEA revealed both common and distinct attributes associated with these conditions. The uDEGs shared processes regulating the last events of MK biology, when MKs cross the endothelial barrier, and/or platelet functions such as signaling, cytoskeletal and membrane dynamics, podosomes and interaction with the extracellular matrix or endothelial cells. Notably, cholesterol biosynthesis appeared to be upregulated between D10 and D14. Hence the D7-D10 downregulation described above was transient. The common dDEG attributes were strongly associated with cell cycle, DNA replication or repair processes. This downregulation probably reflects the arrest of cell division during the D10-D14 period. On the other hand, how these regulatory events affect endomitosis, which occurs during and after this step, is a question that needs further investigation, the mechanism of endomitosis in MKs being as yet poorly characterized.

### Several genes differently expressed in SR1^-^ vs SR1^+^ D10MKp cells and differently expressed between D10 and D14 control key megakaryocytic functions

Given the similarities of the D14MKp cells, the specificities of the D10-D14 step under SR1^-^ and SR1^+^ conditions were analyzed by focusing on the genes differently expressed between the SR1^-^ vs SR1^+^ D10MKp cells. Although low thresholds were used (|log_2_ FC| > log_2_ 1.5, ApVal <0.05), only a few hundred genes were differently expressed (221 uDEGs and 265 dDEGs). These genes seemed to display SR1^+^ D10 expression values lying between SR1^-^ D7 and D10 expression values, before recovering D14 ones. Thus, SR1 appeared to transiently delay the maturation of MKp cells, this delay being caught up by D14.

During D10-D14 megakaryocytic differentiation, many biological pathways seemed to be recruited, qualitatively in an SR1-independent manner (with respect to GO terms or Reactome pathways), but quantitatively in an SR1- and time-dependent manner (with respect to the associated gene lists and FCs). This conclusion can be illustrated by focusing on the genes differently expressed between SR1^-^ and SR1^+^ conditions in D10MKp cells and overexpressed between D10 and D14 in the presence of SR1 (**A2e**; ***Supplemental Table 29***). Several of these genes might play key roles during the later steps of megakaryocytopoiesis or the early ones of thrombopoiesis, some of them pointing to unsuspected processes. One may cite two examples: CKLF, a poorly characterized chemokine which could regulate cellular homeostasis in the bone marrow, or ICAM5, a neuron intercellular adhesion molecule, which here might control the interaction of MKs with stromal cells, endothelial cells, the extracellular matrix or leukocytes, or even explain how emperipolesis occurs. Obviously, ICAM5 could also have important functions in platelet pathophysiology and its expression in these cells deserves to be more closely investigated.

Another gene displaying a similar upregulation profile was PLXNB3. Interestingly, two semaphorin/plexin-dependent attributes, semaphorin interactions (13/64 genes) and parental pathway “axon guidance” (56/389 genes), were present among the D10 and D14 uDEGs, independently of the presence or not of SR1 during differentiation (***Supplemental Figure 11***). We note that PLXNP3 ligands are expressed by ECs (semaphorins 4A and 5A (11)) and by MKp cells themselves (SEMA4B and SEMA4D). Therefore, it is tempting to speculate that semaphorin/plexin interactions could participate in megakaryocytopoiesis and/or thrombopoiesis, *in situ* in the bone marrow or spleen, through interactions with counter receptors expressed by stromal cells (endothelial and/or mesenchymal cells). In particular, enrichment of the axon guidance pathway could suggest by analogy a “proplatelet formation pathway” in relationship with the environment, although up until now, proplatelet formation has been considered to be an MK-autonomous process. Alternatively, in maturing and/or mature MKs, when the DMS expansion allows a particular 3D-organization of the plasma membrane, inter-membrane interactions might occur via these plexins and semaphorins, resulting in the activation of internal pathways, while disruption of such interactions might promote or contribute to the formation of proplatelets. Consistent with this hypothesis, when MKs differentiated under conditions promoting the formation of a morphologically *bona fide* DMS are transferred to free conditions, the release of mechanical constraints leads to DMS remodeling followed by the formation of proplatelets (25).

### The unproductive cell subsets display an intermediate MK maturation profile

The D10MKu_34-_ and D10MKu_9+_ cell populations do not generate proplatelets under our cell culture conditions, their transcriptomes were SR1-independent and intermediate between those of the D10 and D14 MKp cells generated under SR1^+^ conditions, like the D10MKp cells obtained in the absence of SR1. GSEA of the genes differently expressed between D10MKu_34-_ and D10MKu_9+_ cells mostly revealed cellular attributes. Comparison of these cells with D10 and D14 MKp cells (**A3c**) identified five gene expression patterns characterized by different combinations of up- or downregulation profiles, depending on the cell type and period. For example, in the unproductive cells, 267 genes displayed a downregulation to levels intermediate between those observed in D10 and D14 MKp cells. The levels of expression of these genes, which were mainly involved in DNA replication processes, might be too low to allow the cells to progress towards a more mature phenotype. A 216 cluster displayed the inverse expression profile and included genes participating in megakaryocytopoiesis/thrombopoiesis and/or platelet functions. One of these was PTPRJ, which encodes a protein whose deficiency results in MK and platelet defects. This gene was associated with half of the attributes specific to this cluster; hence its relatively low expression could contribute to the unproductive properties of D10MKu_34-_ or D10MKu_9+_ cells. Another 162-gene cluster pointed to possible mechanisms underlying the differences between the two unproductive cell types. This group included genes exhibiting expression values in D10 MKu_34-_ and MKu_9+_ cells close to those in D10 or D14 MKp cells, respectively. Among these genes, 40 were associated with attributes related to the regulation of transcription, including the megakaryocytic TFs LYL1 and ZFPM1. Consistent with this result, ChEA3 analysis of co-regulated genes more or less strongly expressed in unproductive cells than in D10 or D14 MKp cells, respectively, highlighted a positive correlation with ZFPM1 in D10MKu_9+_ cells, but not in D10MKu_34-_ cells.

### The differential expression profiles point to a key regulatory role of mTORCs during MK differentiation

Comparison of the differential expression profiles of the productive and unproductive cells showed that DEPTOR was strongly downregulated between D10 and D14 in productive cells, whereas in D10 unproductive cells it displayed an intermediate expression which was nevertheless close to that in D14 cells. The particular expression profile of this inhibitor of the mTOR pathway prompted us to check the expression patterns of the constituents of the mTOR complexes. DDIT4, another negative regulator of mTORC1, also displayed an unusual expression profile, being better expressed in all D10 cells than in D7 or D14 cells. Thus, the negative regulation of mTORC1 and mTORC2 would appear to be differently controlled in D14 cells on the one hand, and in D10 unproductive or productive cells on the other hand (***Supplemental Figure 12***).

D10 and D14 MKp cells should therefore differ in their responses to the numerous mTORC-dependent pathways (16, 26), suggesting that mTORC1/mTORC2 could be key regulators of the final steps of megakaryocytopoiesis. Several of these pathways are positively regulated at the level of mRNA translation by mTORC1, for example nucleotide synthesis (27), which may be expected to be upregulated during the D10-D14 maturation step of MKp cells when polyploidy increases (1), or the biosynthesis of lipids and cholesterol, which are necessary for DMS biogenesis during the later steps of megakaryocytopoiesis. Both mTORC1 and mTORC2 activate SREBP, a master regulator of cholesterol neosynthesis; hence the downregulation of the two inhibitors of these complexes is in agreement with the observed induction of genes of the cholesterol biosynthetic pathway between D10 and D14 in MKp cells. Furthermore, mTORC1 positively regulates the expression of survival genes, which might be required to complete the final steps of megakaryocytopoiesis, given the atypical biology of mature MKs, although the literature is conflictual on this point. On the other hand, mTORC2 modulates among other metabolic processes, cytoskeletal organization and directly or indirectly the activity and/or specificity of a plethora of protein kinases which could be important at different levels, for example for control of the development and architecture of the DMS, or the formation of podosomes (28). Moreover, the downregulation of DEPTOR might result in the increased phosphorylation, and thereby activation, of ERK1/2 and STAT1 in an mTOR-independent manner in endothelial cells, raising the question of the existence of further levels of post-transcriptional and post-translational regulation (16, 26, 29).

Thus, it remains to clarify how the anticipated enhanced activities of mTORC1- and mTORC2-dependent pathways could contribute to the later steps of megakaryocytopoiesis, not only *in vitro* but also *in vivo*. In particular, it will be necessary to identify their effectors in the bone marrow environment, the downstream effectors engaged in these pathways, the impact on the last steps of MK maturation and finally, the consequences for the mechanisms of thrombopoiesis. The major pathway of mTORC1 activation acts through starvation, which may not be physiologically relevant to megakaryocytopoiesis. Nevertheless, since mTORC1 and mTORC2 cross-regulate in a complex manner, mTORC2 activation could trigger that of mTORC1. Activation of mTORC2 requires the formation of phosphatidic acid, probably under the control of phospholipase D2 (PLD2) (30, 31), although another pathway involving diacylglycerol kinase ζ (DGKZ) activation could also be effective (32). Proteome studies have revealed the presence of DGKZ in human but not mouse platelets, but did not detect PLD2 in the platelets of either species (33, 34). Nonetheless, functional studies indicate that human (35) or mouse (36) platelets and in *in vitro* differentiated mouse MKs (37) express PLD2. The log_2_ mean expression value of PLD2 was low in all D7 to D14 cell subsets (between 5 and 6); in contrast, DGKZ expression increased 4-fold from D7 to D14 in productive cells, in agreement with its reported presence in platelets (***Supplemental Figure 12***).

In addition, mTORC2 signaling participates in the signal transduction initiated by tyrosine kinase receptors or GPCR (35), which in the context of megakaryocytopoiesis might be triggered by stromal cell products. It is also involved in mechanical membrane stretching (16), an event potentially encountered during podosome formation, before crossing of the endothelial barrier (14), or during proplatelet elongation *in vitro* or in the blood stream. The differences between *in vitro* and *in vivo* conditions might lead to differences in the activation pathways of mature MKs, which could result in different mechanisms of proplatelet generation, as recently documented (38). Notwithstanding, the SR1-independent expression profiles of DEPTOR and DDIT4 will need further elucidation.

### The absence of transcriptomic differences between SR1^-^ and SR1^+^ MKp cells on D14 suggests post-transcriptional memory mechanisms

The most unexpected and frustrating conclusion of this work was that the transcriptomes of the D14MKp cells obtained in the presence or absence of SR1 were virtually identical, although their respective biological potentials, namely their abilities to generate proplatelets, were clearly different. This paradoxical observation suggests firstly that the major biological determinants of the advantage of SR1 during the later phase of MK maturation are post-transcriptionally driven and secondly, that these determinants are programed during the preceding period.

This idea would lead us to propose an essential contribution of post-translational modifications of proteins, in connection with the signaling network-learning hypothesis (39). According to this model, prolonged or sustained stimulations can modify for days the homeostasis of signaling networks controlled by post-translational modifications. Our transcriptome studies showed that between D7 and D10, in the absence of SR1, the variations in gene expression were numerous (1034 dDEGs and 1763 uDEGs) and diverse (few uDEG-enriched attributes), whereas only 32 uDEGs and 67 dDEGs were identified for the same period under SR1^+^ conditions. Thus, the SR1^-^ D7-D10 step would be associated with dispensable genetic modulations which could even be detrimental to appropriate MK differentiation, a hypothesis supported by the period-specific regulation of genes participating in the control of GTPase (***Supplemental Figure 13***). As discussed above, mTORC-dependent pathways are also likely to be differently regulated in the various subpopulations. Therefore, cell signaling processes would be differently activated between D7 and D10 or later under SR1^-^ and SR1^+^ conditions, in terms of the effectors and timeframe. Only the cells differentiated in the presence of SR1 might adopt the appropriate coordination of biochemical pathways allowing them to produce D14 cells displaying a more *bona fide* MK phenotype (ploidy and DMS formation) (1), and capable of generating potent proplatelet-forming cells later on.

### General conclusion

This work represents a systematic transcriptomic analysis of the megakaryocytic cells developing during the *in vitro* differentiation of CD34^+^ progenitors. In addition, it describes the genetic effects of an AHR antagonist which mimics aspects of MK-mesenchymal cell interactions in the bone marrow. The study has wide implications, due to the diversity of the processes appearing to accompany MK differentiation, and should thus be of benefit to a diverse panel of scientists. Combining the data obtained here with single cell transcriptome studies should provide a better understanding of MK differentiation.

The apparent restriction of the specificity of SR1 to MKp cells during the D7-D10 period raises the question of whether it acts on particular MK progenitors. The data suggest that the unproductive cell subsets display a distinctive expression profile which may be inappropriate for proplatelet formation, although it remains to explain why these cells do not seem to be able to progress towards proplatelet-forming cells.

An unanticipated major paradox was that the D14MKp cells generated in the presence or absence of SR1 exhibited an identical transcriptomic profile, whereas they differentiated into mature MKs displaying marked differences in terms of proplatelet production. These observations point to the relevance of other processes not analyzed in this study, such as the differential expression of miRNA, whose importance is well documented (40), and post-translational events, some of which would support the memory-signaling hypothesis that could explain the different biological properties of the SR1^-^ and SR1^+^ D14MKp cells.

## Abbreviations

ApVal: adjusted pValue
dDEGs: downregulated (or overexpressed) differentially expressed genes
uDEGs: upregulated differentially expressed genes
DGE: differential gene expression
DMS: Demarcation membrane system
D7MKp: D7 megakaryocytic precursors
FC: fold change
FDR: false discovery rate
GSEA: Gene Set Enrichment Analysis
GO CC, GO BP: Gene Ontology Cellular Compartment, Gene Ontology Biological Process
PSMG: proteasome subunit gene
SR1: StemRegenin 1
TFR: Transcription Factors or Regulator
D10MKp or D14MKp: D10 or D14 precursor of productive MKs
D10MKu_34-_ or D10MKu_9+_: CD34^-^CD41^+^ or CD34^+^CD41^+^CD9^+^ unproductive megakaryocytic cells

## Acknowledgements

This work was supported a grant from by the Agence Nationale pour la Recherche, ANR-17-CE14-0001-01. RNA-SEQ data were generated at Genomeast, France. We thank D Plassard (Genomeast), for the DEseq2 analyses.

---

**Supplemental Figure 1.**
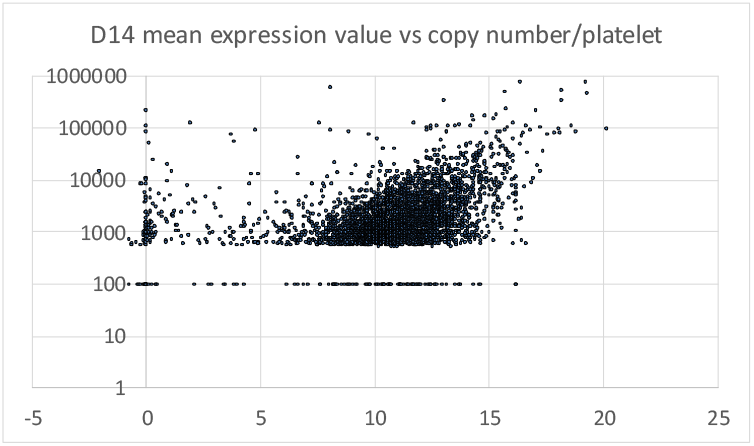
Distribution of the log_2_ normalized expression values/ number molecules per platelet. The proteomic data (Burkhart et al.) were curated by selecting proteins with corresponding HGNC gene name and provided quantitative data (3690). The number of molecules per platelet were plotted against the mean expression value in CD41^+^CD41^+^CD9^-^ cells at D14. For clarity, when the copy number of a protein was <500/platelet, a value of 100/platelet was plotted. So, 3652 proteins with a corresponding gene in the gene expression table were analyzed, of which 115 had an estimated copy number of <500 molecules/platelet, 196 had a mean expression value <5. Among these latter proteins, 38 were associated to liver tissue (ENRICHR analysis) 24 corresponding to immunoglobulins, thus overall displaying a strong bias with respect to plasma proteins (supplemental Table 3).

**Supplemental figure 2.**
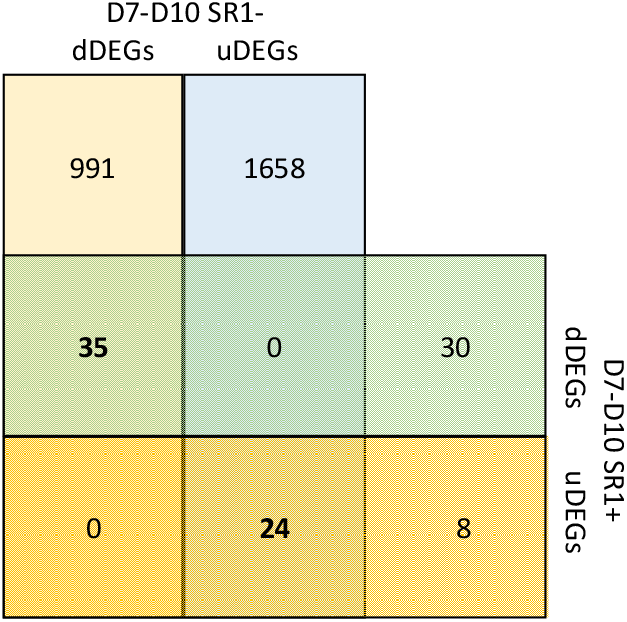
Venn diagram analysis of uDEG and dDEG lists associated to D7-D10 step in SR1^-^ and SR1^+^ conditions.

**Supplemental Figure 3a.**
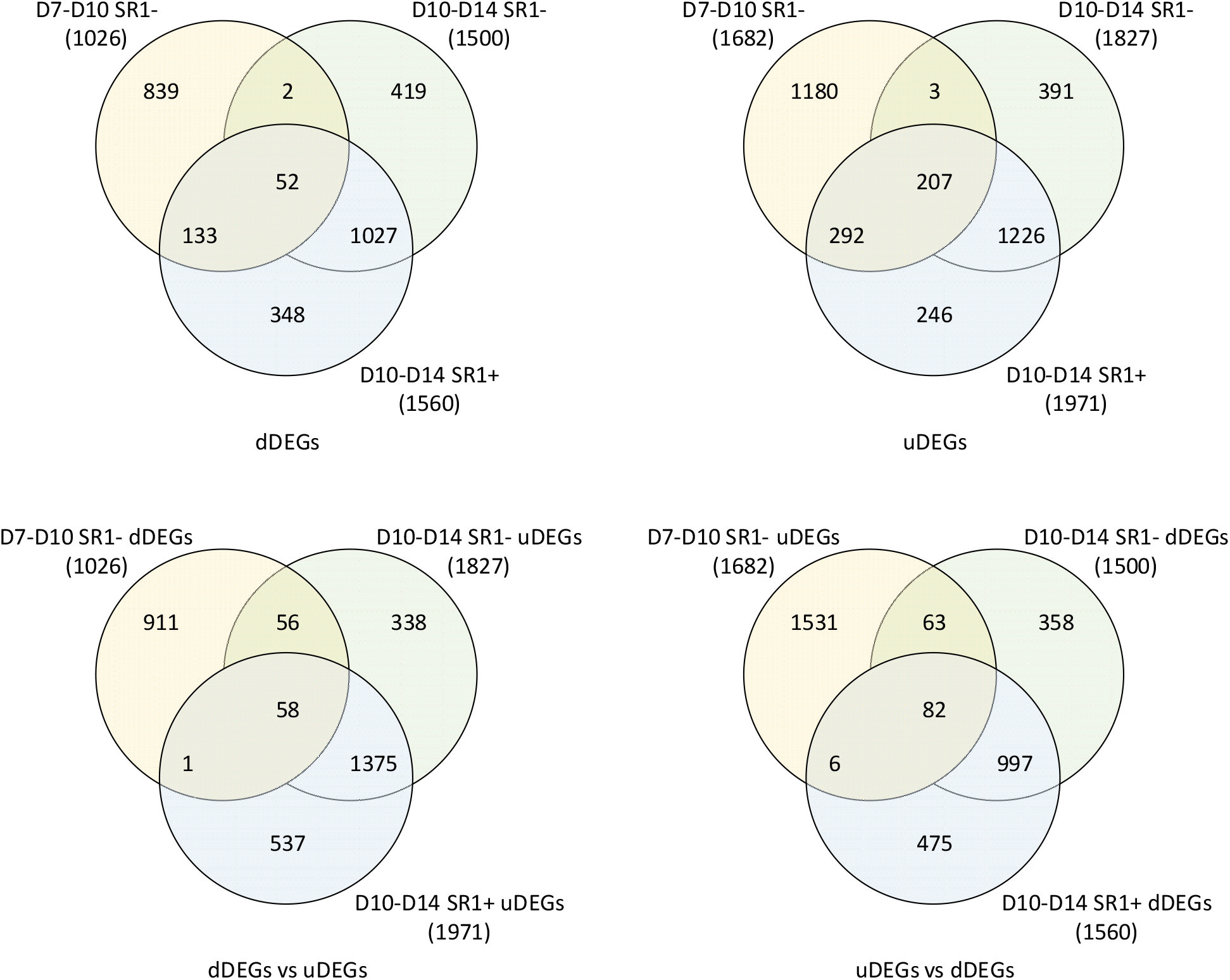
Venn diagram analysis of the distribution of the dDEGs and uDEGs in SR1^-^ (D7-D10 and D10-D14) and SR1^+^ (D10-D14) conditions between D10 and D14 (|Log_2_ FC|>0.8 and an ApVal<0.01). The SR1-D7-D10 dDEGs and uDEGs were compared to the SR1^-^ and SR1^+^ D10-D14 dDEGs and uDEGs, respectively (first row), or uDEGs and dDEGs, respectively (second row)

**Supplemental Figure 3b.**
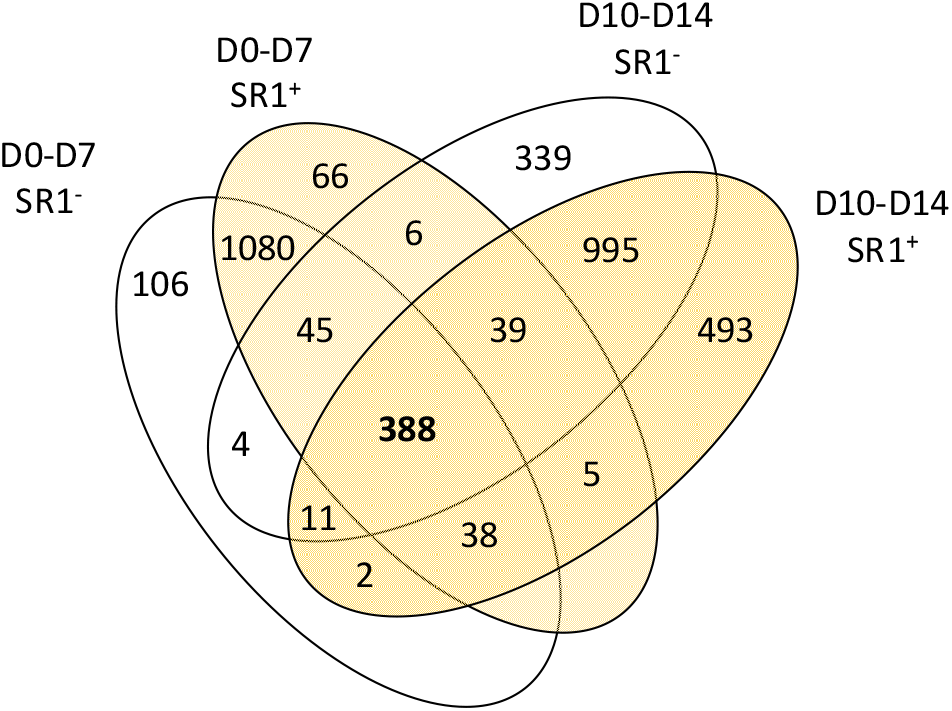
Venn diagram analysis of the distribution of the uDEGs during D7-D10 and D10-D14 periods and -Dunder SR1^-^ or SR1^+^ conditions (supplemental table 4), 388 genes are common to the 4 gene sets.

**Supplemental Table A.**
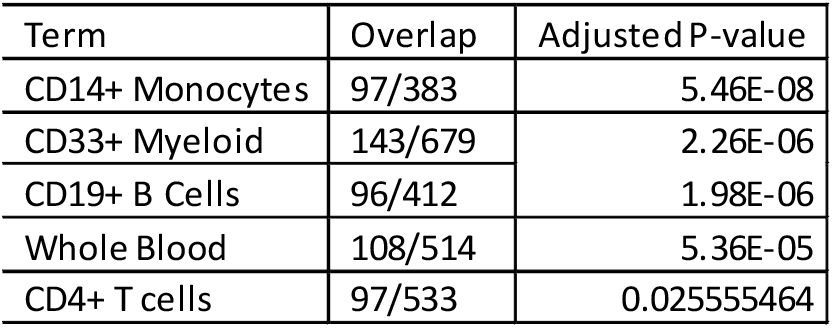
The 2740 dDEG list was analyzed using ENRICHR. A focus on Human Gene Atlas analysis revealed enrichments for genes expressed in lymphoid and myeloid cells. The overlap column provides the ratios of the number of genes present in the submitted list to the total number of genes inventoried in the respective cell type (term).

| Term | Overlap | Adjusted P-value |
| --- | --- | --- |
| CD14+ Monocytes | 97/383 | 5.46E-08 |
| CD33+ Myeloid | 143/679 | 2.26E-06 |
| CD19+ B Cells | 96/412 | 1.98E-06 |
| Whole Blood | 108/514 | 5.36E-05 |
| CD4+ T cells | 97/533 | 0.025555464 |

**Supplemental Table B.**
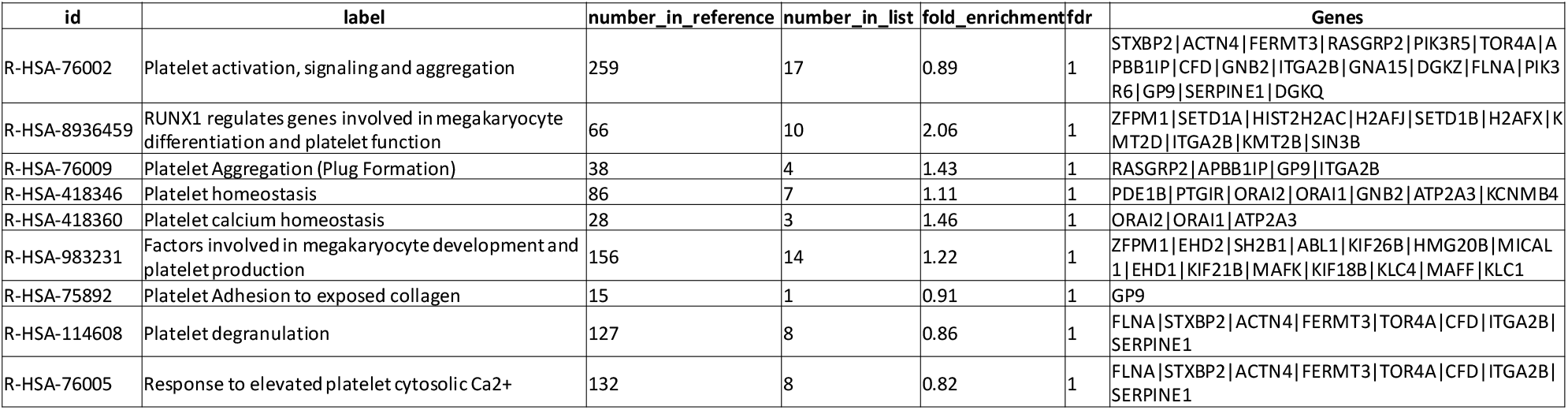
GSEA of SR1- D7-D10 uDEGs, platelet and megakaryocytic-related Reactome categories.

**Supplemental Table C.**
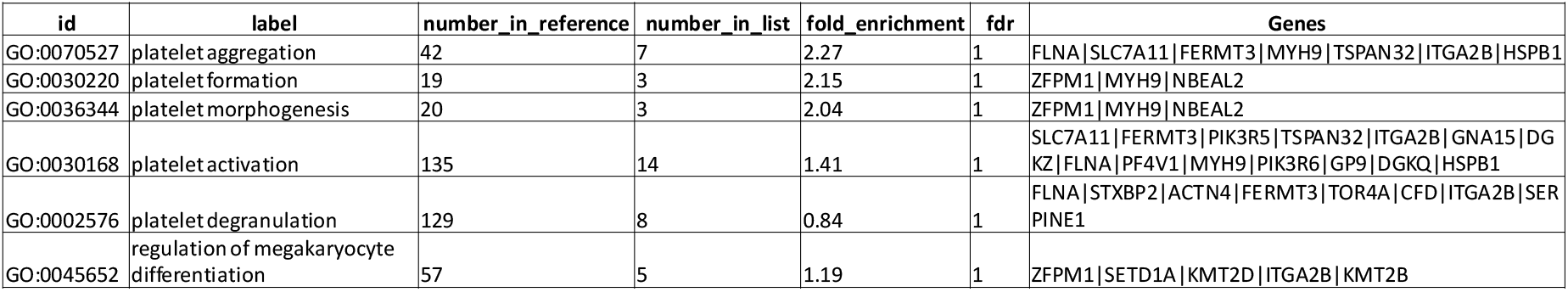
GSEA of SR1- D7-D10 uDEGs, platelet and megakaryocytic-related GO BP categories.

**Supplemental Figure 4a.**
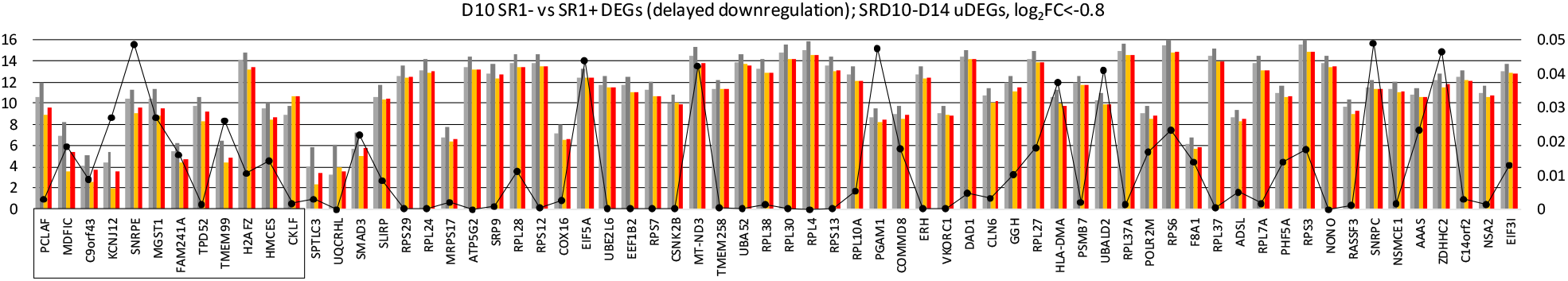
D10-D14 expression profiles in SR1^-^ or SR1^+^ condition of the DEGs better expressed at D10 in SR1^+^ condition (log_2_FC>log_2_1.5|, ApVal<0.05) and, differentially expressed between D10 and D14 in the SR1^+^ condition (|log2FC<0.8|, ApVal<0.01). The means of the triplicated normalized expression values at D10 SR1^-^ (light grey) or SR1^+^ (deep grey), D14 SR1^-^ (orange) or SR1^+^ (red) conditions are represented (scale, left axis). ApVals of the FCs of SR1^-^ vs SR1^+^ conditions at D10 are depicted by the broken line (scale, right axis). Genes were divided in two groups, those also differentially expressed in SR1^-^ condition (|log_2_FC<-0.8|, ApVal<0.01) (boxed), and the others. Within each group, the genes were plotted according to the decreasing values of the D10 SR1^-^ vs SR1^+^ FCs (D10 DEGs SR1- vs SR1_D10 D14_V2.xlsx).

**Supplemental Figure 4b.**
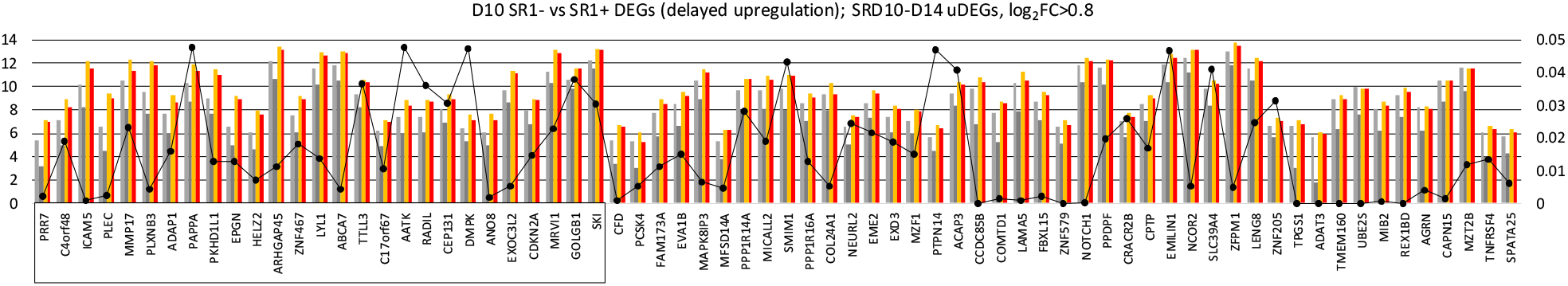
D10-D14 expression profiles in SR1^-^ or SR1^+^ condition of the DEGs less expressed at D10 in SR1^+^ condition (log_2_FC<-log_2_1.5, ApVal<0.05) and differentially expressed between D10 and D14 in the SR1^+^ condition (|log_2_FC|>0.8, ApVal<0.01). The means of the triplicated normalized expression values at D10 SR1^-^ (light grey) or SR1^+^ (deep grey), D14 SR1^-^ (orange) or SR1^+^ (red) conditions are represented (scale, left axis). ApVals of the FCs of SR1^-^ vs SR1^+^ conditions at D10 are depicted by the broken line (scale, right axis). Genes were divided in two groups, those also differentially expressed in SR1^-^ condition (|log_2_FC|>0.8, ApVal<0.01) (boxed), and the others. Within each group, the genes were plotted according to the decreasing values of the D10 SR1^-^ vs SR1^+^ FCs (D10 DEGs SR1- vs SR1_D10 D14_V2.xlsx). Only the first 70 genes were plotted.

**Supplemental Figure 5.**
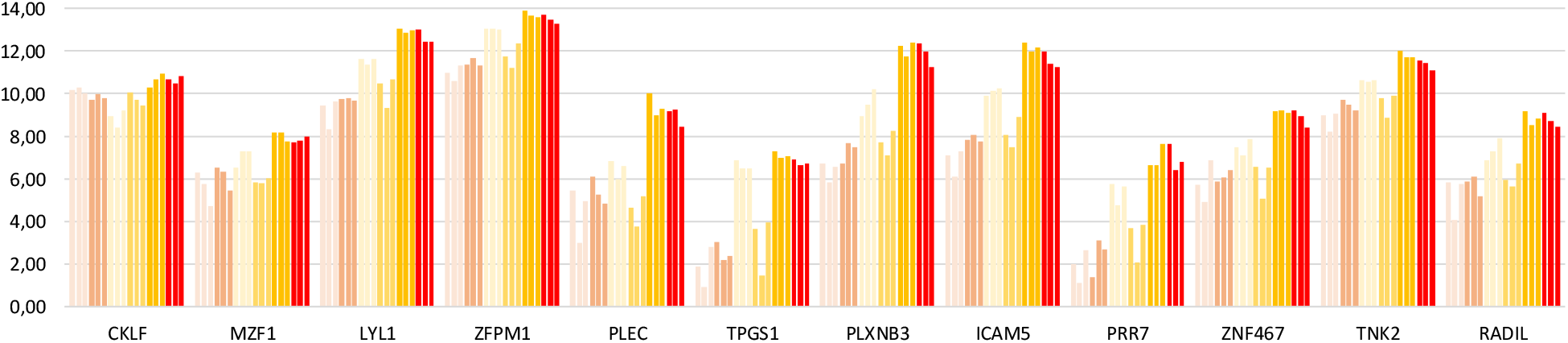
Expression profile of CKLF, MZF1, LYL1 and ZFPM1,. PLEC, TPGS1, PLXN3B, ICAM5, PRR7, ZNF467, TNK2, RADIL, CKLF. The triplicated normalized expression values are represented; from left to right D7 SR1^-^ and SR1^+^, D10 SR1^-^ and SR1^+^, D14 SR1^-^ and SR1^+^. Genes are ordered as they appear in the text (§A2e).

**Supplemental Figure 6.**
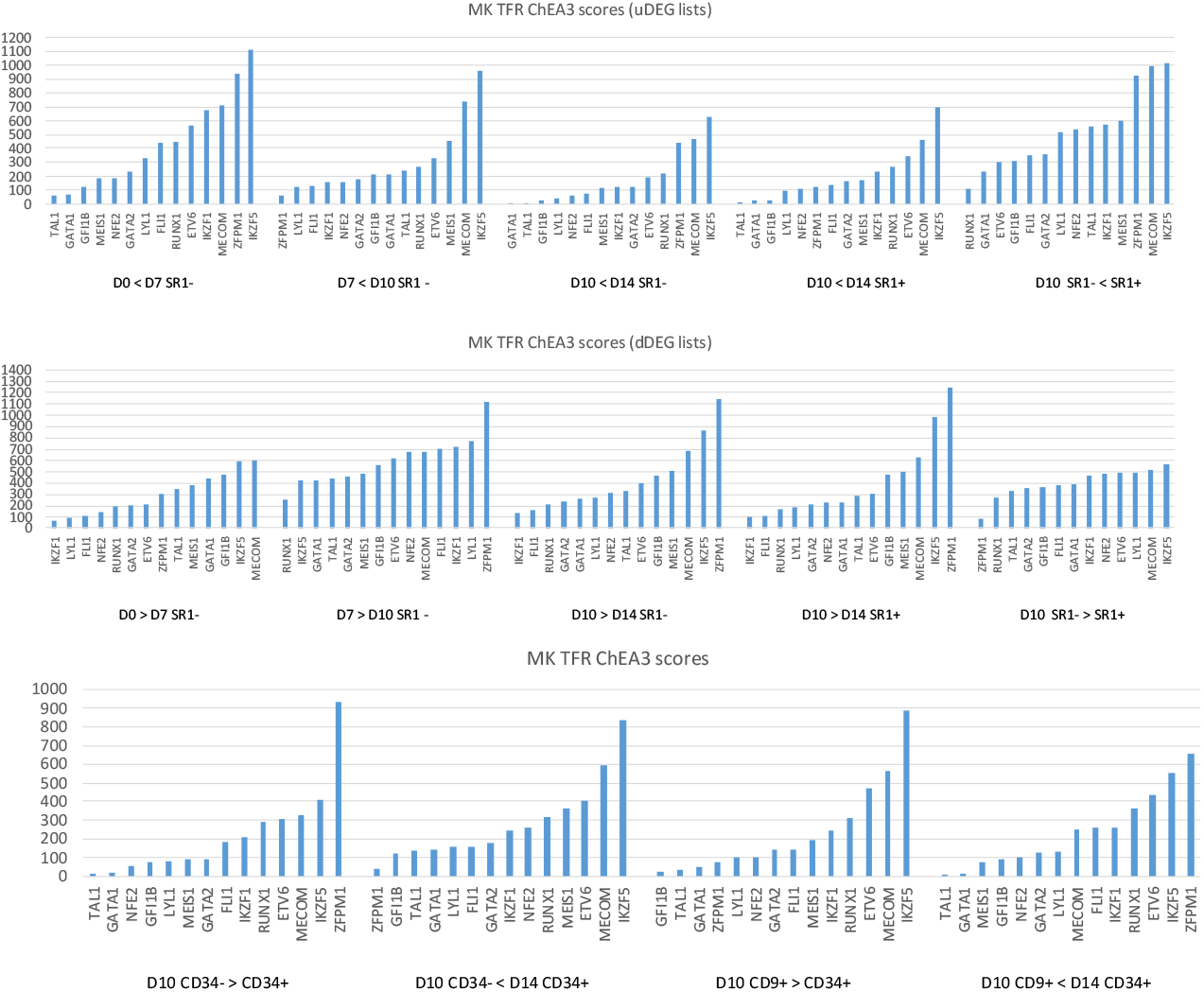
Scores of MK TFRs (mean rank table). DEG lists corresponding to pairs of conditions were analyzed using ChEA3 algorithm. The scores of MK TFR for DEGs between pairs of conditions enriched in megakaryocyte or platelet GSEA attributes are represented (other conditions are provided in supplemental data). Top (uDEGs) and middle (dDEGs) histograms, D0, D7, and D10 or D14 correspond to D0 CD34+, D17 CD34+CD41+ and D10 or D14 CD34+CD41+CD9-cells, respectively; SR1- and SR1+ refers to the analyzed culture condition. Bottom histogram, CD34- and CD9+ stands for CD34-CD34+ and CD34+CD41+CD9+ cells, CD34+ for CD34+CD41+CD9-cells, all generated in the presence of SR1. < and > symbols represent the down- or up-regulation tendency of the differential expression in the analyzed lists.

**Supplemental Figure 7.**
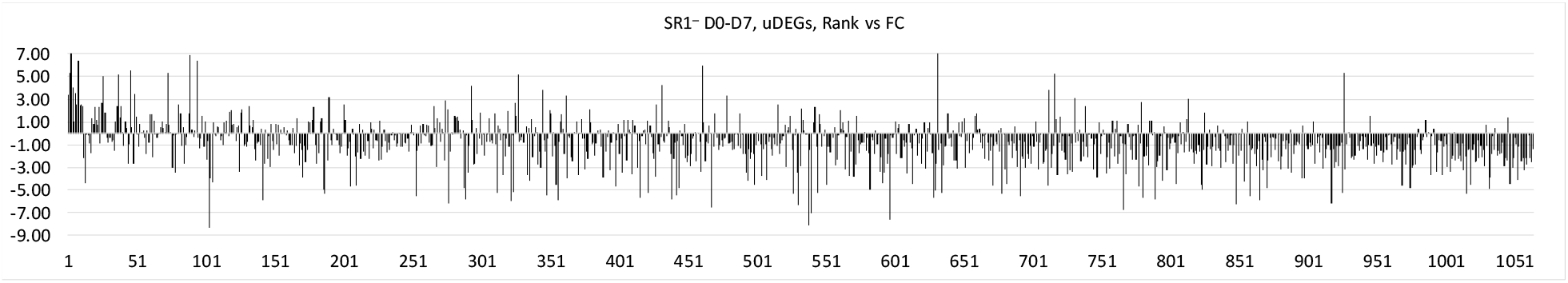
CHEA3 analysis of uDEGs during D0-D7 step. The D0-D7 Log_2_ FCs of the 1064 expressed TFRs (arbitrarily defined by mean expression value>5 for at least one of the D0 and D7 condition) are plotted versus the increasing mean ranks.

**Supplemental Figure 8.**
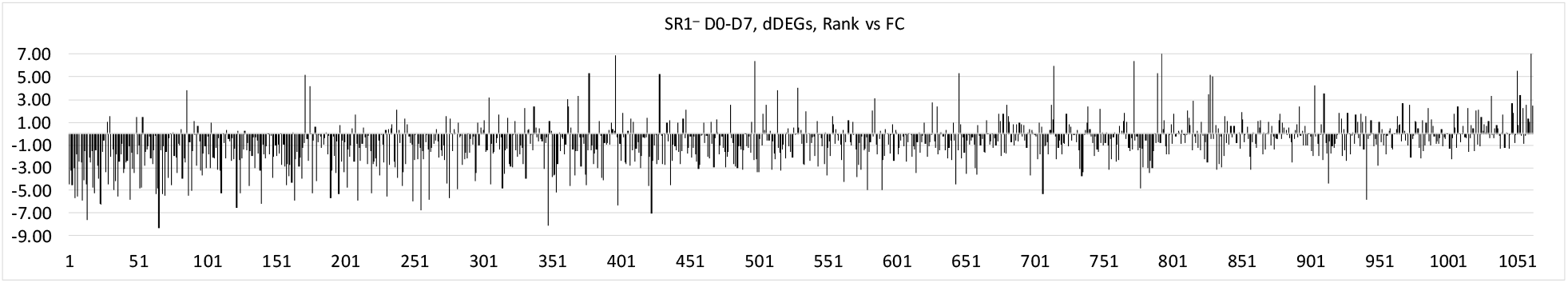
CHEA3 analysis of dDEGs during D0-D7 step. The D0-D7 Log_2_ FCs of the 1064 expressed TFRs (arbitrarily defined by mean expression value>5 for at least one of the D0 and D7 condition) are plotted versus the increasing mean ranks.

**Supplemental Figure 9.**
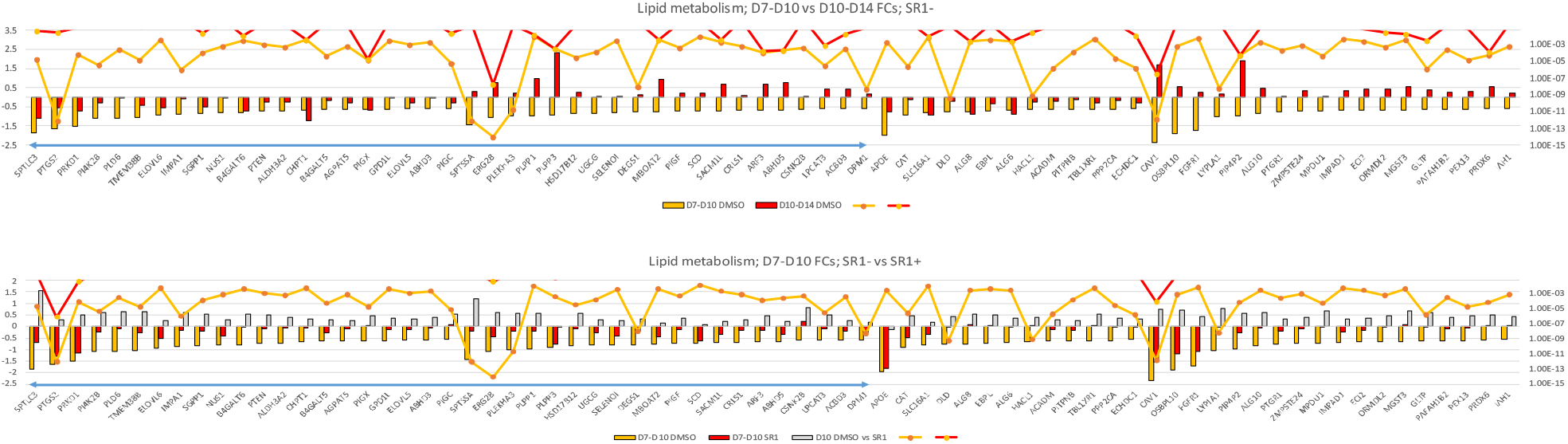
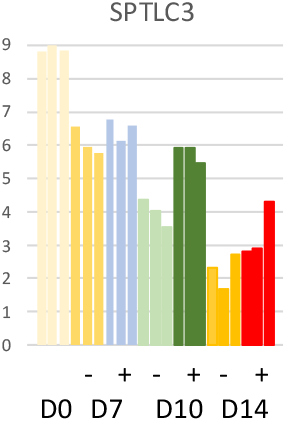
Differential expression of genes participating to lipid biosynthetic process (GO:0008610, underlined by the bidirectional blue arrow) or other lipid metabolic process (GO:0006629). Top histogram compares the FCs of the DEGs during D7-D10 (orange) and D10-D14 (red) periods in the SR1^-^ condition. Genes were first grouped according to negative or positive regulation during the two periods, then according decreasing D7-D10 |FCs|. Bottom diagram compares the FCs during D7-D10 period under SR1^-^ (orange) and SR1^+^ (red) conditions. ApVals of the FCs under the different conditions are depicted by the colored lines (only ApVal<0.05 are represented). Right, gene expression profile of SPTLC3 between D0 and D14 in SR1^+^ and SR1^-^ conditions.

**Supplemental Figure 10.**
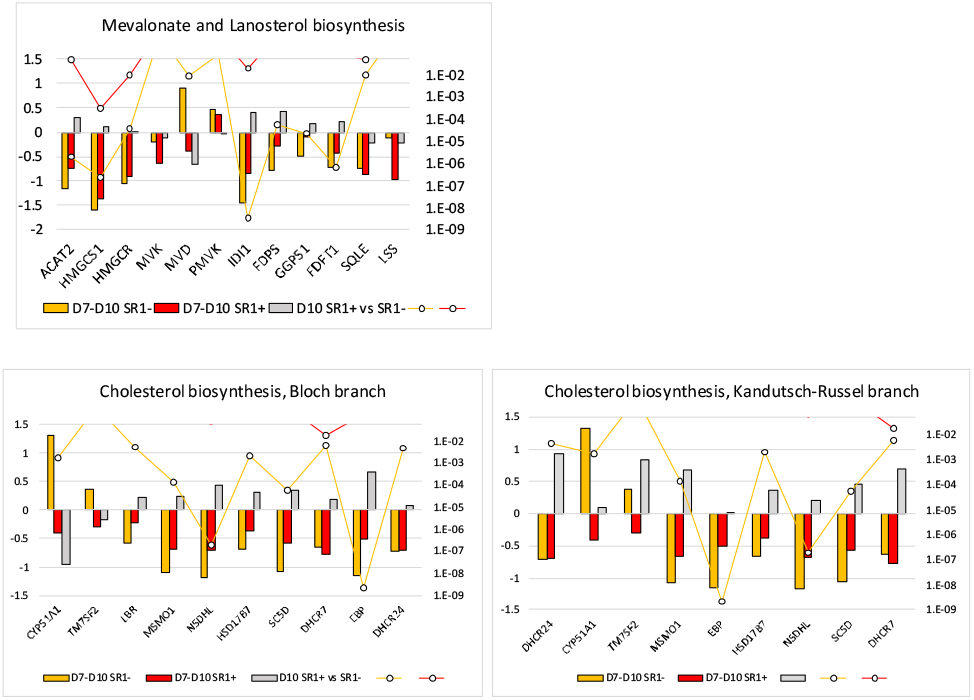
Comparison of the D7-D10 log2 FCs of the genes participating to the cholesterol biosynthetic pathways, in the SR1^-^ (orange) or SR1^+^ (red), or between SR1^+^ and the SR1^-^ at D10 (grey). The ApVal of the FCs in the SR1^-^ and SR1^+^ conditions below 0.05 are represented by the orange and red lines, respectively. Only EBP was differentially expressed at D10 between SR1^-^ and SR1^+^ conditions (ApVal = 4.23 E-03). The genes are ordered according to their sequence in the biosynthetic pathways.

**Supplemental Figure 11.**
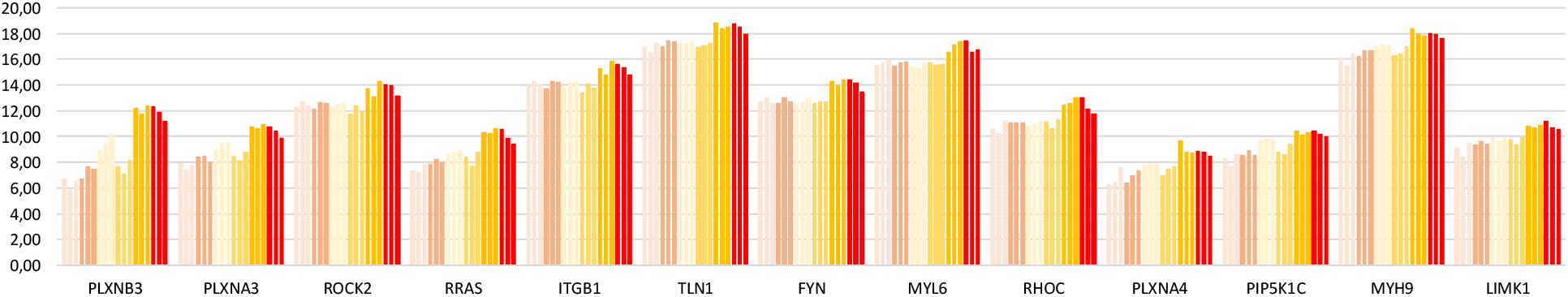
D7-D10 expression profile of uDEGs belonging to the gene list associated to Reactome pathway “semaphorin interactions” (R-HSA-373755). The triplicated normalized expression values are represented; from left to right D7 SR1^-^ and SR1^+^, D10 SR1^-^ and SR1^+^, D14 SR1^-^ and SR1^+^. Genes are ordered according to decreasing FC for SR1+ D10-D14 condition.

**Supplemental Figure 12.**
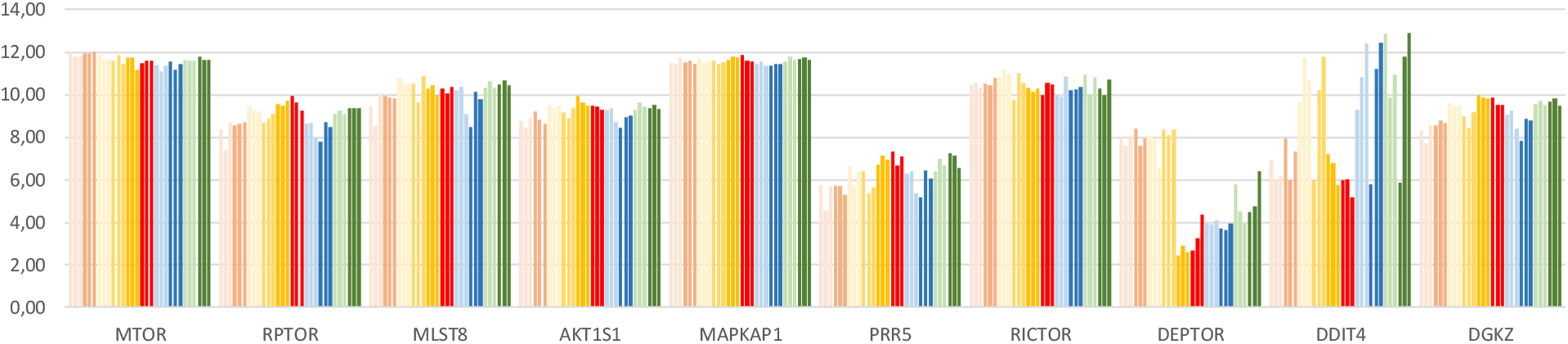
Expression profiles of mTOR complex-associated protein and negative regulators, and DGKZ in productive and unproductive cells. The triplicated normalized expression values are represented; from left to right D7MKp (orange), D10MKp, D14MKp, D10MKu^_34-_^ and D10MKu^_9+_^ cells in SR1^-^ (light colors) or SR1^+^ (deep colors).

**Supplemental Figure 13.**
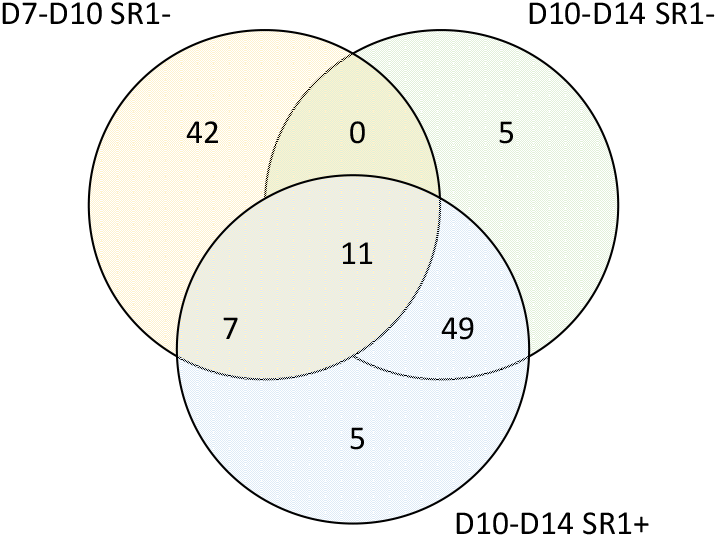
Venn diagram of DEGs participating to the regulation of GTPase activity (GO:0043087) and significantly upregulated between D7 and D14 (Figure 3a).

## Notes

### Competing Interest Statement

The authors have declared no competing interest.

