## Supplemental tables for "Transcriptome analysis of the effect of AHR on productive and unproductive pathways of *in vitro* megakaryocytopoiesis": Supplemental Figures.pptx

### Slide 1
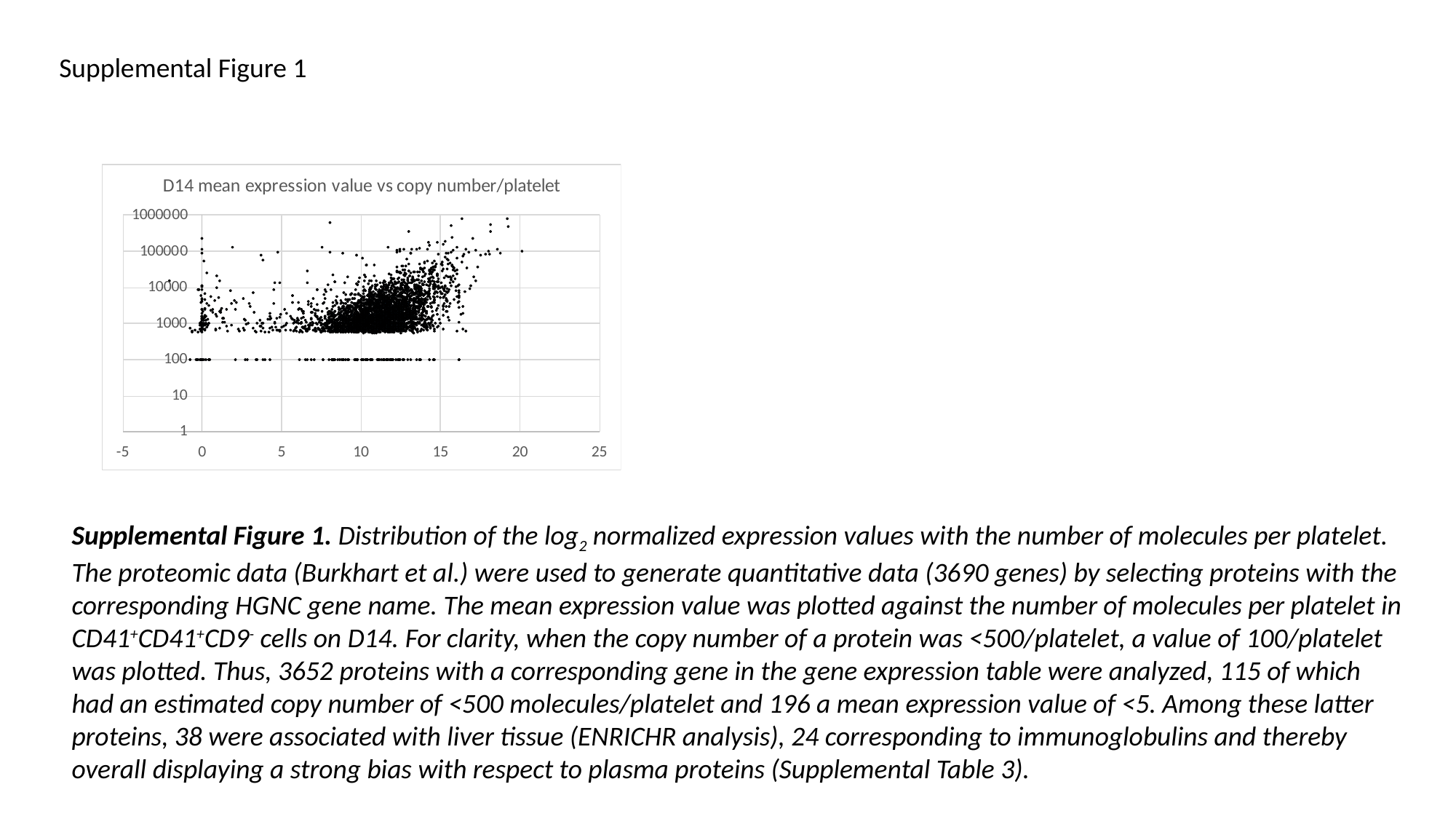

Supplemental Figure 1
Supplemental Figure 1. Distribution of the log_2_ normalized expression values with the number of molecules per platelet. The proteomic data (Burkhart et al.) were used to generate quantitative data (3690 genes) by selecting proteins with the corresponding HGNC gene name. The mean expression value was plotted against the number of molecules per platelet in CD41^+^CD41^+^CD9^-^ cells on D14. For clarity, when the copy number of a protein was <500/platelet, a value of 100/platelet was plotted. Thus, 3652 proteins with a corresponding gene in the gene expression table were analyzed, 115 of which had an estimated copy number of <500 molecules/platelet and 196 a mean expression value of <5. Among these latter proteins, 38 were associated with liver tissue (ENRICHR analysis), 24 corresponding to immunoglobulins and thereby overall displaying a strong bias with respect to plasma proteins (Supplemental Table 3).

### Slide 2
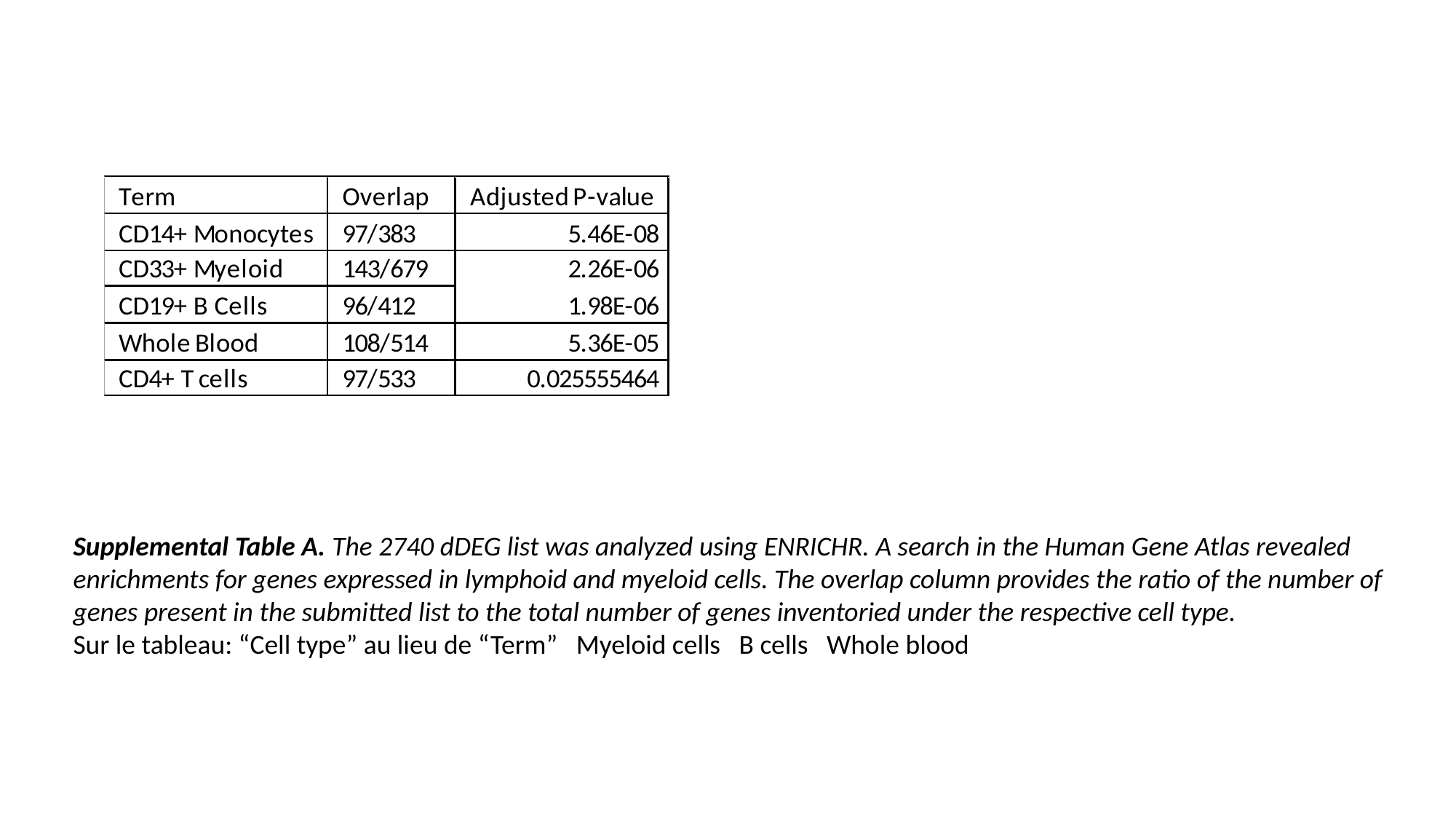

Supplemental Table A. The 2740 dDEG list was analyzed using ENRICHR. A search in the Human Gene Atlas revealed enrichments for genes expressed in lymphoid and myeloid cells. The overlap column provides the ratio of the number of genes present in the submitted list to the total number of genes inventoried under the respective cell type.
Sur le tableau: “Cell type” au lieu de “Term” Myeloid cells B cells Whole blood

### Slide 3
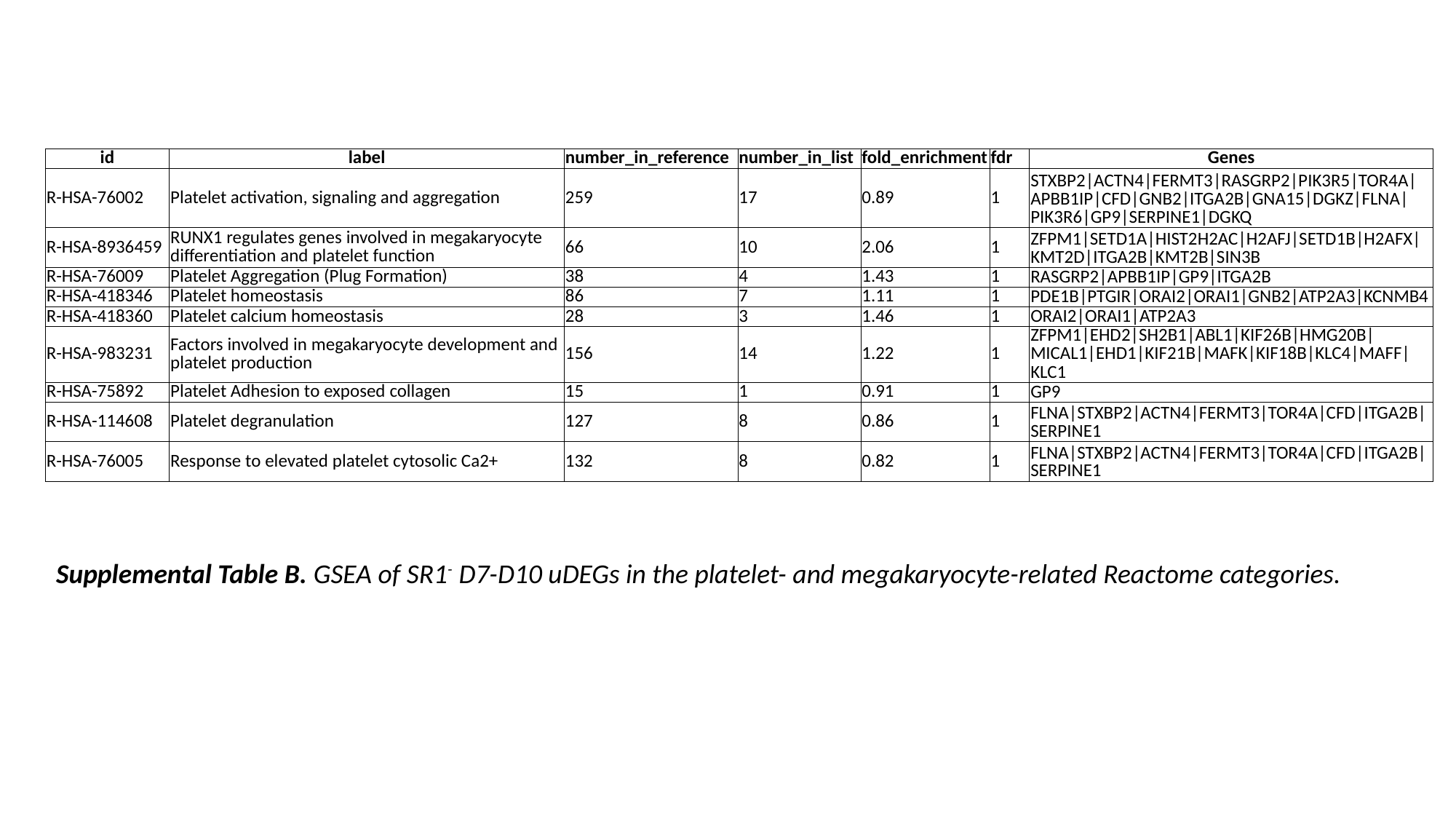

| id | label | number\_in\_reference | number\_in\_list | fold\_enrichment | fdr | Genes |
| --- | --- | --- | --- | --- | --- | --- |
| R-HSA-76002 | Platelet activation, signaling and aggregation | 259 | 17 | 0.89 | 1 | STXBP2|ACTN4|FERMT3|RASGRP2|PIK3R5|TOR4A|APBB1IP|CFD|GNB2|ITGA2B|GNA15|DGKZ|FLNA|PIK3R6|GP9|SERPINE1|DGKQ |
| R-HSA-8936459 | RUNX1 regulates genes involved in megakaryocyte differentiation and platelet function | 66 | 10 | 2.06 | 1 | ZFPM1|SETD1A|HIST2H2AC|H2AFJ|SETD1B|H2AFX|KMT2D|ITGA2B|KMT2B|SIN3B |
| R-HSA-76009 | Platelet Aggregation (Plug Formation) | 38 | 4 | 1.43 | 1 | RASGRP2|APBB1IP|GP9|ITGA2B |
| R-HSA-418346 | Platelet homeostasis | 86 | 7 | 1.11 | 1 | PDE1B|PTGIR|ORAI2|ORAI1|GNB2|ATP2A3|KCNMB4 |
| R-HSA-418360 | Platelet calcium homeostasis | 28 | 3 | 1.46 | 1 | ORAI2|ORAI1|ATP2A3 |
| R-HSA-983231 | Factors involved in megakaryocyte development and platelet production | 156 | 14 | 1.22 | 1 | ZFPM1|EHD2|SH2B1|ABL1|KIF26B|HMG20B|MICAL1|EHD1|KIF21B|MAFK|KIF18B|KLC4|MAFF|KLC1 |
| R-HSA-75892 | Platelet Adhesion to exposed collagen | 15 | 1 | 0.91 | 1 | GP9 |
| R-HSA-114608 | Platelet degranulation | 127 | 8 | 0.86 | 1 | FLNA|STXBP2|ACTN4|FERMT3|TOR4A|CFD|ITGA2B|SERPINE1 |
| R-HSA-76005 | Response to elevated platelet cytosolic Ca2+ | 132 | 8 | 0.82 | 1 | FLNA|STXBP2|ACTN4|FERMT3|TOR4A|CFD|ITGA2B|SERPINE1 |
Supplemental Table B. GSEA of SR1^-^ D7-D10 uDEGs in the platelet- and megakaryocyte-related Reactome categories.

### Slide 4
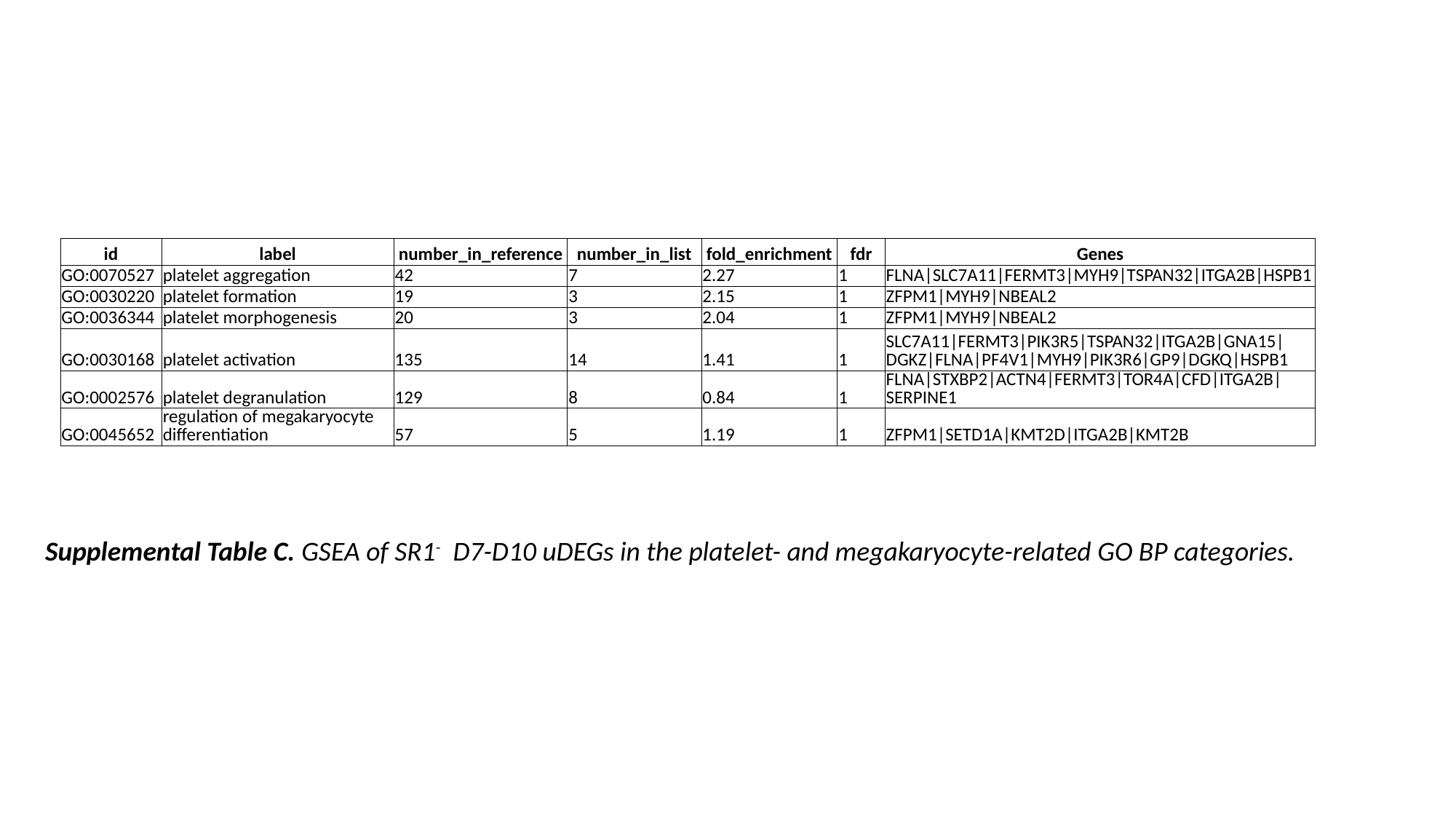

| id | label | number\_in\_reference | number\_in\_list | fold\_enrichment | fdr | Genes |
| --- | --- | --- | --- | --- | --- | --- |
| GO:0070527 | platelet aggregation | 42 | 7 | 2.27 | 1 | FLNA|SLC7A11|FERMT3|MYH9|TSPAN32|ITGA2B|HSPB1 |
| GO:0030220 | platelet formation | 19 | 3 | 2.15 | 1 | ZFPM1|MYH9|NBEAL2 |
| GO:0036344 | platelet morphogenesis | 20 | 3 | 2.04 | 1 | ZFPM1|MYH9|NBEAL2 |
| GO:0030168 | platelet activation | 135 | 14 | 1.41 | 1 | SLC7A11|FERMT3|PIK3R5|TSPAN32|ITGA2B|GNA15|DGKZ|FLNA|PF4V1|MYH9|PIK3R6|GP9|DGKQ|HSPB1 |
| GO:0002576 | platelet degranulation | 129 | 8 | 0.84 | 1 | FLNA|STXBP2|ACTN4|FERMT3|TOR4A|CFD|ITGA2B|SERPINE1 |
| GO:0045652 | regulation of megakaryocyte differentiation | 57 | 5 | 1.19 | 1 | ZFPM1|SETD1A|KMT2D|ITGA2B|KMT2B |
Supplemental Table C. GSEA of SR1^-^ D7-D10 uDEGs in the platelet- and megakaryocyte-related GO BP categories.

### Slide 5
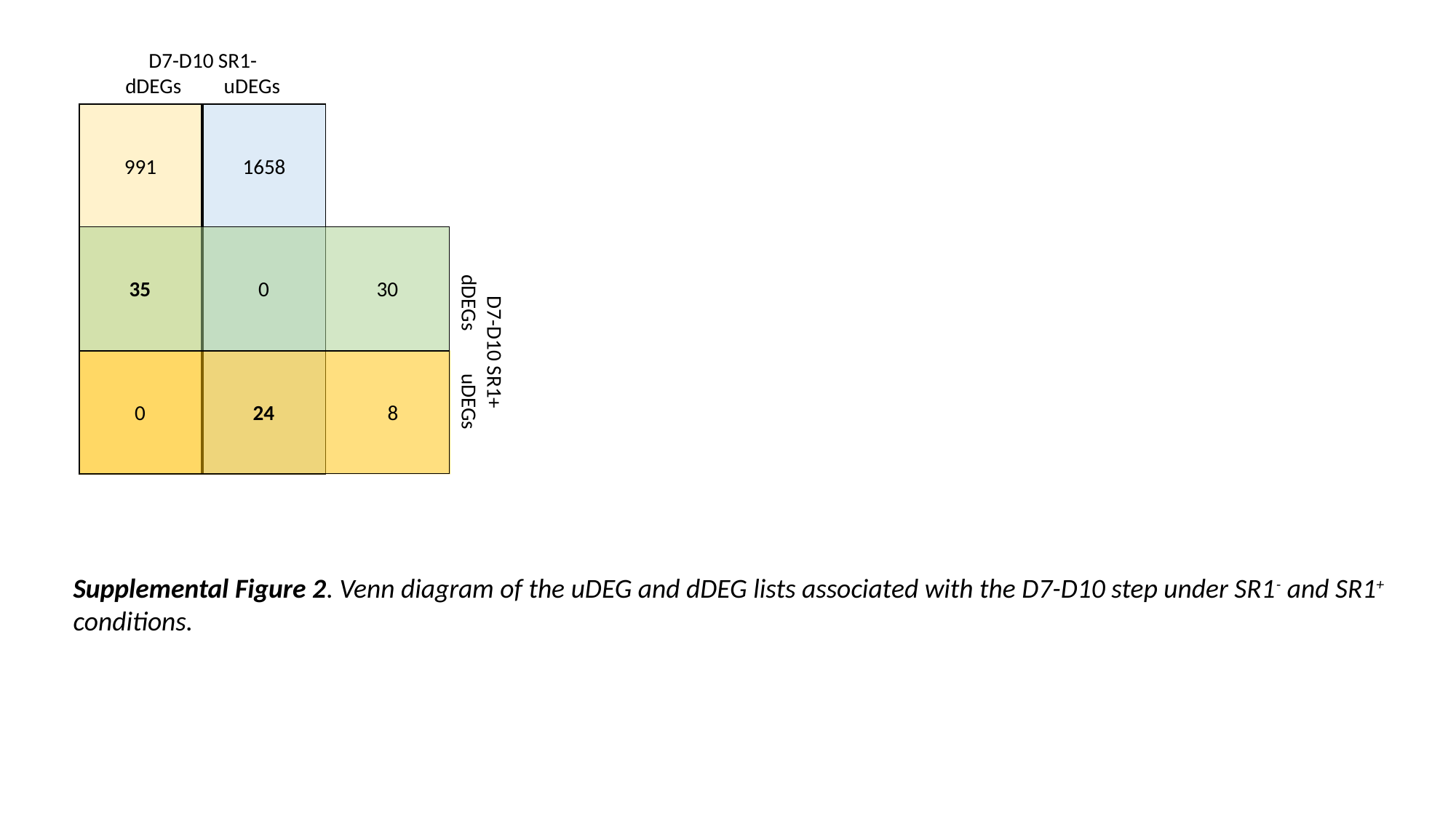

D7-D10 SR1-
dDEGs uDEGs
991
1658
35
0
30
D7-D10 SR1+
dDEGs uDEGs
0
24
8
Supplemental Figure 2. Venn diagram of the uDEG and dDEG lists associated with the D7-D10 step under SR1^-^ and SR1^+^ conditions.

### Slide 6
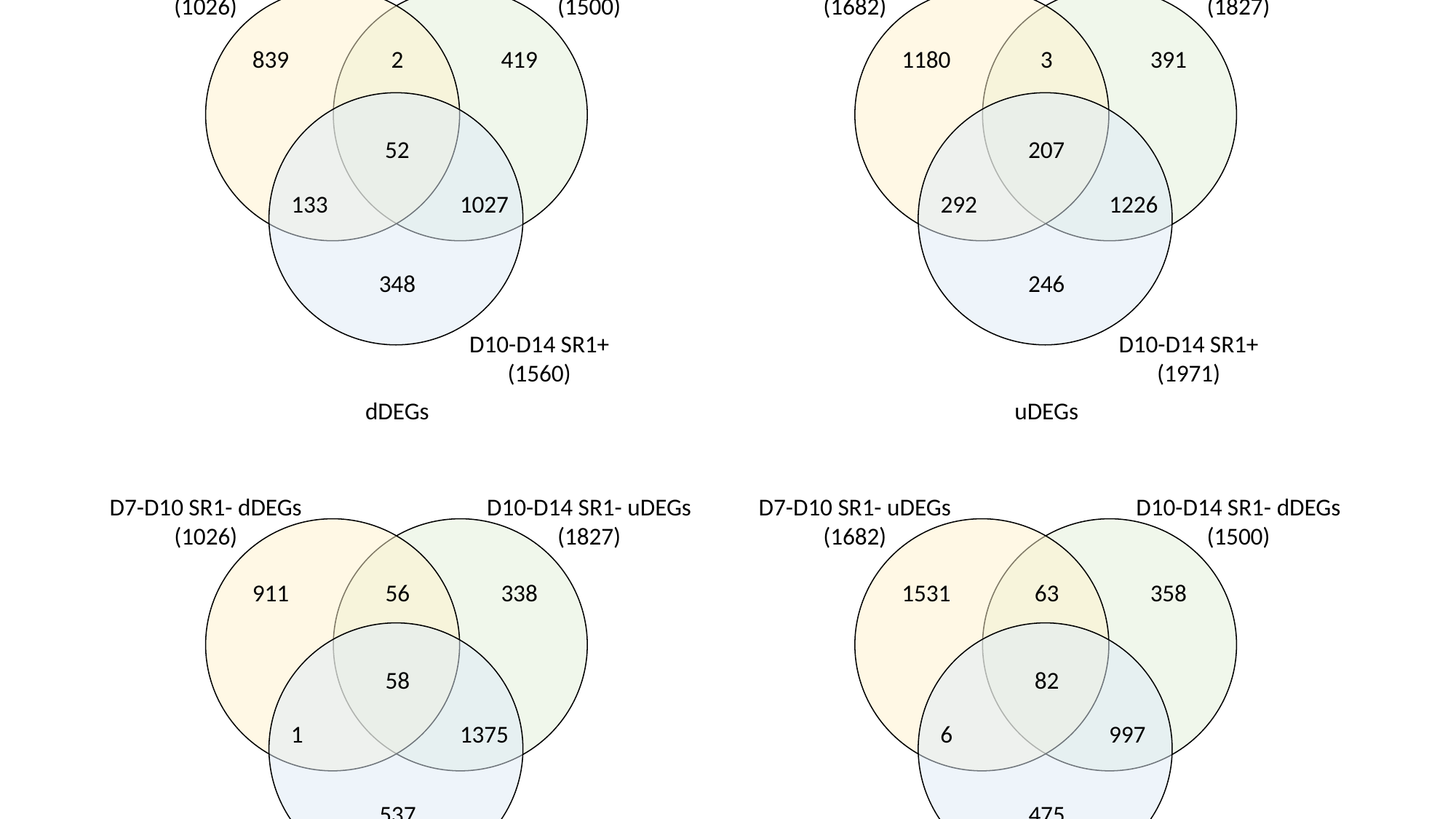

D7-D10 SR1-
(1026)
D10-D14 SR1-
(1500)
D7-D10 SR1-
(1682)
D10-D14 SR1-
(1827)
839
2
419
1180
3
391
52
207
133
1027
292
1226
348
246
D10-D14 SR1+
(1560)
D10-D14 SR1+
(1971)
dDEGs
uDEGs
D7-D10 SR1- dDEGs
(1026)
D10-D14 SR1- uDEGs
(1827)
D7-D10 SR1- uDEGs
(1682)
D10-D14 SR1- dDEGs
(1500)
911
56
338
1531
63
358
58
82
1
1375
6
997
537
475
D10-D14 SR1+ uDEGs
(1971)
D10-D14 SR1+ dDEGs
(1560)
dDEGs vs uDEGs
uDEGs vs dDEGs

### Slide 7
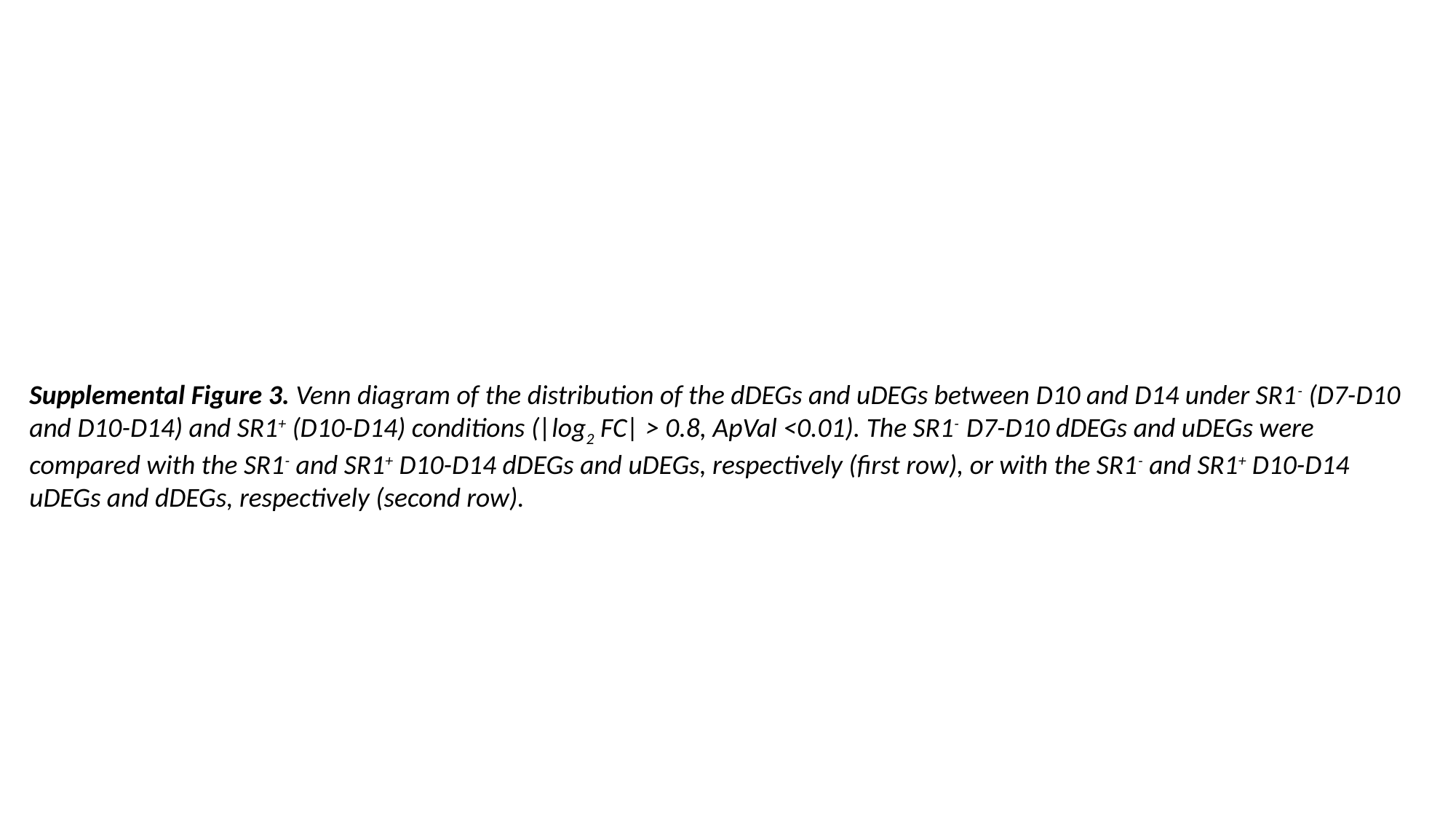

Supplemental Figure 3. Venn diagram of the distribution of the dDEGs and uDEGs between D10 and D14 under SR1^-^ (D7-D10 and D10-D14) and SR1^+^ (D10-D14) conditions (|log_2_ FC| > 0.8, ApVal <0.01). The SR1^-^ D7-D10 dDEGs and uDEGs were compared with the SR1^-^ and SR1^+^ D10-D14 dDEGs and uDEGs, respectively (first row), or with the SR1^-^ and SR1^+^ D10-D14 uDEGs and dDEGs, respectively (second row).

### Slide 8
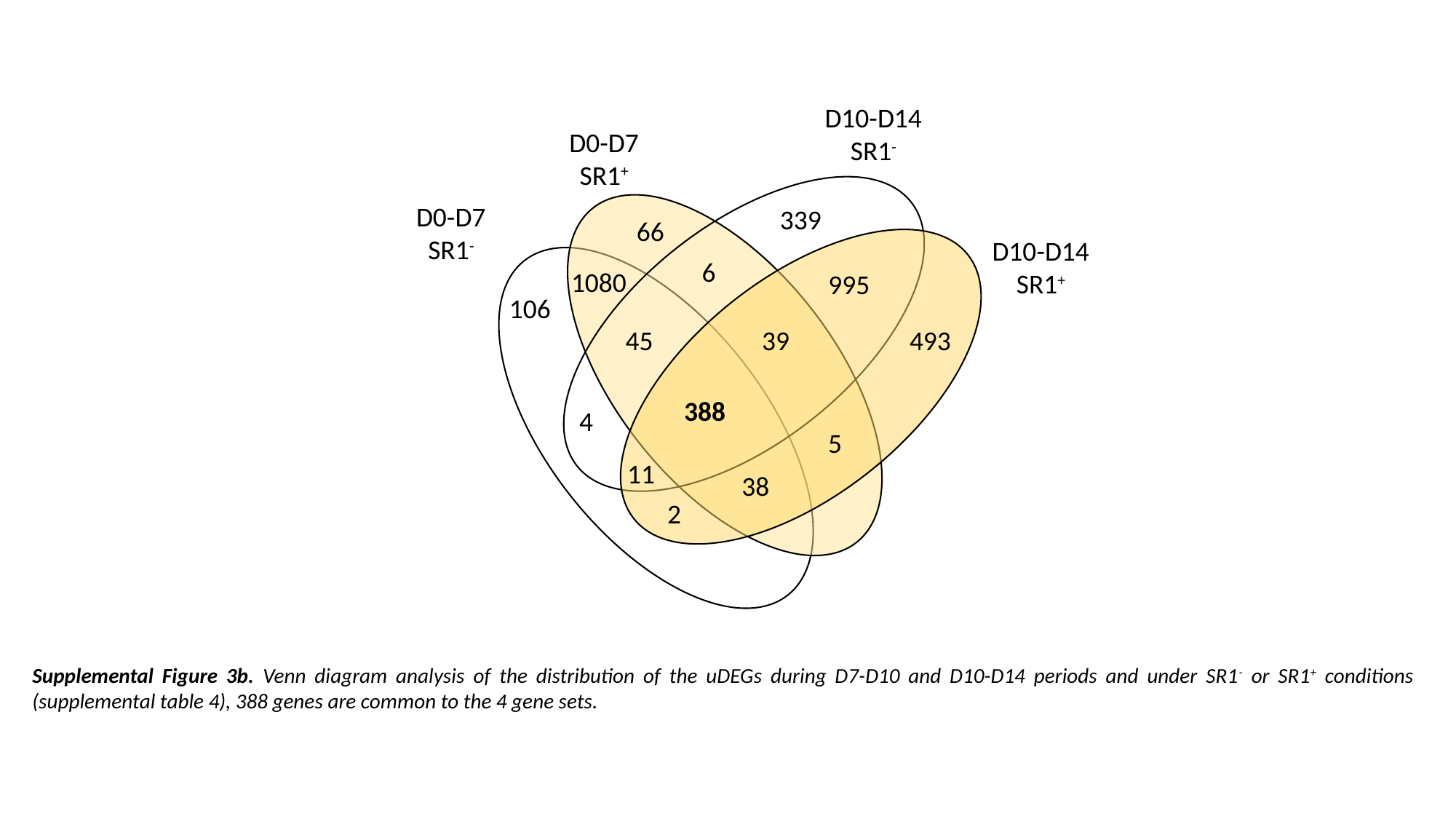

D10-D14
SR1^-^
D0-D7
SR1^+^
D0-D7
SR1^-^
339
66
D10-D14
SR1^+^
6
1080
995
106
45
39
493
388
4
5
11
38
2
Supplemental Figure 3b. Venn diagram analysis of the distribution of the uDEGs during D7-D10 and D10-D14 periods and under SR1^-^ or SR1^+^ conditions (supplemental table 4), 388 genes are common to the 4 gene sets.

### Slide 9
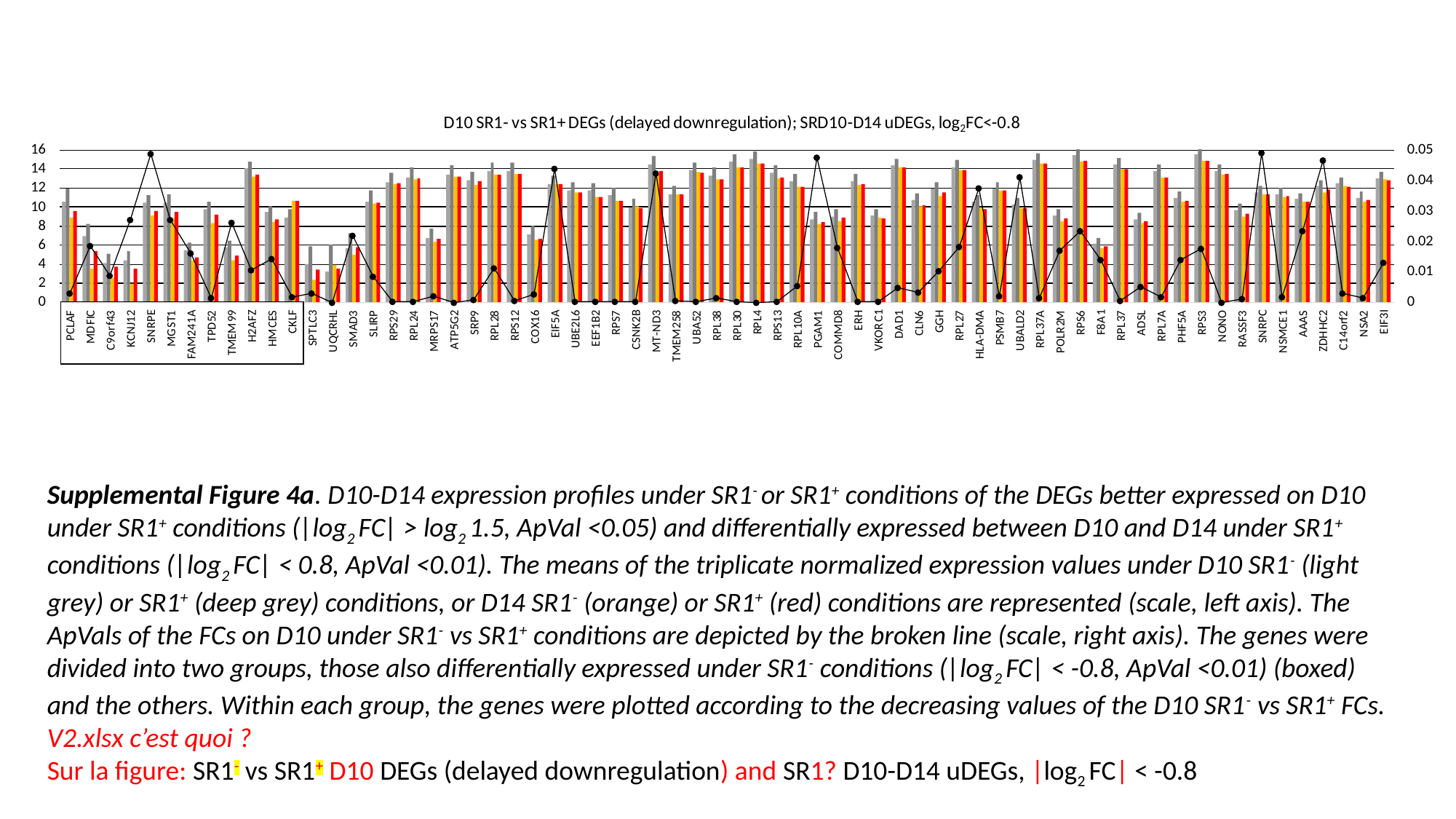

Supplemental Figure 4a. D10-D14 expression profiles under SR1^-^ or SR1^+^ conditions of the DEGs better expressed on D10 under SR1^+^ conditions (|log_2_ FC| > log_2_ 1.5, ApVal <0.05) and differentially expressed between D10 and D14 under SR1^+^ conditions (|log_2_ FC| < 0.8, ApVal <0.01). The means of the triplicate normalized expression values under D10 SR1^-^ (light grey) or SR1^+^ (deep grey) conditions, or D14 SR1^-^ (orange) or SR1^+^ (red) conditions are represented (scale, left axis). The ApVals of the FCs on D10 under SR1^-^ vs SR1^+^ conditions are depicted by the broken line (scale, right axis). The genes were divided into two groups, those also differentially expressed under SR1^-^ conditions (|log_2_ FC| < -0.8, ApVal <0.01) (boxed) and the others. Within each group, the genes were plotted according to the decreasing values of the D10 SR1^-^ vs SR1^+^ FCs. V2.xlsx c’est quoi ?
Sur la figure: SR1^-^ vs SR1^+^ D10 DEGs (delayed downregulation) and SR1? D10-D14 uDEGs, |log_2_ FC| < -0.8

### Slide 10
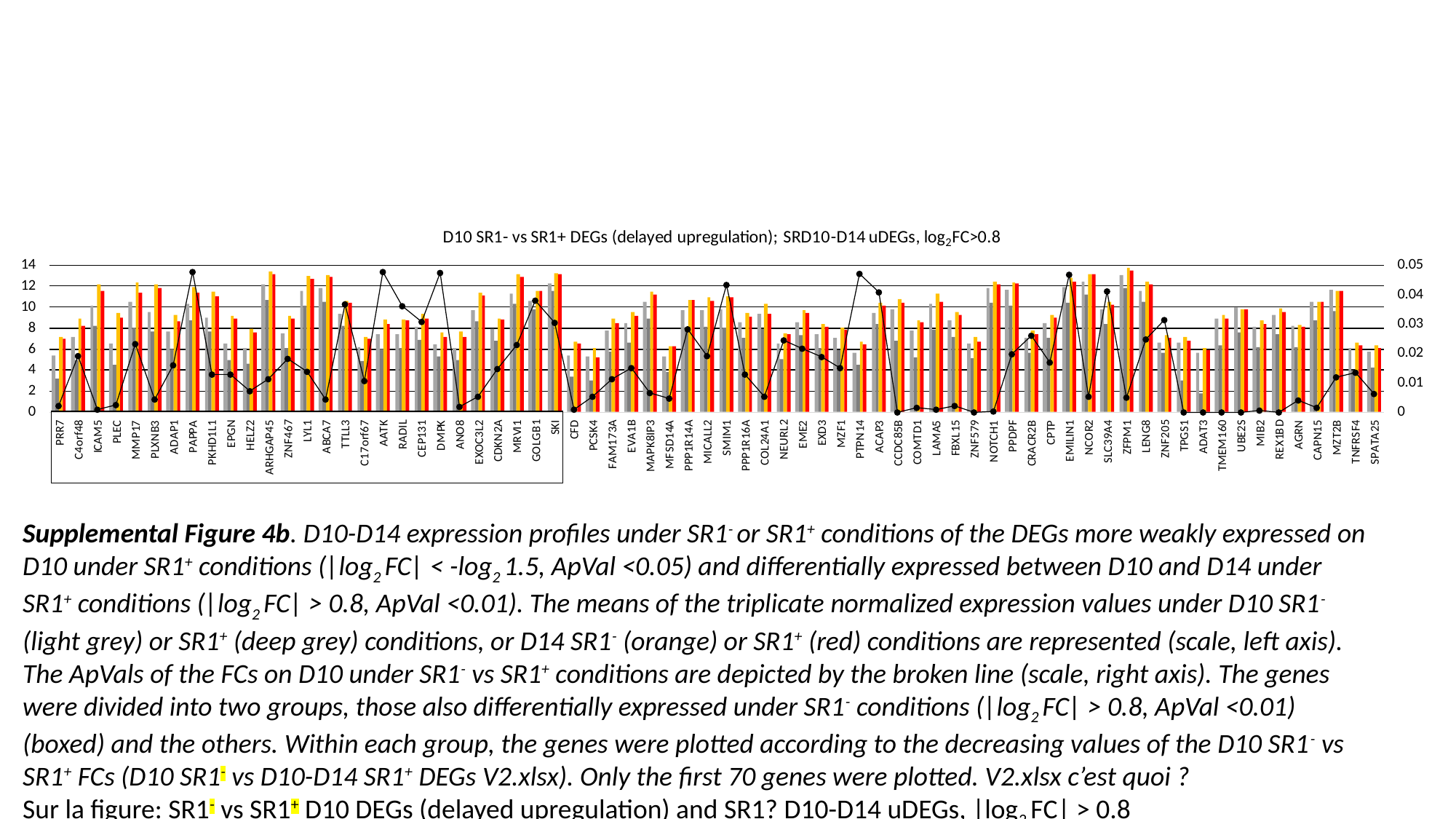

Supplemental Figure 4b. D10-D14 expression profiles under SR1^-^ or SR1^+^ conditions of the DEGs more weakly expressed on D10 under SR1^+^ conditions (|log_2_ FC| < -log_2_ 1.5, ApVal <0.05) and differentially expressed between D10 and D14 under SR1^+^ conditions (|log_2_ FC| > 0.8, ApVal <0.01). The means of the triplicate normalized expression values under D10 SR1^-^ (light grey) or SR1^+^ (deep grey) conditions, or D14 SR1^-^ (orange) or SR1^+^ (red) conditions are represented (scale, left axis). The ApVals of the FCs on D10 under SR1^-^ vs SR1^+^ conditions are depicted by the broken line (scale, right axis). The genes were divided into two groups, those also differentially expressed under SR1^-^ conditions (|log_2_ FC| > 0.8, ApVal <0.01) (boxed) and the others. Within each group, the genes were plotted according to the decreasing values of the D10 SR1^-^ vs SR1^+^ FCs (D10 SR1^-^ vs D10-D14 SR1^+^ DEGs V2.xlsx). Only the first 70 genes were plotted. V2.xlsx c’est quoi ?
Sur la figure: SR1^-^ vs SR1^+^ D10 DEGs (delayed upregulation) and SR1? D10-D14 uDEGs, |log_2_ FC| > 0.8

### Slide 11
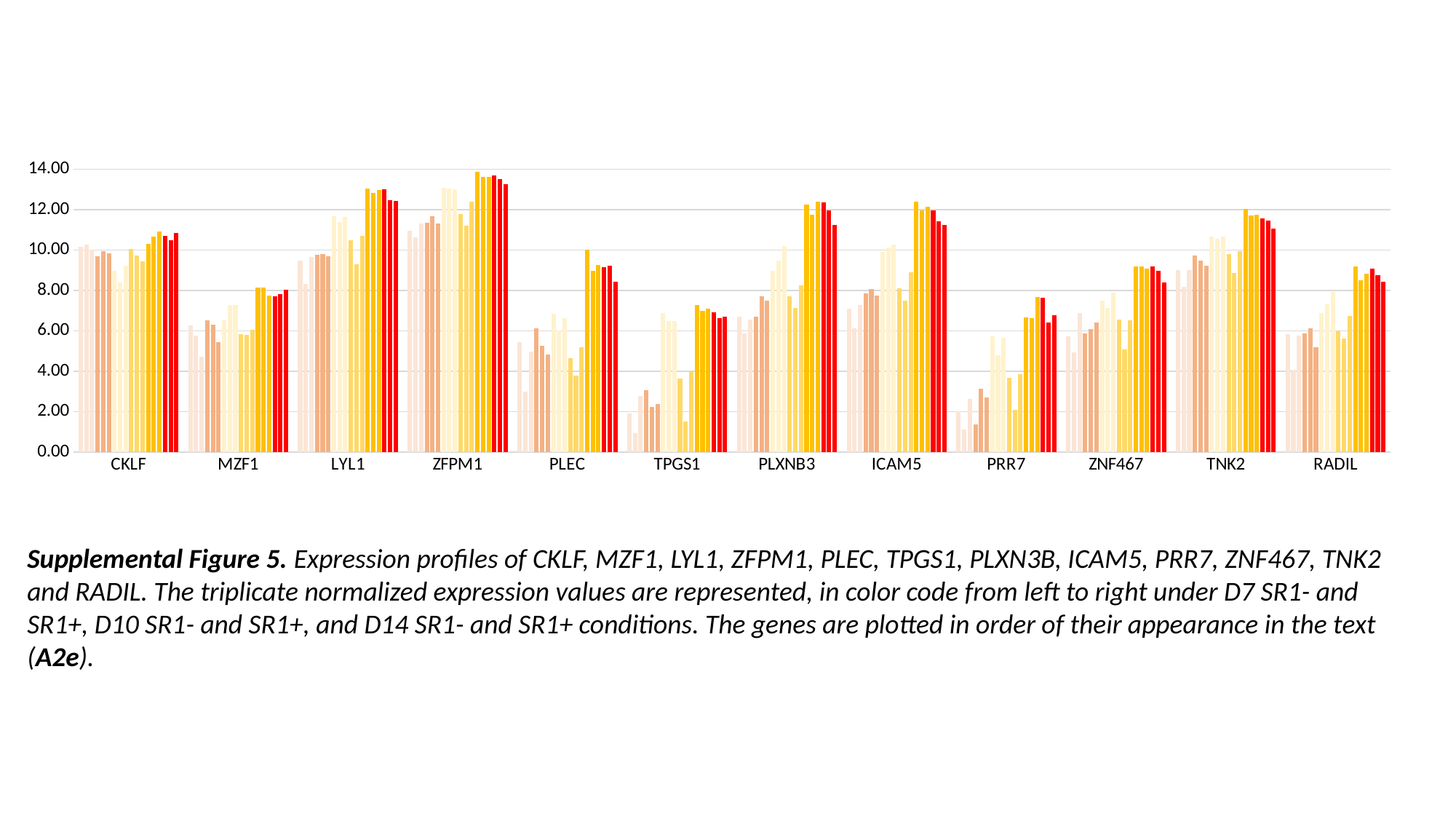

#### Chart
| Category | CNSL1 | CNSL2 | CNSL3 | CNSL4 | CNSL5 | CNSL6 | VNDT4 | VNDT5 | VNDT6 | CNSL13 | CNSL14 | CNSL15 | VNDT13 | VNDT14 | VNDT15 | VNDT19 | VNDT20 | VNDT21 |
|---|---|---|---|---|---|---|---|---|---|---|---|---|---|---|---|---|---|---|
| CKLF | 10.150734603541588 | 10.27438892080295 | 10.022624996753347 | 9.713056136094059 | 9.960411578860036 | 9.828399617592057 | 8.964965338366342 | 8.417168436423143 | 9.221274191385387 | 10.066233212880292 | 9.744206473715238 | 9.459068366524452 | 10.299690001568496 | 10.671234633151258 | 10.91245614719609 | 10.700278853986136 | 10.493873246230562 | 10.832353875885556 |
| MZF1 | 6.270568012614466 | 5.763017583541499 | 4.707580722206286 | 6.528352345616749 | 6.317152017925406 | 5.44817446249122 | 6.507784999833562 | 7.289525726338847 | 7.297958364291122 | 5.841190395478585 | 5.800472621097428 | 6.047390675345824 | 8.15620168903009 | 8.153871937996897 | 7.75473480662676 | 7.717047263258273 | 7.813841832712292 | 8.024461834051039 |
| LYL1 | 9.463432401621096 | 8.329064353239694 | 9.652620703786672 | 9.7779797821536 | 9.79768480552086 | 9.69476117828114 | 11.663177673405434 | 11.385045442952961 | 11.633343368271605 | 10.500354955745532 | 9.31659416546394 | 10.69034271581018 | 13.042155372334875 | 12.84991713983184 | 12.969686816272706 | 13.027366826478682 | 12.459571288753418 | 12.44966307149005 |
| ZFPM1 | 10.95773571386197 | 10.626254652592479 | 11.317805741261537 | 11.356045800100363 | 11.68397066493974 | 11.32014634151981 | 13.07052664710763 | 13.065341819220475 | 13.015486669320612 | 11.78211964690544 | 11.224209946329136 | 12.389744998959296 | 13.874294610838392 | 13.64230087112354 | 13.616966303014953 | 13.687031615019627 | 13.505022894550871 | 13.271697527811465 |
| PLEC | 5.4505183654065155 | 2.997176469926232 | 4.9614704480900205 | 6.109601509749886 | 5.277328942592255 | 4.838035662401311 | 6.849578896629947 | 6.011965686594166 | 6.613818982757438 | 4.654140758640722 | 3.7836587632123804 | 5.191340907941992 | 10.003770851992426 | 8.990494890259143 | 9.27728108618212 | 9.16997837878017 | 9.22988045954252 | 8.449531029299282 |
| TPGS1 | 1.8965997386440197 | 0.9317931550919683 | 2.791638159624415 | 3.048133244919008 | 2.2178221940946634 | 2.3932387580189305 | 6.874060822337087 | 6.477565321321837 | 6.469639667017006 | 3.64513991370398 | 1.5042854680373714 | 3.978183213117868 | 7.271124845253237 | 6.993290799764531 | 7.084284519830934 | 6.935458469523216 | 6.644010346845682 | 6.714078676797434 |
| PLXNB3 | 6.714250517386525 | 5.866483355024694 | 6.5544733310164265 | 6.719789039017936 | 7.704308030813346 | 7.496528561927403 | 8.958678558661132 | 9.47089451968325 | 10.212576717963646 | 7.7308960878277055 | 7.136048535012794 | 8.240038156304102 | 12.245479262759478 | 11.750474678247654 | 12.393332190796384 | 12.356248500078395 | 11.954196732158604 | 11.23508284831445 |
| ICAM5 | 7.1112149487307645 | 6.122362891212069 | 7.291097990508095 | 7.851679857929858 | 8.089113413741348 | 7.7511595519175165 | 9.927679534387725 | 10.144470106914708 | 10.261246666716335 | 8.09570999556824 | 7.4981632931523725 | 8.909109132604947 | 12.393270285204556 | 11.979938718046903 | 12.156861941815054 | 11.952350573079904 | 11.414573556990819 | 11.234777130589483 |
| PRR7 | 2.0114137172914437 | 1.1169964616307946 | 2.626344219079105 | 1.38533964490288 | 3.139991094799691 | 2.7039729466416125 | 5.748476591593743 | 4.783623022735046 | 5.654842973555827 | 3.6859533109076277 | 2.1045262738550377 | 3.841458424769333 | 6.648973229734456 | 6.63417359007954 | 7.662386348379355 | 7.64384807919651 | 6.417969727264108 | 6.7912222374788085 |
| ZNF467 | 5.717541522125675 | 4.941180027986737 | 6.889200843905546 | 5.884371215497174 | 6.106354672115042 | 6.402625630913696 | 7.502825606085463 | 7.1275230651967085 | 7.890378992382759 | 6.547611582682236 | 5.098634841653293 | 6.530712411039164 | 9.187833119027497 | 9.198264025913812 | 9.075775639223792 | 9.204997239718656 | 8.976692150391717 | 8.416860124452674 |
| TNK2 | 9.011813893979369 | 8.194505564016488 | 9.029851684962892 | 9.740557015593051 | 9.472467321751061 | 9.212520026999865 | 10.665846173346594 | 10.566337780764052 | 10.661962748050128 | 9.812033246934684 | 8.882062483184706 | 9.943781075655048 | 12.02368612427764 | 11.7312148621292 | 11.734966184836123 | 11.582995293956948 | 11.464547946819232 | 11.082671196170125 |
| RADIL | 5.85077861059306 | 4.0731750199151575 | 5.76309502300529 | 5.883088108395022 | 6.127878512811395 | 5.202944551403193 | 6.896066627881398 | 7.30684217374331 | 7.915027718730292 | 5.979166856469567 | 5.6277454248633 | 6.7431891990313195 | 9.186199238736595 | 8.51996282047251 | 8.837411749806448 | 9.07779989210768 | 8.741947421186477 | 8.446104033647943 |Supplemental Figure 5. Expression profiles of CKLF, MZF1, LYL1, ZFPM1, PLEC, TPGS1, PLXN3B, ICAM5, PRR7, ZNF467, TNK2 and RADIL. The triplicate normalized expression values are represented, in color code from left to right under D7 SR1- and SR1+, D10 SR1- and SR1+, and D14 SR1- and SR1+ conditions. The genes are plotted in order of their appearance in the text (A2e).

### Slide 12
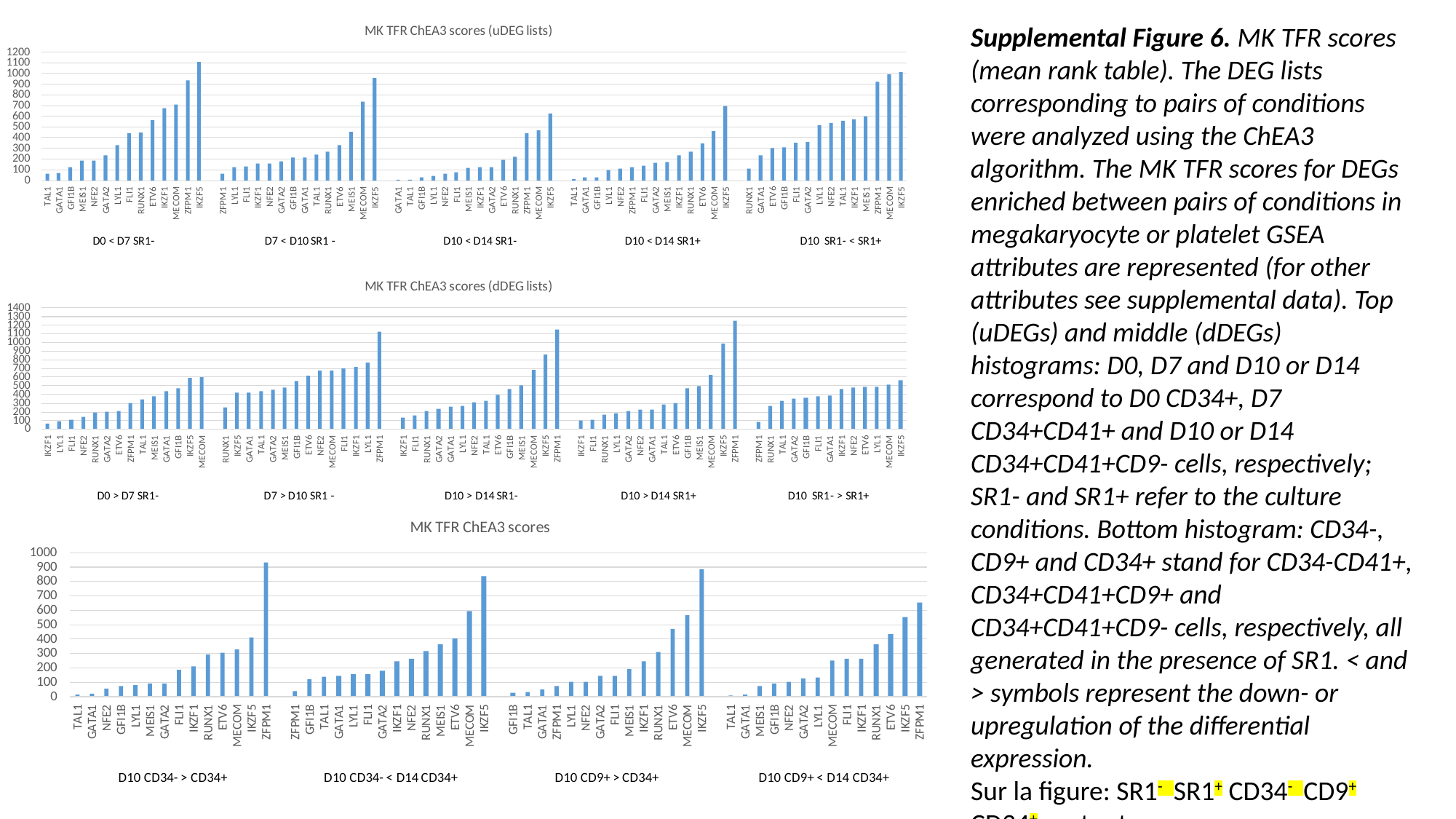

Supplemental Figure 6. MK TFR scores (mean rank table). The DEG lists corresponding to pairs of conditions were analyzed using the ChEA3 algorithm. The MK TFR scores for DEGs enriched between pairs of conditions in megakaryocyte or platelet GSEA attributes are represented (for other attributes see supplemental data). Top (uDEGs) and middle (dDEGs) histograms: D0, D7 and D10 or D14 correspond to D0 CD34+, D7 CD34+CD41+ and D10 or D14 CD34+CD41+CD9- cells, respectively; SR1- and SR1+ refer to the culture conditions. Bottom histogram: CD34-, CD9+ and CD34+ stand for CD34-CD41+, CD34+CD41+CD9+ and CD34+CD41+CD9- cells, respectively, all generated in the presence of SR1. < and > symbols represent the down- or upregulation of the differential expression.
Sur la figure: SR1^-^ SR1^+^ CD34^-^ CD9^+^ CD34^+^ partout.

### Slide 13
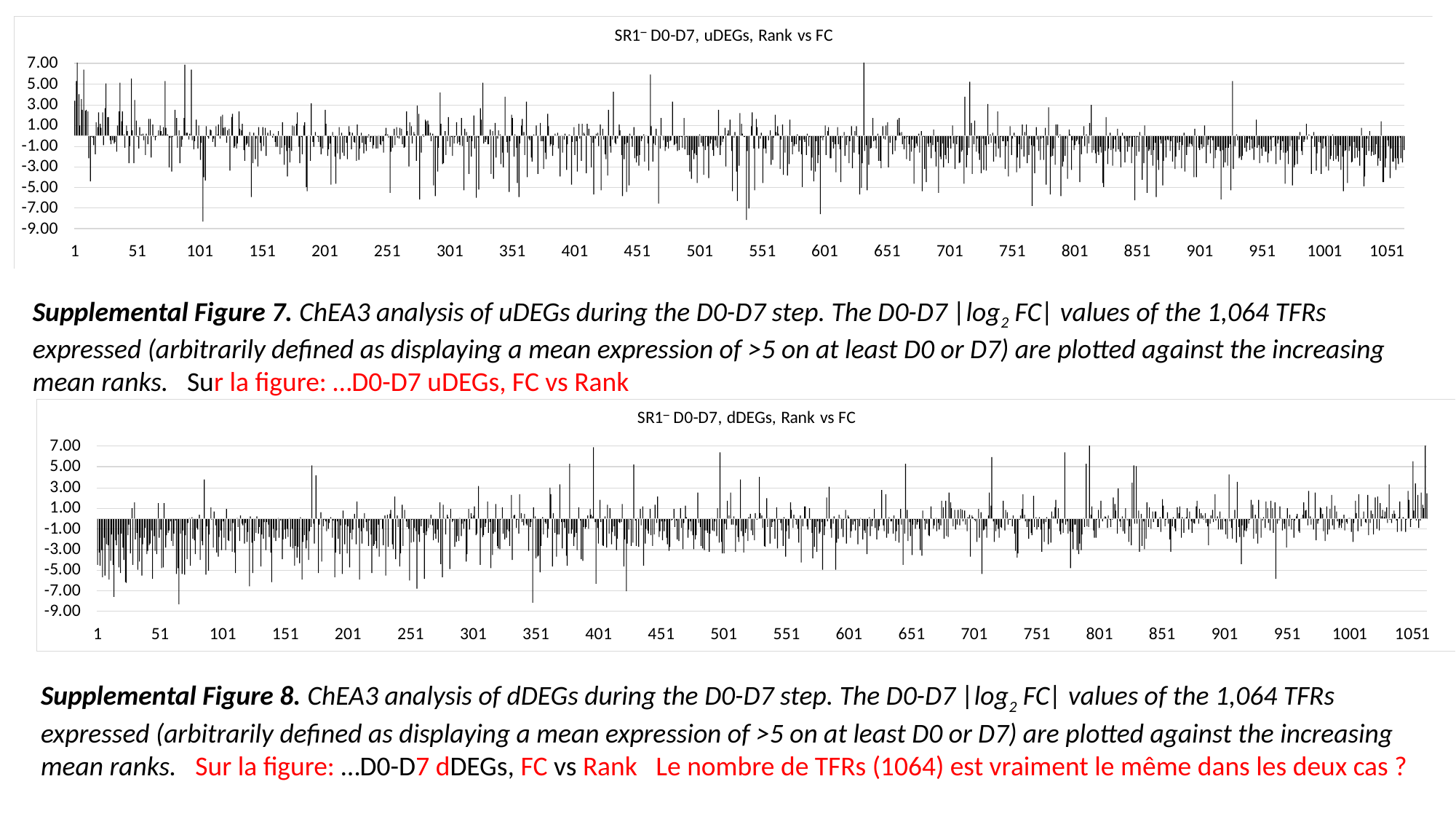

Supplemental Figure 7. ChEA3 analysis of uDEGs during the D0-D7 step. The D0-D7 |log_2_ FC| values of the 1,064 TFRs expressed (arbitrarily defined as displaying a mean expression of >5 on at least D0 or D7) are plotted against the increasing mean ranks. Sur la figure: …D0-D7 uDEGs, FC vs Rank
Supplemental Figure 8. ChEA3 analysis of dDEGs during the D0-D7 step. The D0-D7 |log_2_ FC| values of the 1,064 TFRs expressed (arbitrarily defined as displaying a mean expression of >5 on at least D0 or D7) are plotted against the increasing mean ranks. Sur la figure: …D0-D7 dDEGs, FC vs Rank Le nombre de TFRs (1064) est vraiment le même dans les deux cas ?

### Slide 14
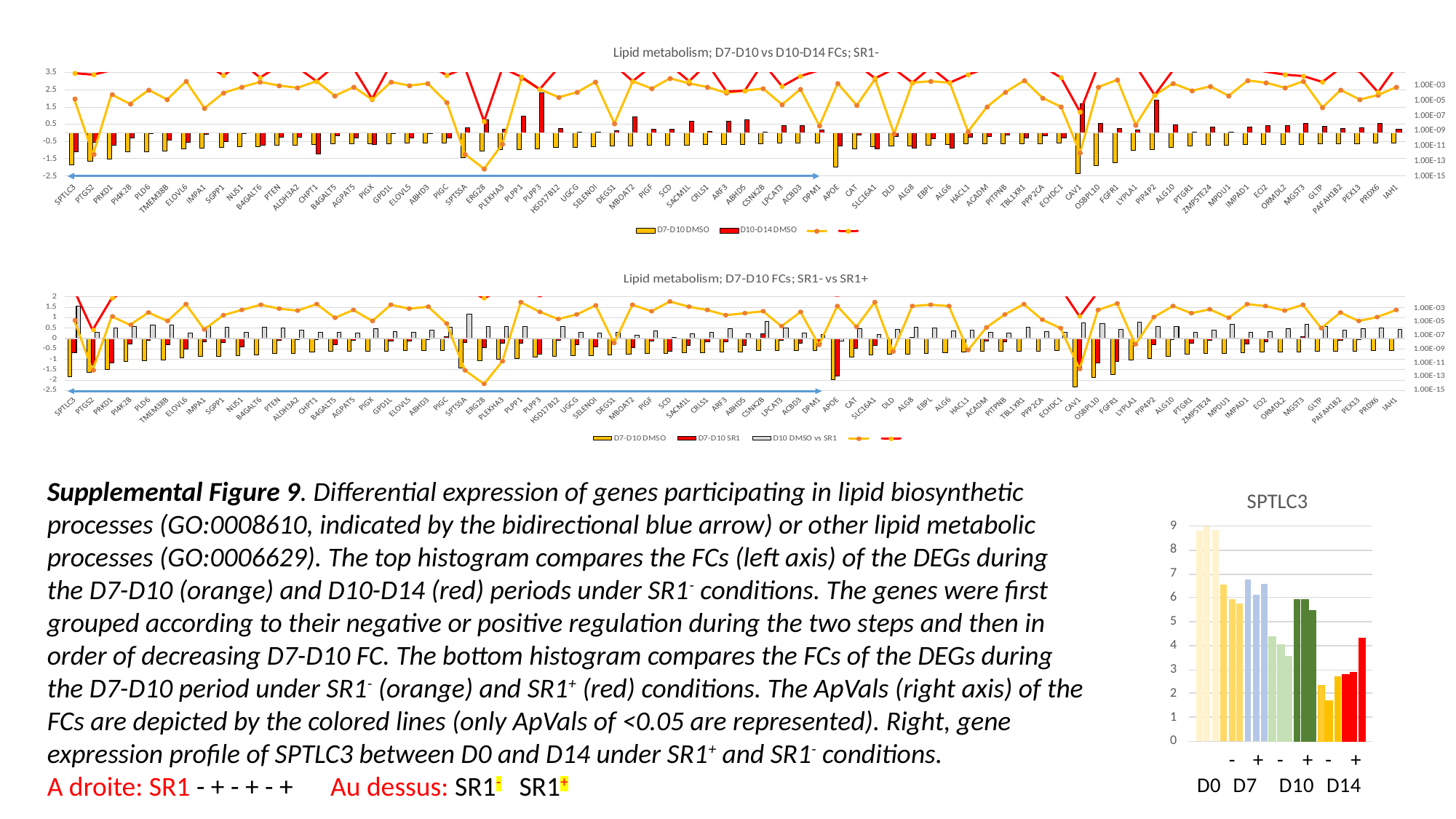

Supplemental Figure 9. Differential expression of genes participating in lipid biosynthetic processes (GO:0008610, indicated by the bidirectional blue arrow) or other lipid metabolic processes (GO:0006629). The top histogram compares the FCs (left axis) of the DEGs during the D7-D10 (orange) and D10-D14 (red) periods under SR1^-^ conditions. The genes were first grouped according to their negative or positive regulation during the two steps and then in order of decreasing D7-D10 FC. The bottom histogram compares the FCs of the DEGs during the D7-D10 period under SR1^-^ (orange) and SR1^+^ (red) conditions. The ApVals (right axis) of the FCs are depicted by the colored lines (only ApVals of <0.05 are represented). Right, gene expression profile of SPTLC3 between D0 and D14 under SR1^+^ and SR1^-^ conditions.
A droite: SR1 - + - + - + Au dessus: SR1^-^ SR1^+^

### Slide 15
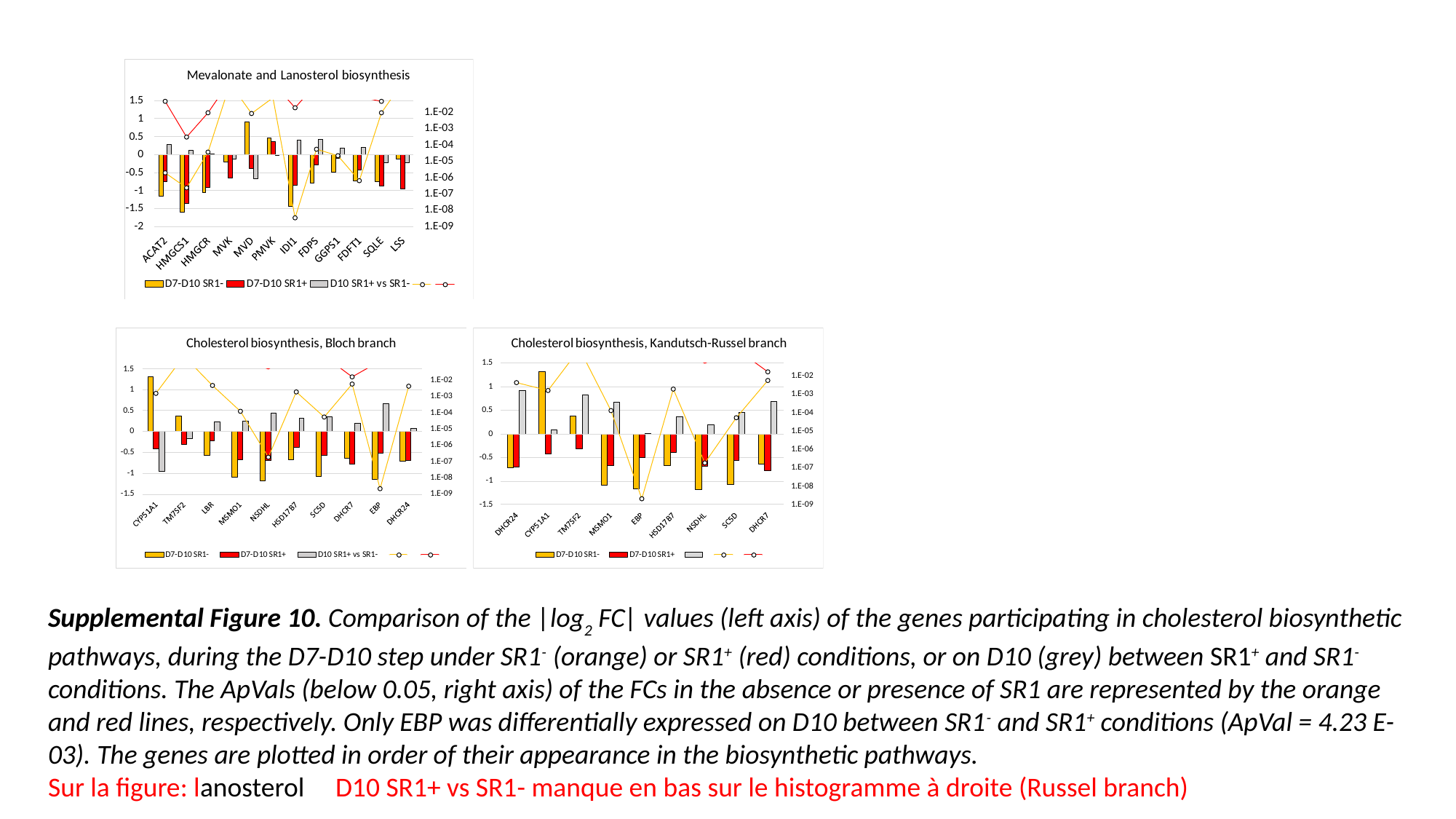

Supplemental Figure 10. Comparison of the |log_2_ FC| values (left axis) of the genes participating in cholesterol biosynthetic pathways, during the D7-D10 step under SR1^-^ (orange) or SR1^+^ (red) conditions, or on D10 (grey) between SR1^+^ and SR1^-^ conditions. The ApVals (below 0.05, right axis) of the FCs in the absence or presence of SR1 are represented by the orange and red lines, respectively. Only EBP was differentially expressed on D10 between SR1^-^ and SR1^+^ conditions (ApVal = 4.23 E-03). The genes are plotted in order of their appearance in the biosynthetic pathways.
Sur la figure: lanosterol D10 SR1+ vs SR1- manque en bas sur le histogramme à droite (Russel branch)

### Slide 16
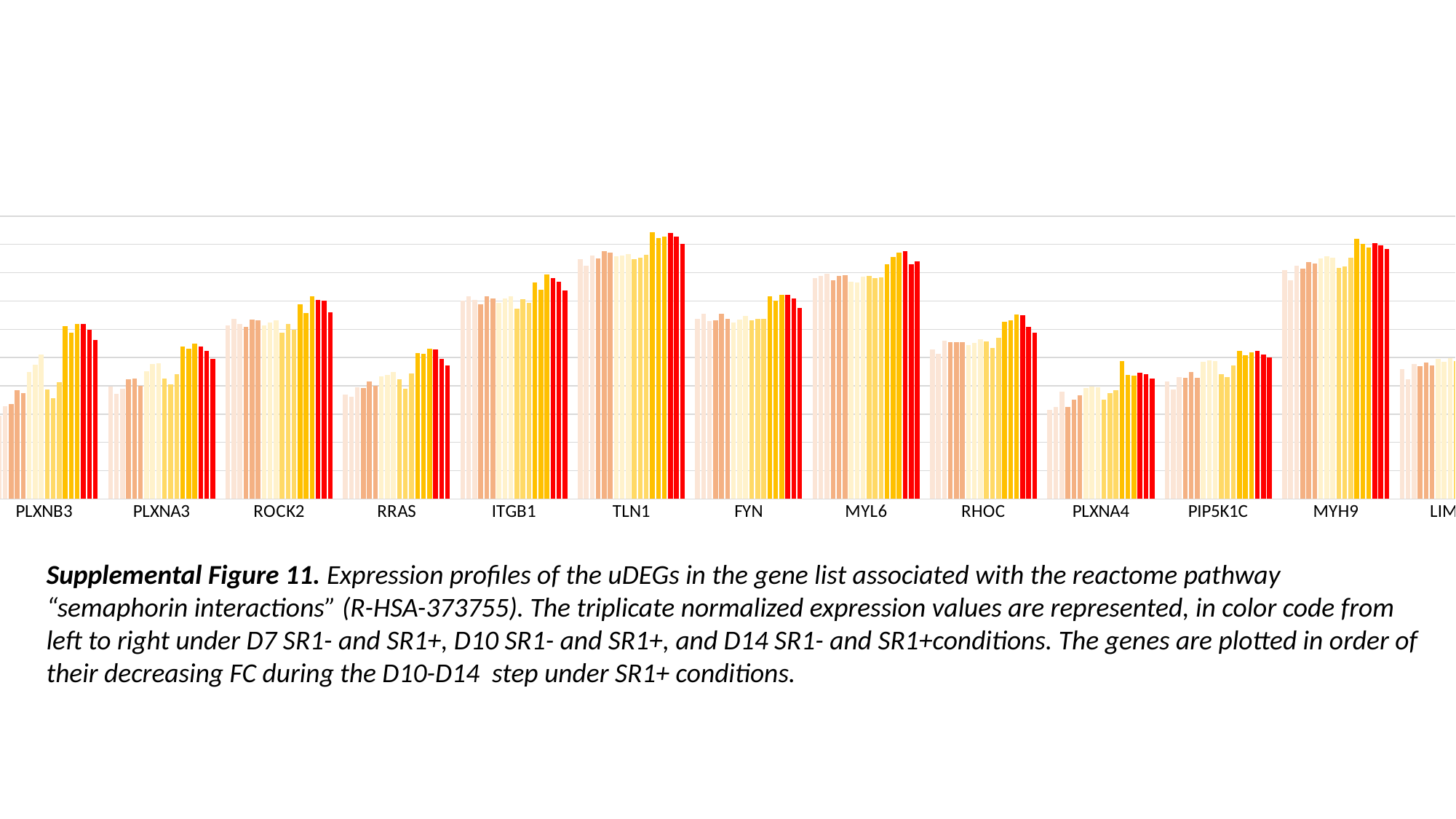

#### Chart
| Category | | | | | | | | | | | | | | | | | | |
|---|---|---|---|---|---|---|---|---|---|---|---|---|---|---|---|---|---|---|
| PLXNB3 | 6.714250517386525 | 5.866483355024694 | 6.5544733310164265 | 6.719789039017936 | 7.704308030813346 | 7.496528561927403 | 8.958678558661132 | 9.47089451968325 | 10.212576717963646 | 7.7308960878277055 | 7.136048535012794 | 8.240038156304102 | 12.245479262759478 | 11.750474678247654 | 12.393332190796384 | 12.356248500078395 | 11.954196732158604 | 11.23508284831445 |
| PLXNA3 | 7.946309642040518 | 7.404942998763008 | 7.808457844199195 | 8.436749578378379 | 8.50334503458993 | 7.980142315997474 | 9.016252871401752 | 9.557714324370503 | 9.593799519573386 | 8.50027729058697 | 8.101009996826948 | 8.808942276018682 | 10.793568401134054 | 10.617265545683336 | 10.96153483190157 | 10.788885533442157 | 10.462675286009556 | 9.905720706605019 |
| ROCK2 | 12.295117739962683 | 12.7455718276353 | 12.380383519208532 | 12.151357673004846 | 12.680651660529852 | 12.624669554996014 | 12.280507377616745 | 12.4736414384319 | 12.634079140798136 | 11.762140330589107 | 12.399972440251613 | 11.971050045912719 | 13.786838034290383 | 13.144396201041461 | 14.34262338334203 | 14.063550165116636 | 13.998608031399288 | 13.205227411815969 |
| RRAS | 7.360173519172375 | 7.2247481640079325 | 7.877548711537362 | 7.861478753260041 | 8.28656204606736 | 8.00254699135433 | 8.641215403000032 | 8.74352925714079 | 8.973341560250674 | 8.470050173677938 | 7.775214170623333 | 8.85123496758317 | 10.32344560133027 | 10.266663444717667 | 10.62667296876784 | 10.566843975205543 | 9.899392502259206 | 9.458123746960378 |
| ITGB1 | 14.006735660431776 | 14.35496097333657 | 14.068645957905542 | 13.754403634331645 | 14.312316433115509 | 14.199221286647303 | 13.887048272406359 | 14.161275131415895 | 14.325436753747839 | 13.439175008285043 | 14.117757354267486 | 13.845338837756994 | 15.319098102509535 | 14.796562741482049 | 15.874951808304576 | 15.632530824973806 | 15.36839546372852 | 14.768556939090448 |
| TLN1 | 16.95588586347509 | 16.5122757817238 | 17.227183062431074 | 17.024965080470995 | 17.51170558832795 | 17.400337301858173 | 17.171947405279976 | 17.21086710344592 | 17.30152530395527 | 16.96135192918856 | 17.075426839718688 | 17.24727205134404 | 18.883751046848076 | 18.425734427838567 | 18.566182693631504 | 18.8177595095264 | 18.54808057614137 | 18.01670707936193 |
| FYN | 12.717757367496874 | 13.075986906862408 | 12.59804153078106 | 12.619046301015922 | 13.087364707334121 | 12.712787184825919 | 12.496113110731743 | 12.699986170498176 | 12.956452398950095 | 12.623662397037332 | 12.752333438126227 | 12.758289524220274 | 14.331487083185154 | 14.008911361632098 | 14.451261177060315 | 14.441357662990528 | 14.178585294210425 | 13.490178367857881 |
| MYL6 | 15.60260828604577 | 15.775082114858954 | 15.926338561286471 | 15.473549544274636 | 15.788734099378884 | 15.841360966430793 | 15.369234880979317 | 15.323608366351301 | 15.73709388404567 | 15.748224984076682 | 15.620293111856702 | 15.66384248372919 | 16.574885547038345 | 17.113789049366552 | 17.437206571881035 | 17.50482485169493 | 16.596381019741905 | 16.780657111103494 |
| RHOC | 10.569933544134996 | 10.25201721044731 | 11.180323695862407 | 11.081569035919586 | 11.090990235803075 | 11.092001009104553 | 10.8690243431917 | 11.025171178542363 | 11.276039994816278 | 11.116896522075155 | 10.683116419753008 | 11.382622883988457 | 12.522595999194479 | 12.643811497234957 | 13.029517365103786 | 13.005850012379916 | 12.163390028964217 | 11.744362231528717 |
| PLXNA4 | 6.28671215692518 | 6.497731176255129 | 7.570818163337086 | 6.478093596563191 | 7.023711007978573 | 7.342644199460063 | 7.8188627302255975 | 7.935845709643037 | 7.896721344528397 | 7.023378231696851 | 7.48067371898372 | 7.702157275293282 | 9.733246347971477 | 8.781789476774371 | 8.71622671522298 | 8.923082933389418 | 8.806190004693367 | 8.51532215760712 |
| PIP5K1C | 8.326663935030998 | 7.730151545051195 | 8.63611686271189 | 8.58489203278367 | 8.974307631016938 | 8.564929615426184 | 9.691027186413816 | 9.8255921083873 | 9.761847343557136 | 8.79887288164371 | 8.598011126612823 | 9.458197845538768 | 10.458095761607051 | 10.185526841445272 | 10.342554720840822 | 10.460612108273944 | 10.216150002739305 | 10.029656858167527 |
| MYH9 | 16.17137709799093 | 15.456670652720895 | 16.46871697151915 | 16.289721548084366 | 16.75837108175367 | 16.66871434755281 | 17.027746864971764 | 17.15809745445845 | 17.084580160513834 | 16.359833622235 | 16.457496309162767 | 17.042297191040944 | 18.416041699577015 | 18.012222764334144 | 17.80557426975371 | 18.070834545878792 | 17.951790655903483 | 17.658283876932067 |
| LIMK1 | 9.164906056940099 | 8.445324985581058 | 9.527546094415902 | 9.381998679487582 | 9.666826463287935 | 9.426805518641734 | 9.91275303743926 | 9.706957438403803 | 9.94161780724719 | 9.749263471200562 | 9.347877637697525 | 9.941665893733864 | 10.850715673304968 | 10.761312005423427 | 10.915798454263006 | 11.191625711216927 | 10.754092971349223 | 10.56317010380358 |Supplemental Figure 11. Expression profiles of the uDEGs in the gene list associated with the reactome pathway “semaphorin interactions” (R-HSA-373755). The triplicate normalized expression values are represented, in color code from left to right under D7 SR1- and SR1+, D10 SR1- and SR1+, and D14 SR1- and SR1+conditions. The genes are plotted in order of their decreasing FC during the D10-D14 step under SR1+ conditions.

### Slide 17
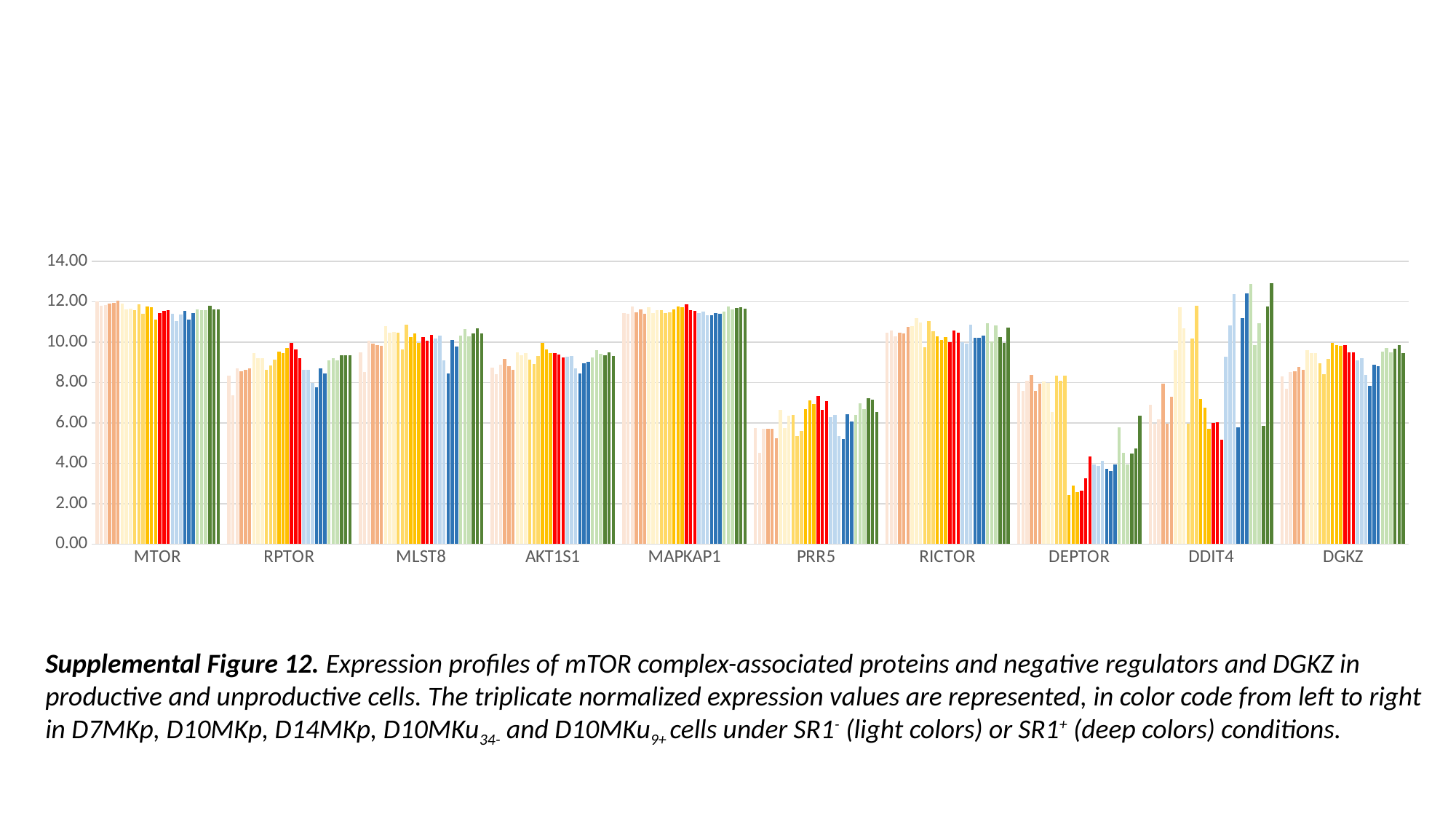

#### Chart
| Category | | | | | | | | | | | | | | | | | | | | | | | | | | | | | | |
|---|---|---|---|---|---|---|---|---|---|---|---|---|---|---|---|---|---|---|---|---|---|---|---|---|---|---|---|---|---|---|
| MTOR | 12.016168932526496 | 11.80003132840276 | 11.828100917300691 | 11.92820957182363 | 11.935479917533776 | 12.037877484058102 | 11.90852464402226 | 11.63239471057597 | 11.66012574027988 | 11.573393090116582 | 11.863101575050996 | 11.420052762253828 | 11.757017629243395 | 11.734963687732213 | 11.125058279847401 | 11.447767016461563 | 11.566851447213551 | 11.582862844965423 | 11.402256839809121 | 11.056769367566655 | 11.35429771238184 | 11.54390072552676 | 11.134196093632264 | 11.431385582784179 | 11.60615800643157 | 11.568791594083839 | 11.568530198902765 | 11.789946353602085 | 11.622240055846262 | 11.606550774638492 |
| RPTOR | 8.342934130201597 | 7.3708959774184715 | 8.712446769554084 | 8.549894695810375 | 8.640520221385001 | 8.706391231776667 | 9.463621551151277 | 9.225629408756655 | 9.194950008277399 | 8.644483659747424 | 8.853658129707863 | 9.120511417785346 | 9.538963245387173 | 9.463684291623847 | 9.71385954753285 | 9.955074493219321 | 9.625501422898807 | 9.207814735715026 | 8.641067922652105 | 8.648391916059902 | 7.991403030669094 | 7.772867595950475 | 8.70561110605118 | 8.456138402318768 | 9.091933365563783 | 9.216638108024705 | 9.11139338150977 | 9.352863129395084 | 9.357539005689151 | 9.346023104664582 |
| MLST8 | 9.482979605455968 | 8.510652439934127 | 10.058624810998747 | 9.928841859542432 | 9.865394229020831 | 9.831071452547453 | 10.795291558381155 | 10.484047288754732 | 10.499528129500757 | 10.48430817516983 | 9.626459556242192 | 10.860555973148887 | 10.250852684220813 | 10.436569829200502 | 9.964310750729235 | 10.264914553602917 | 10.055610426206028 | 10.359227468133803 | 10.178653817654352 | 10.337936137416532 | 9.107289775248544 | 8.450312408958485 | 10.106788166130366 | 9.789850867122036 | 10.317516368328066 | 10.640413009070725 | 10.291331991972795 | 10.446235301468697 | 10.676160921137077 | 10.44359892804093 |
| AKT1S1 | 8.750472294693823 | 8.41972990275637 | 8.894882740321082 | 9.184597622082029 | 8.8122098833897 | 8.636690123899616 | 9.497695553713418 | 9.339454058713146 | 9.462005030105617 | 9.132457611781154 | 8.902862695563126 | 9.324509240934368 | 9.962130128648747 | 9.628735614626272 | 9.447308027532696 | 9.467667479699907 | 9.40433459884422 | 9.243704154107661 | 9.267367853887388 | 9.325195693592388 | 8.721438773373617 | 8.433850200964892 | 8.946824628078591 | 9.02274613135227 | 9.248967272607667 | 9.609248250561137 | 9.409682209029206 | 9.356850394903269 | 9.500609100021116 | 9.300165322211942 |
| MAPKAP1 | 11.448092214430753 | 11.409956725122232 | 11.764464886078514 | 11.496146583067436 | 11.604109332807043 | 11.411970709411893 | 11.72116188248596 | 11.45655041584573 | 11.59494660016903 | 11.569785807733256 | 11.432073348878527 | 11.493892436585023 | 11.617502891135361 | 11.778553086879862 | 11.745821069730516 | 11.868158201335797 | 11.578260191347798 | 11.564560138826973 | 11.426206896878112 | 11.53029303768027 | 11.337407347887801 | 11.350919962563799 | 11.437302526183922 | 11.410057670930254 | 11.529505063757039 | 11.766306432583585 | 11.629596242120828 | 11.676388113184302 | 11.74600871870119 | 11.643591421722824 |
| PRR5 | 5.74584062876405 | 4.531055792203654 | 5.693897432043939 | 5.710961010808158 | 5.6950943068277375 | 5.2543537484272544 | 6.634618316953865 | 5.7354629172989915 | 6.349774565082823 | 6.377341382633795 | 5.353526358157406 | 5.610849008496769 | 6.695699534896655 | 7.132962659427913 | 6.928857568908121 | 7.3166028798788245 | 6.643637307735851 | 7.0964321433611275 | 6.275980056886098 | 6.398977214179542 | 5.34053265774555 | 5.200823583548955 | 6.420720596910589 | 6.055442113684726 | 6.400157622932895 | 6.968599879497383 | 6.675223016830139 | 7.2388143081992675 | 7.161747035051799 | 6.534722977653502 |
| RICTOR | 10.463098244698736 | 10.560557925415813 | 10.300421167822849 | 10.48537537930281 | 10.42749460329948 | 10.773588733156831 | 10.810693999032152 | 11.182713805214991 | 10.967453151392572 | 9.744512409331403 | 11.040845658498126 | 10.545590933030246 | 10.297622035853367 | 10.117503423766403 | 10.262588814259184 | 9.988996311432972 | 10.559514745203408 | 10.455248928773962 | 9.99394045518796 | 9.921796614147853 | 10.84960513076087 | 10.199387438021226 | 10.227673684567947 | 10.343015454376166 | 10.926120908738193 | 10.031438272001928 | 10.814716605992219 | 10.247787975182243 | 9.976829408862875 | 10.718753276365032 |
| DEPTOR | 7.964160174909521 | 7.584316092078634 | 8.07823380895734 | 8.36845975156448 | 7.569997566779476 | 7.9545733641681995 | 8.052467486359664 | 7.976461720697983 | 6.532808334165354 | 8.3535715918413 | 8.090506034655881 | 8.335808261186315 | 2.426544425223023 | 2.8915525402371878 | 2.5601667856835837 | 2.649688262358191 | 3.2432376647839694 | 4.327577137339273 | 3.9506818691749377 | 3.8874239913664206 | 4.118559484801994 | 3.7151766539785624 | 3.601985864579288 | 3.9601809076563543 | 5.795399481474828 | 4.506799691881363 | 3.94828076041834 | 4.479333571942557 | 4.731936961680209 | 6.375742061977135 |
| DDIT4 | 6.903217588049518 | 5.980231180192099 | 6.174846344157251 | 7.949901620877124 | 5.97995603403815 | 7.304173259369123 | 9.617912649323365 | 11.730435840944288 | 10.682802684602096 | 5.970287281534981 | 10.180330331267154 | 11.791024015091054 | 7.197300281086169 | 6.765015444264465 | 5.724795530827535 | 6.0007301317330235 | 6.027932711604993 | 5.162603859710441 | 9.276920347549195 | 10.832602923642918 | 12.387790647111586 | 5.776711906158411 | 11.199864912260113 | 12.42689399633952 | 12.878994245572676 | 9.86361619888283 | 10.955003127968798 | 5.862796919009767 | 11.780229096811873 | 12.903802929393647 |
| DGKZ | 8.316132368373918 | 7.683043196596665 | 8.528927883699849 | 8.553884303788882 | 8.772763943566371 | 8.646121518339381 | 9.58824360348326 | 9.473844978116812 | 9.45283241173926 | 8.972790442426405 | 8.43124539295603 | 9.160881268629442 | 9.96702304587145 | 9.865771828610463 | 9.811663745133453 | 9.863092050635984 | 9.512482472384004 | 9.512569136254621 | 9.0827016387012 | 9.203681312841272 | 8.383129760351332 | 7.8404419334269315 | 8.870937471275786 | 8.797579928290995 | 9.531651778652567 | 9.706572333121773 | 9.504498028508923 | 9.677213928233952 | 9.842354476721262 | 9.455997606507937 |Supplemental Figure 12. Expression profiles of mTOR complex-associated proteins and negative regulators and DGKZ in productive and unproductive cells. The triplicate normalized expression values are represented, in color code from left to right in D7MKp, D10MKp, D14MKp, D10MKu_34-_ and D10MKu_9+_ cells under SR1^-^ (light colors) or SR1^+^ (deep colors) conditions.

### Slide 18
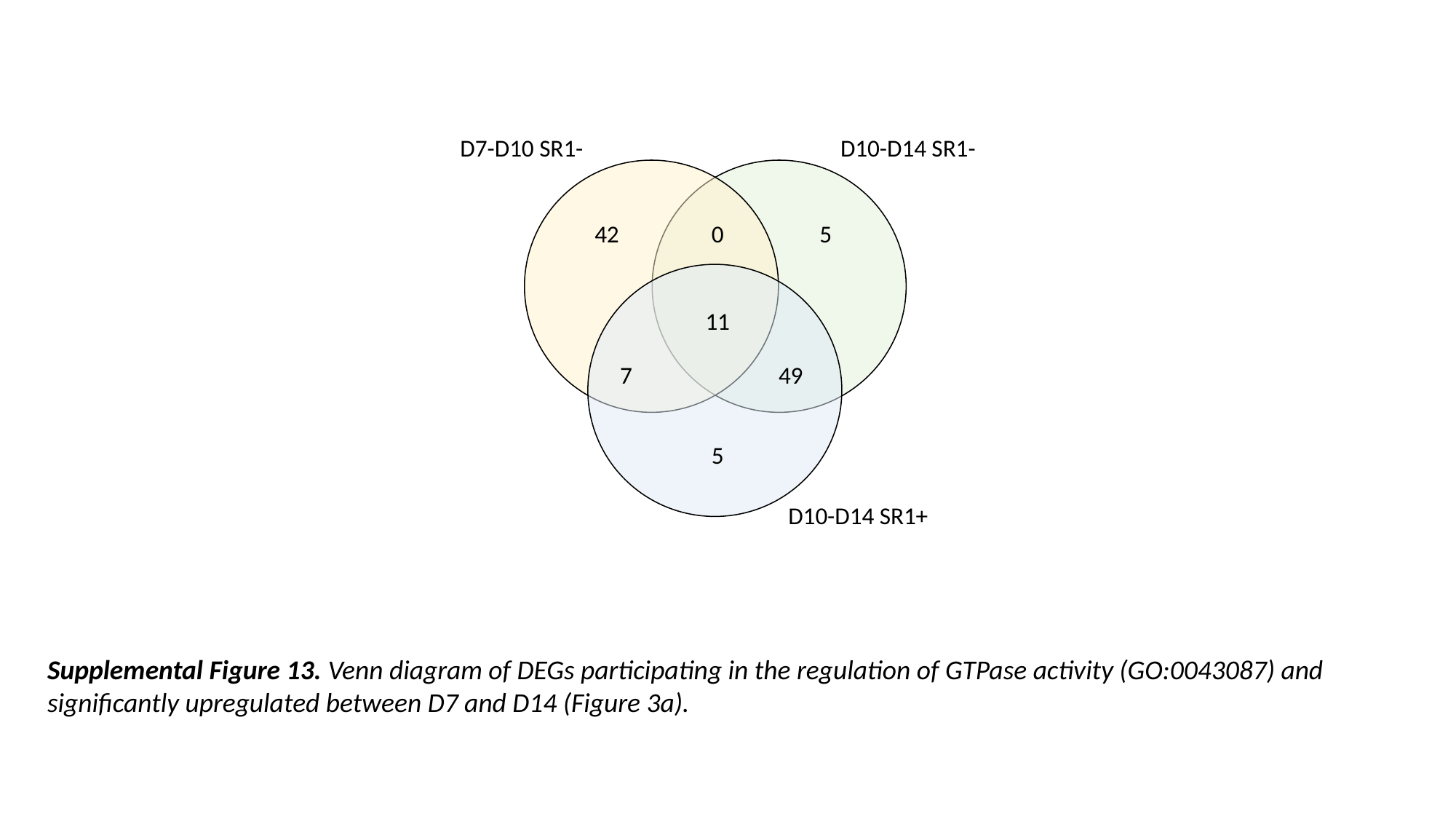

D10-D14 SR1-
D7-D10 SR1-
42
0
5
11
7
49
5
D10-D14 SR1+
Supplemental Figure 13. Venn diagram of DEGs participating in the regulation of GTPase activity (GO:0043087) and significantly upregulated between D7 and D14 (Figure 3a).
